# Genome-wide analysis of MIKC-type MADS-box genes in wheat: pervasive duplications may have facilitated adaptation to different environmental conditions

**DOI:** 10.1101/585232

**Authors:** Susanne Schilling, Alice Kennedy, Sirui Pan, Lars S. Jermiin, Rainer Melzer

## Abstract

**Background:** Wheat (*Triticum aestivum*) is one of the most important crops worldwide. Given a growing global population coupled with increasingly challenging climate and cultivation conditions, facilitating wheat breeding by fine-tuning important traits such as stress resistance, yield and plant architecture is of great importance. Since they are involved in virtually all aspects of plant development and stress responses, prime candidates for improving these traits are MIKC-type (type II) MADS-box genes.

**Results:** We present a detailed overview of number, phylogeny, and expression of 201 wheat MIKC-type MADS-box genes, which can be assigned to 15 subfamilies. Homoeolog retention is significantly above the average genome-wide retention rate for wheat genes, indicating that many MIKC-type homoeologs are functionally important and not redundant. Gene expression is generally in agreement with the expected subfamily-specific expression pattern, indicating broad conservation of function of MIKC-type genes during wheat evolution.

We find the extensive expansion of some MIKC-type subfamilies to be correlated with their chromosomal location and propose a link between MADS-box gene duplications and the adaptability of wheat. A number of MIKC-type genes encode for truncated proteins that lack either the DNA-binding or protein-protein interaction domain and occasionally show novel expression patterns, possibly pointing towards neofunctionalization.

**Conclusions:** Conserved and neofunctionalized MIKC-type genes may have played an important role in the adaptation of wheat to a diversity of conditions, hence contributing to its importance as a global staple food. Therefore, we propose that MIKC-type MADS-box genes are especially well suited for targeted breeding approaches and phenotypic fine tuning.

## Background

Bread wheat (*Triticum aestivum*) is one of the most important crops worldwide, contributing a significant amount of calories and proteins to the global human diet [1, 2]. Bread wheat is hexaploid and was first domesticated some 8 - 25,000 years ago in the region that today is called the middle-east [3, 4]. Wheat originated from three diploid progenitor species: *Triticum urartu* (A-genome donor), an *Aegilops speltoides*-related grass (B-genome donor) and *Aegilops tauschii* (D-genome donor) [5]. Because of its hexaploidy and an abundance of repetitive and transposable elements, bread wheat has one of the largest crop plant genomes (approximately 16 Gbp), making it challenging to work with from a genetics, genomics and breeding perspective [6]. However, recent advances in sequencing technology have led to a high-quality genome assembly and annotation by the International Wheat Genome Sequencing Consortium (IWGSC) [7]. Further, large scale RNA-seq analyses provided insights into expression patterns of homoeologous genes in different development stages and under a variety of stress conditions; building a rich resource for more detailed analyses [8].

Transcription factors (TFs) are a major driver in evolution as well as in domestication and bear the potential for crop improvement and trait fine-tuning [9]. MADS-box genes constitute one of the largest families of plant TFs [10]. They can be divided into two phylogenetically distinct groups: type I and type II [11]. While the function of most type I MADS-box genes remains to be illuminated, several type II genes are key domestication genes in different eudicot and monocot crops (reviewed in [12]). Plant type II MADS-domain proteins possess a typical domain structure, which is composed of the MADS-, I-, K- and C-terminal domain [13]. The MADS-domain enables the DNA-binding, nuclear localization and dimerization of the TF [14, 15], while I- and K-domain facilitate dimerization and higher-order complex formation of two or more MADS-domain proteins [16, 17]. The C-terminal domain allows for transcriptional activation of some MADS-domain proteins [18]. Because of this characteristic domain structure, type II genes are also referred to as MIKC-type MADS-box genes [13]. MIKC-type MADS-box genes are involved in virtually all aspects of plant development, including the root, flower, seed and embryo [19]. They have also been reported to be involved in different stress responses [20–22]. Thus, understanding MIKC-type MADS-box genes is important for understanding plant development, which is in turn crucial for plant breeding and crop improvement [23].

MIKC-type MADS-box genes have been phylogenetically and functionally characterized in a variety of model systems (Arabidopsis (*Arabidopsis thaliana*) and *Brachypodium distachyon* [24, 25]) as well as important crop plants (banana, rice, brassica, cotton [21, 26–28]). Individual wheat MIKC-type MADS-box genes have also been studied for almost two decades. The most prominent example is probably *VERNALIZATION1* (*VRN1*), an *APETALA1* (*AP1*)-like key regulator of flowering time as well as floral meristem determination [29–33] and one of the most important loci that distinguishes spring from winter wheat varieties [33]. Other wheat MADS-box genes have been implicated in the control of flowering time, ovule development and pistilloidy [34–37]. However, a detailed genome-wide phylogenetic and functional characterization of wheat MIKC-type MADS-box genes is still missing.

To better understand the dynamics of MIKC-type gene evolution in wheat and to facilitate future research on this important TF family, we provide a detailed overview of the number, phylogeny and expression of MIKC-type MADS-box genes in the recently released genome of *Triticum aestivum* [7]. We find a number of wheat MIKC-type subfamilies to be significantly larger than expected and suggest that extensive sub- and neofunctionalization in those subfamilies contributed to the global distribution of wheat.

## Results

### The wheat genome contains 201 MIKC-type MADS-box genes

A total of 439 coding sequences (including splice variants) were identified on the basis of the functional annotation (PFAM domains) in the recently released IWGSC wheat genome (Table S1) [7]. This dataset was simplified by keeping only one splice variant from each genomic locus for further analyses (Table S2). MIKC-type MADS-box genes were differentiated from type I MADS-box genes using a phylogenetic approach (Table S1 see Methods for details). An additional eight MIKC-type genes were identified using BLAST search. Altogether, we identified 201 MIKC-type (type II) MADS-box genes in wheat (Table 1, Table S2). Because MIKC-type MADS-box gene nomenclature in wheat is currently not consistent, with several genes having several synonymous names, we renamed all genes according to their subfamily association (Table 1, Table S2, see Methods for details).

**Table 1.**
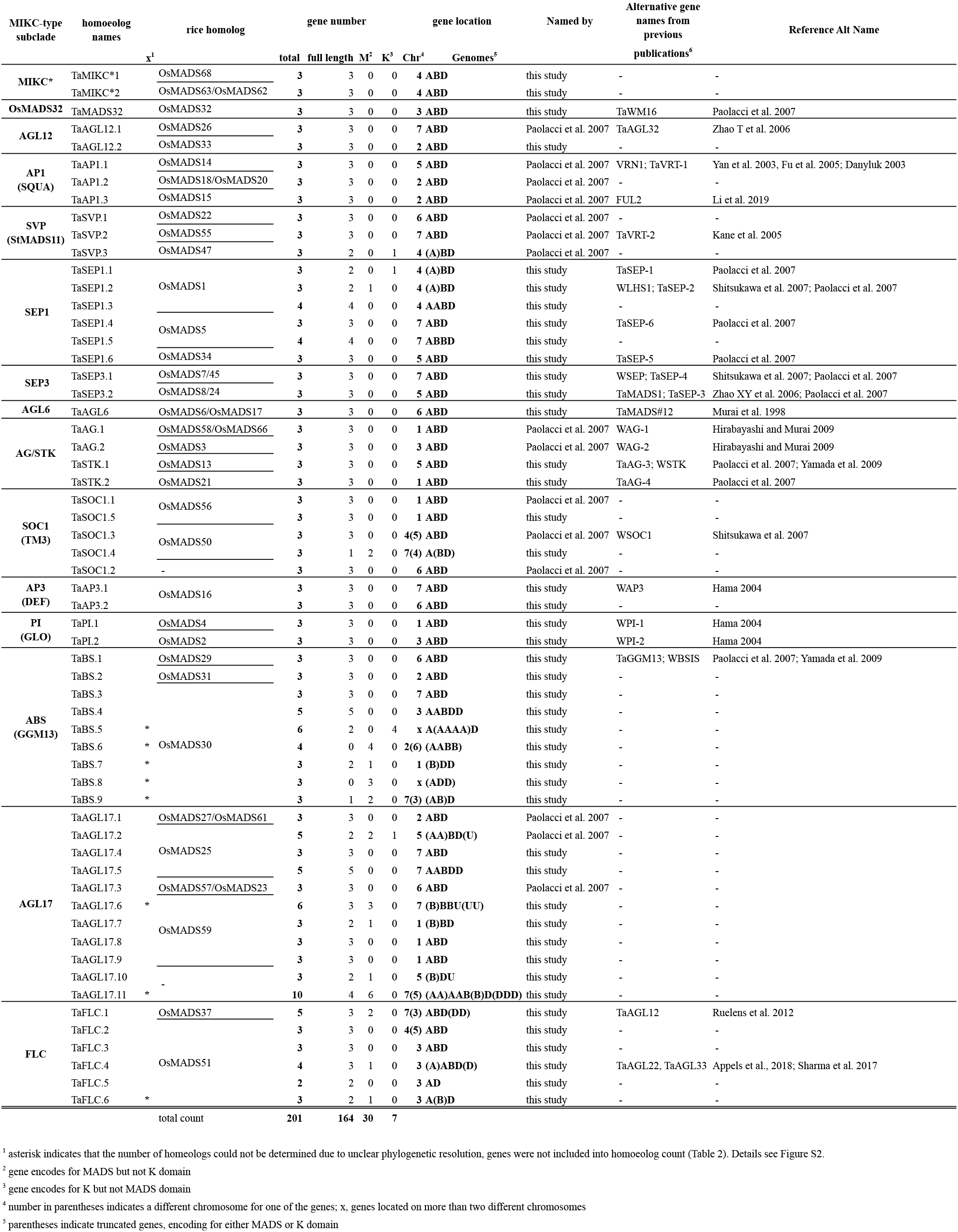
Subfamilies, names and numbers of wheat MIKC-type MADS-box genes. A complete list of all wheat MIKC-type genes can be found in Table S2.

The domain structure of MIKC-type MADS-domain proteins is crucial for their function, MADS- and K-domain being indispensable for DNA-binding and protein complex formation, respectively [13], although genes encoding truncated proteins may act as dominant negative versions [38, 39]. A total of 164 out of 201 MIKC-type genes encoded for both, a MADS- as well as a K-domain, while 29 genes lacked a K-box (14 %) and 7 lacked a MADS-box (3 %) (Table 1, Table S2, Figure 1). One gene encoded for a MADS- and an SRP54–domain (*TaBS.8A*, *TraesCS4A01G044400LC*) (Figure S3B, Table S2).

**Figure 1.**
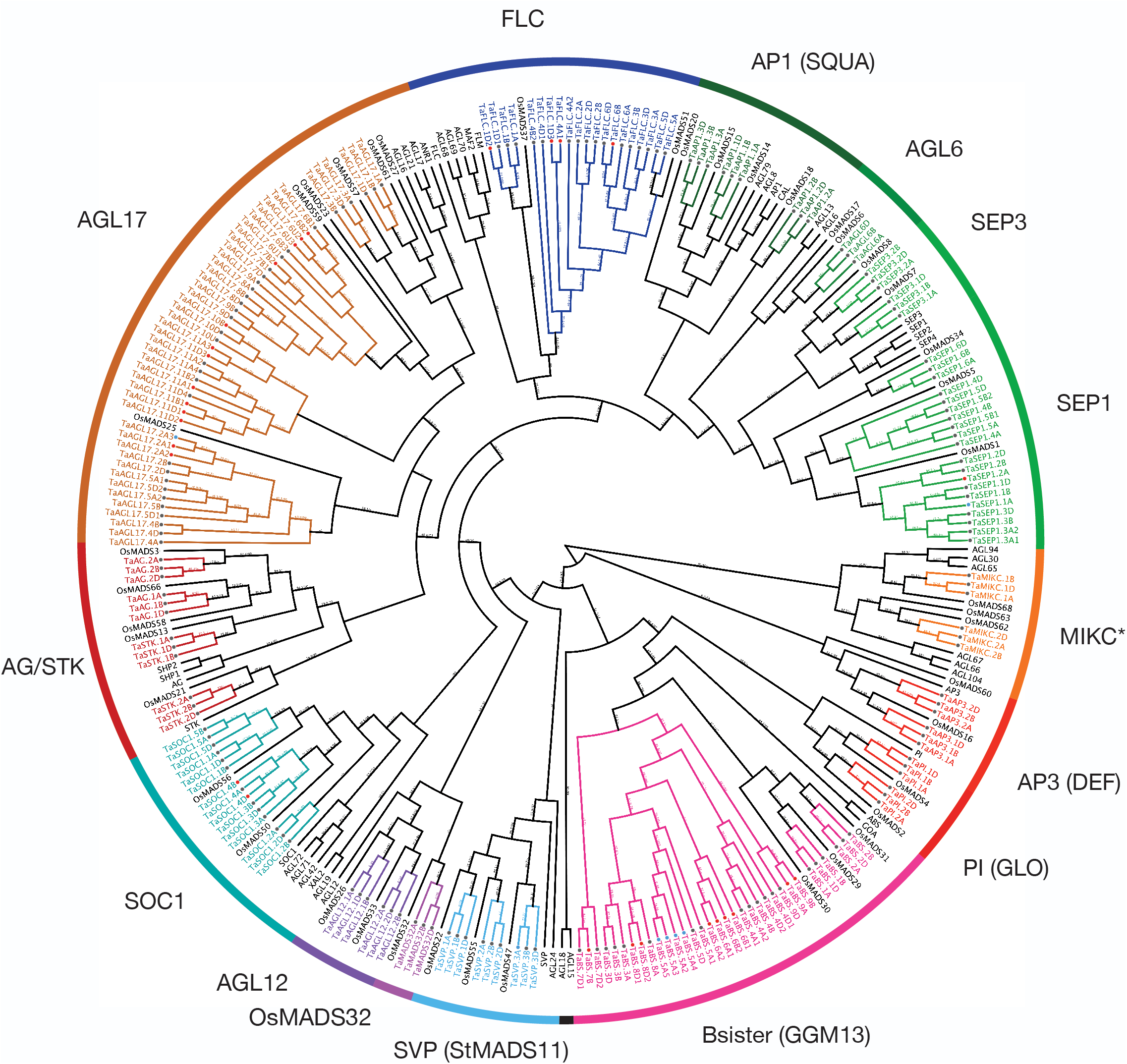
Maximum likelihood phylogeny of MIKC-type MADS-domain proteins from bread wheat, rice and Arabidopsis. A phylogenetic tree of MADS-domain proteins from bread wheat, cultivated rice and Arabidopsis was inferred using IQ-TREE [78, 79]. Wheat genes are coloured, rice and Arabidopsis genes are in black. The Arabidopsis genes *AGL15* and *AGL18* were included in the phylogeny; however, the *AGL15*-subfamily is lost in the grass lineage [40], therefore no rice or wheat gene clustered with them. *OsMADS32*-like genes are monocot specific [40], hence in this clade there is no Arabidopsis gene. SH-aLRT and Ultrafast bootstrap values are indicated on the branches in %, values equal to 100 are designated by a -. Accession numbers of all wheat genes can be found in Table S2, a version of the tree with untransformed branches can be found in Figure S2. The tree is unrooted, the MIKC* subclade was set as the outgroup.

For a number of genes, the predicted gene structures were very long (30 kb and above, Table S2). We confirmed mRNA transcripts spanning over especially long introns (ca. 29 kbp) for two selected genes from the *FLC*-clade using RT-PCR and Sanger sequencing (Figure S1).

### MIKC-type MADS-box genes belong to well-defined subfamilies

A Maximum Likelihood phylogenetic tree of all MIKC-type MADS-box genes from *Arabidopsis thaliana*, rice (*Oryza sativa*) and wheat shows that the wheat genome retained all 15 major grass MIKC-type MADS-box gene subfamilies: *AP1* (*SQUAMOSA* (*SQUA*)), *AP3 (DEF (DEFICIENS))*, *PISTILLATA* (*PI; GLOBOSA* (*GLO*)), *AGAMOUS* (*AG*) /*SEEDSTICK* (*STK*), *AG-LIKE6* (*AGL6*), *AGL12*, *AGL17*, *BSISTER* (*BS*; *GNETUM GNEMON MADS13,* (*GGM13*)), *SUPPRESSOR OF CONSTANS1* (*SOC1*), *SHORT VEGETATIVE PHASE* (*SVP*; *StMADS11*), *MIKC*\*, *OsMADS32*, *FLOWERING LOCUS C* (*FLC*), *SEPALLATA1*(*SEP1*) and *SEP3* (Figure 1, Figure S2) [40].

In many subclades, the gene phylogeny roughly followed species phylogeny; with the *Arabidopsis* genes displaying a sister-group relationship to the grass genes; and one or more rice MADS-box genes closely related to a triad of three wheat homoeologs (e.g. the *SVP*-, *AGL12*-, *OsMADS32*-, *MIKC*\*, *AP1*- and *AG*-subclades (Figure 1)). The topology in other subclades is more complex, suggesting multiple duplication events, before and/or after polyploidization of wheat (e.g. *SOC1*- and *SEP1* subclades, Figure S3A). Especially *Bsister*-, *AGL17*- and *FLC*- subclades are significantly expanded in wheat compared to Arabidopsis and rice (Figure 1, Figure 2, Figure S3B-D).

**Figure 2.**
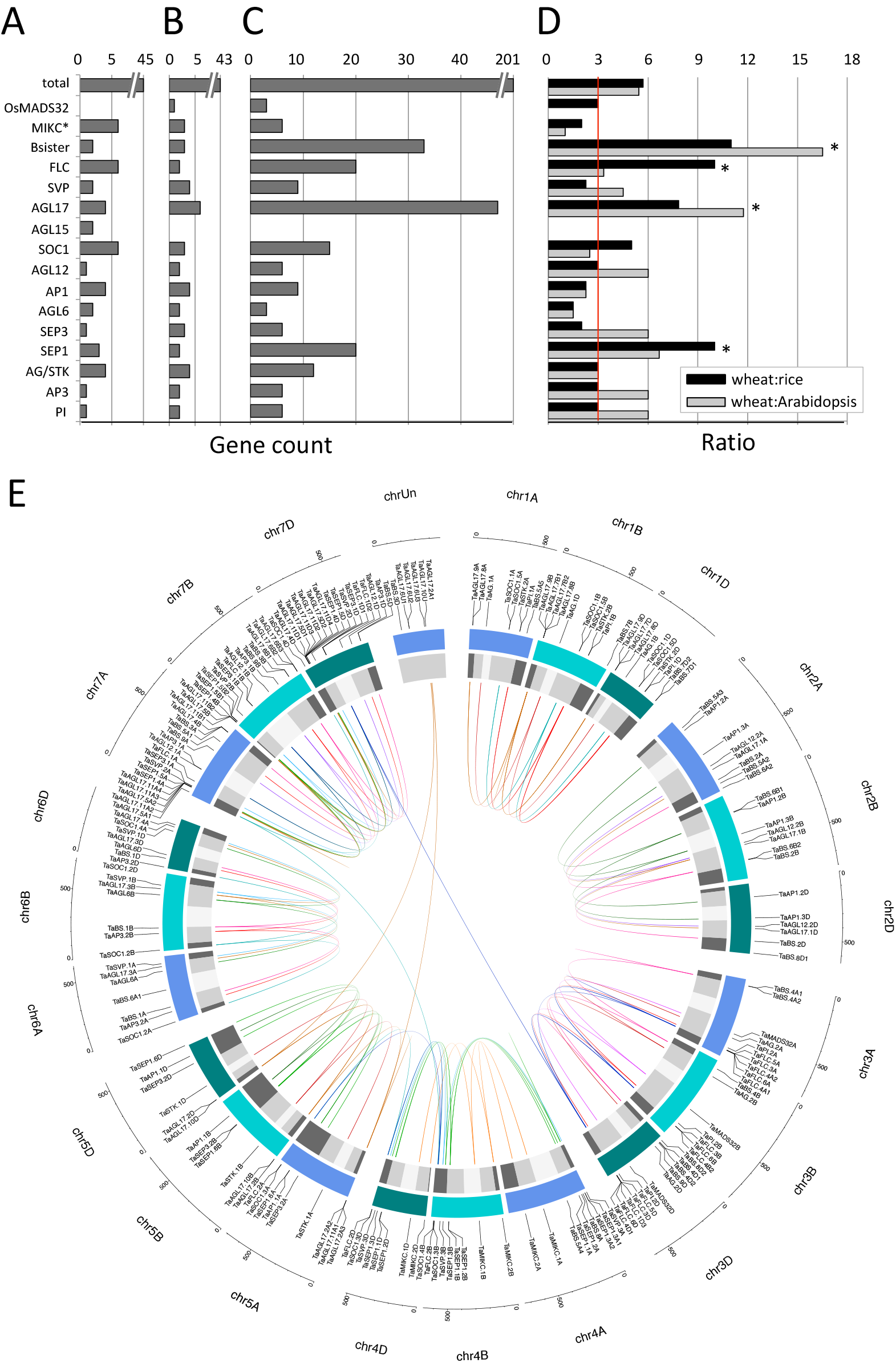
Number and location of MIKC-type genes. The number of MADS-box genes identified per MIKC-type subfamily in Arabidopsis (A), rice (B) and wheat (C) [21, 24]. The ratio of MIKC-type gene numbers total and in all subfamilies is shown for wheat to rice (black) and wheat to Arabidopsis (grey) (D). In (D) the expected ratio (3:1) is indicated by a red line, asterisks mark significant deviation from expected value (χ^2^ test, p < 0.05). All MIKC-type MADS-box genes were mapped to their respective locus in the wheat genome in a circular diagram using shinyCircos [92] (E). Subgenomes are indicated by different shades of blue (outer track), chromosomal segments are indicated by shades of grey (inner track) [7]. Homoeologous genes were inferred by mainly by phylogeny (details see Methods) and linked with subfamily-specific colors (inside of circle).

### Wheat MIKC-type genes exhibit a high rate of homoeolog retention and gene duplication

In many flowering plants, the number of MIKC-type MADS-box genes is between 40 and 70 [41]. Rice and Arabidopsis for example, despite their phylogenetic distance, have a similar number of MIKC-type MADS-box genes (43 and 45, respectively) [24, 42, 43]. With 201 genes, the total number of MIKC-type MADS-box genes in wheat is among the highest of hitherto characterized flowering plant species [27, 44, 45]. This is partly due to the hexaploid nature of wheat. However, even when corrected for ploidy-level, the number of MIKC-type MADS-box genes in bread wheat was significantly higher than in rice (≈1.5 fold higher, χ^2^ test p = 0.028, Figure 2A-D). Even when truncated genes lacking either a MADS-box or a K-box were excluded, the gene number was still higher than in rice (164/3=54.7 > 42). This increase in number is mainly due to the gene count in four subfamilies (*SEP1-, AGL17*-, *FLC*- and *Bsister*-like genes) that are significantly larger than expected (χ^2^ test p = 0.002, p << 0.001, p << 0.001, p << 0.001, respectively; Figure 2A-D). The remaining subclades showed the expected 3:1 ratio of wheat-to-rice genes; and some were below the expected ratio (Figure 2D). In some of the latter cases, wheat orthologs of rice genes could not be identified, indicating gene loss in the lineage leading to *Triticum* (e.g. *AP1-* and *AGL6-*like genes, Figure 1, Figure 2A-D).

To better understand why MIKC-type MADS-box genes are so abundant in the wheat genome, we analyzed homoeologous groups in detail (Table 2). Approximately one third of all wheat genes (i.e. all genes annotated in the current version of the wheat genome) are present in homoeologous groups of 3, also termed triads (1:1:1; 35.8 % of genes) [7]. In contrast, almost two thirds of the 201 MIKC-type genes identified are present in triads (62.7 %, Table 2). If only ‘full-length’ MADS-box genes (defined here as containing a MADS- and K-box) are considered, this ratio is even higher (72.7 %, Table 2). Also, the percentage of MIKC-type genes with homoeolog-specific duplications is higher for MIKC-type genes as compared to all wheat genes (8.5 % vs. 5.7 % Table 2). Loss of one homoeolog, on the other hand, is less pronounced in MIKC-type genes (1 % vs. 13.2 %, Table 2). 38 genes were excluded from the analysis (18.9 %; 13 % for full-length only), because the relationship of the genes could not be reliably resolved. Thus, the high homoeolog retention rate can partly explain the high number of wheat MIKC-type genes.

**Table 2.** Groups of homoeologous MIKC-type MADS-box genes in wheat.

| homoeologous group (A:B:D) | all wheat genes<sup>1</sup> | wheat MIKC MADS (all) |  |  | wheat MIKC MADS (full-length only) |  |  |
| --- | --- | --- | --- | --- | --- | --- | --- |
|  |  | # of groups | # of genes | % genes<sup>2</sup> | # of groups | # of genes | % of genes<sup>3</sup> |
| 1:1:1 | 35.8% | 42 | 126 | 62.7% | 40 | 120 | 73.2% |
| n:1:1/1:n:1/1:1:n | 5.7% | 4 | 17 | 8.5% | 2 | 8 | 4.9% |
| 1:1:0/1:0:1/0:1:1 | 13.2% | 1 | 2 | 1.0% | 6 | 12 | 7.3% |
| other ratios<sup>4</sup> | 8.0% | 4 | 18 | 9.0% | 3 | 10 | 6.1% |
| orphans/singletons | 37.1% | 0 | 0 | 0.0% | 1 | 1 | 0.6% |
| excluded<sup>5</sup> | - | - | 38 | 18.9% | - | 13 | 7.9% |
|  | 99.8% |  | 201 | 100.0% |  | 164 | 100.0% |

<sup>1</sup> according to Appels et al., 2018
<sup>2</sup> % calculated with 201 genes
<sup>3</sup> % calculated with 164 genes
<sup>4</sup> either n:1:n or 0:1:n
<sup>5</sup> see Table 1, Figure S2 and Methods

In addition to this, a variety of duplication patterns were observed in some subfamilies (Figure S3, Supplementary Text 1). For example, the rice *SEP1*-like genes *OsMADS1* and *OsMADS5* are sister to 10 and 7 wheat genes, respectively. In this case, the phylogenetic analysis suggests gene duplications in the lineage leading to *Triticum* but before the polyploidization of wheat (Figure S3A). The *Bsister* gene *OsMADS30* from rice is sister to 27 wheat genes, many of which lack a MADS- or K-box and are found in non-syntenic regions, indicating gene amplification occurred through transposable elements (Table 1, Figure S3B). For the *AGL17*- and *FLC*-subfamily from wheat, a number of triads can be assigned to a single rice gene (Figure S3C, D). Several genes from these two subfamilies are found in close vicinity to each other, pointing towards tandem duplications as a mechanism for subfamily expansion (Table 1, Figures 2E and S2C, S2D).

### Gene duplications and truncated genes are prevalent among MIKC-type genes in subtelomeric segments

MIKC-type MADS-box genes were generally equally distributed among the chromosomes, the only exception being the three homoeologous chromosomes 7, which contained significantly more genes than expected from the chromosome lengths (χ^2^ test p << 0.001, Figure 2E). This is mainly due to *AGL17*-like genes; with the majority of them located in tandem locations on the distal subtelomeric ends of chromosomes 7 (Figure 2E, Table S2). Overall, MIKC-type genes were located equally likely in the more central segments of the chromosomes (R2a, R2b and C) and in the subtelomeric parts of the chromosomes (R1 and R3) (48 and 50 % of genes, respectively). However, gene location varied greatly among subfamilies (Figure 3). In general, genes belonging to smaller subfamilies tended to be located in more central segments of the chromosomes; whereas a larger percentage of genes belonging to more expanded subfamilies were located in subtelomeric segments (Figure 4, Table S2). Further, full-length MIKC-type genes were about equally likely to be located within sub-telomeric versus central segments (45 % vs. 54 %, respectively), but truncated genes (MADS- or K-box only) were two times as prevalent in subtelomeric segments (61 % vs. 31 %; 8 % of genes were not assigned to a chromosome) (Figure 3A). Genes in subtelomeric segments were often found to be in close vicinity to each other, with over half of the genes in subtelomeric segment R1 being under 1000 kbp downstream of the closest MIKC-type gene (Table S3). In contrast, only 15 % of genes in the centromere segment have a distance under 1000 kb to the next downstream MIKC-type gene (Table S3). This might be due to more frequent tandem duplication events in subtelomeric segments. These findings are in line with the observation that subtelomeric segments are targets of recombination events and that many fast evolving genes lie within these segments [46].

**Figure 3.**
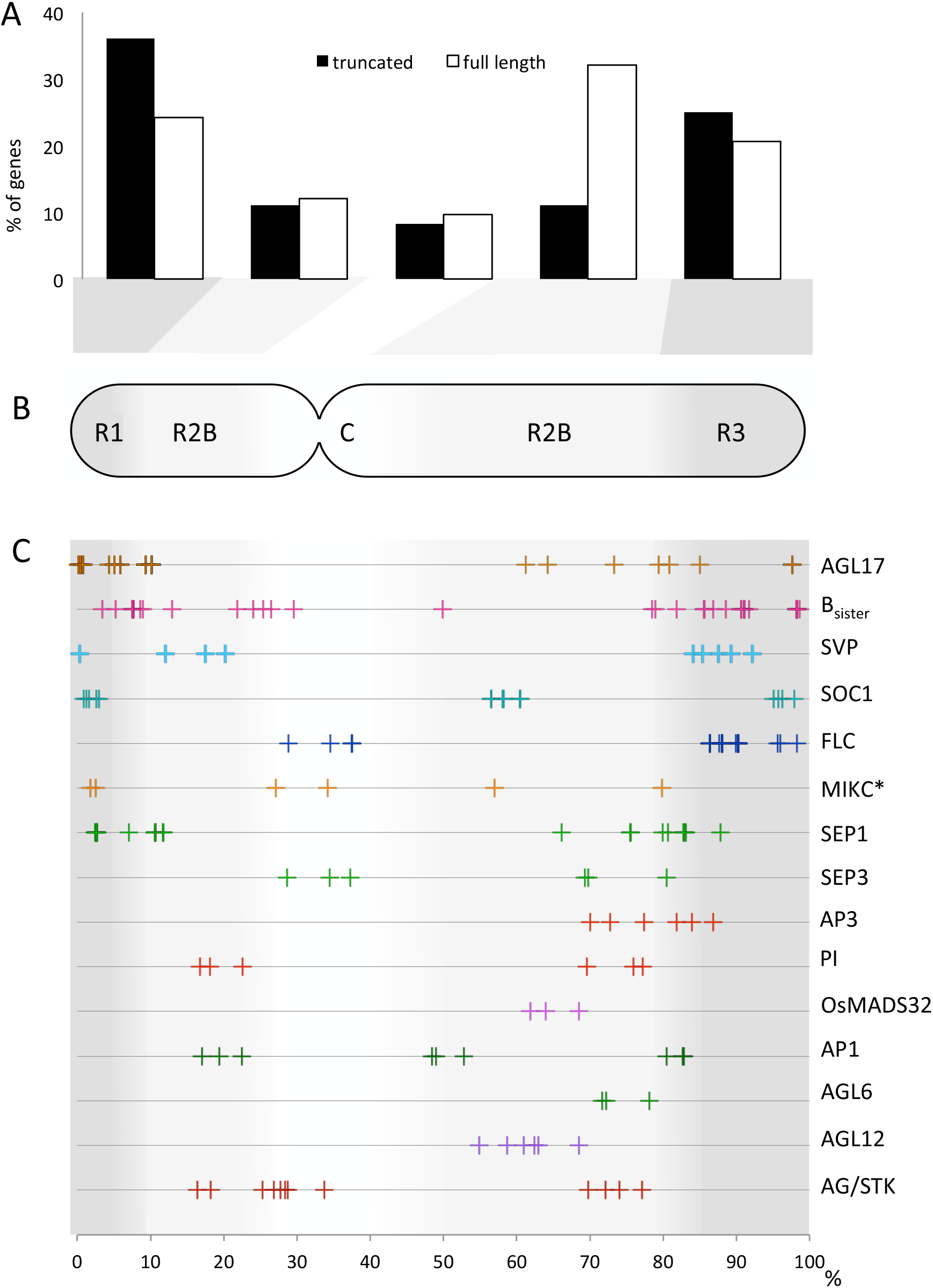
Chromosomal distribution of wheat MIKC-type MADS-box genes. The proportion of truncated (black) and full-length (white) MIKC-type genes in every chromosome segment is shown in a bar diagram (A). A schematic overview of the chromosome indicates the different segments, R1 and R3 (dark grey), R2A and R2B (light grey) and C (white) (B). The location of all genes belonging to each MIKC-type subfamily is shown as percentage of chromosome length (C). Segments have been averaged over all chromosomes, segment lengths according to [7].

**Figure 4.**
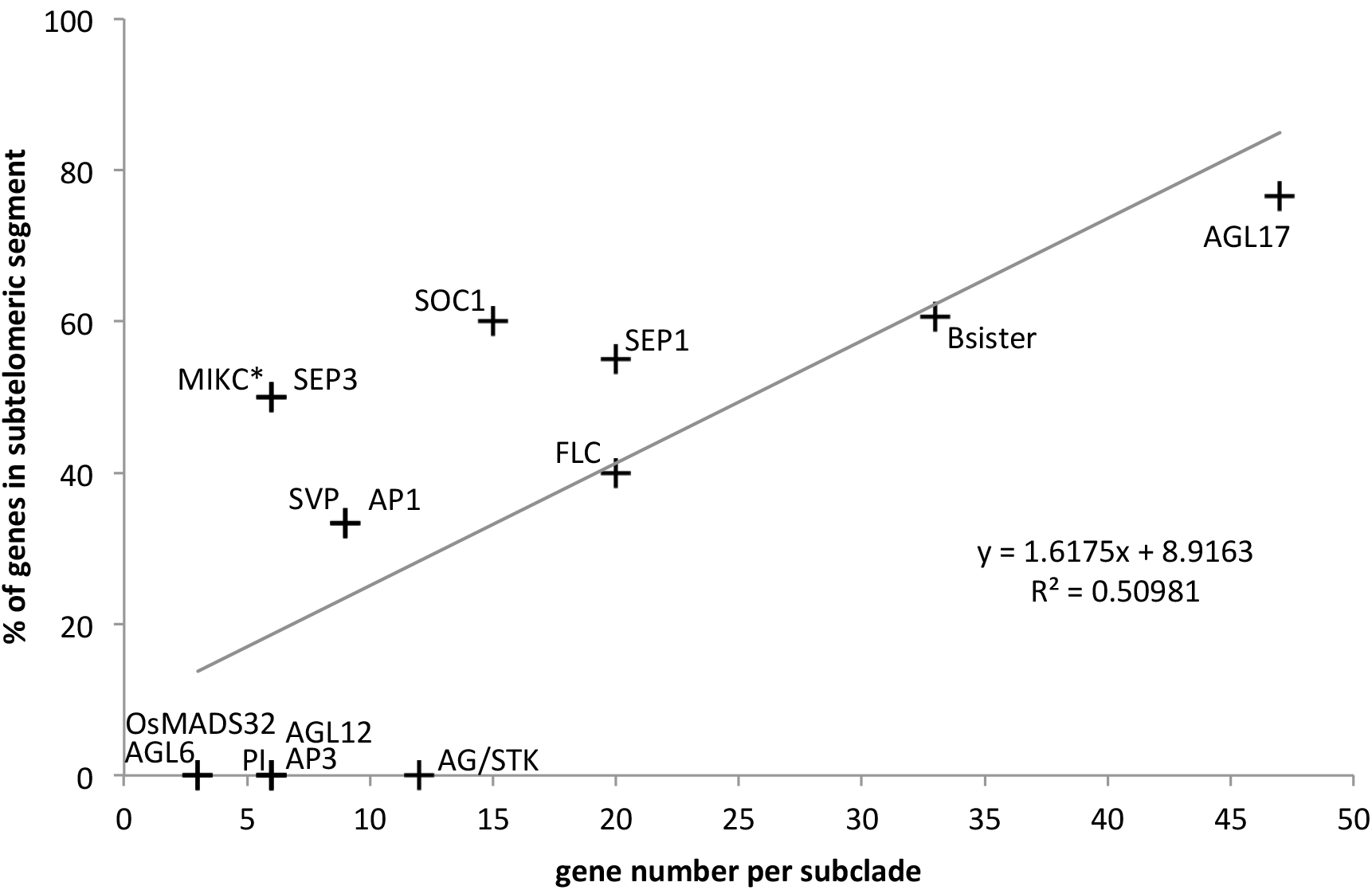
Distribution of MIKC-type genes to subtelomeric or central chromosome segments. The number of genes per MIKC-type subclade was plotted against the fraction of genes located in central chromosome segments (R2a, R2b and C) and subtelomeric segments (R1 and R3). Subclades are indicated next to data points. Data points for some subfamilies are identical (*AGL6* and *OsMADS32; SVP* and *AP1; PI, AP3* and *AGL12; SEP3* and *MIKC*\*).

### Conserved and divergent patterns of MIKC-type MADS-box gene expression during wheat development

To characterize the expression of wheat MIKC-type MADS-box genes, we analyzed RNA-seq data of 193 wheat MADS-box genes [8, 47]. Out of the 159 full-length genes, 83 % were expressed in at least one developmental stage, with a wide expression range with a maximum count of 1 to 424 transcripts per million (tpm_max_) (Figure 5, Table S2). The remaining 17 % of full-length genes showed a very low expression with a tpm_max_ below 1 (Figure 5A, Table S2). Of the 33 truncated genes, encoding only for either K or MADS-domain, 30 % were expressed (4 and 6 genes, respectively, tpm_max_1 to 51; Figure 5A, Table S2). One gene, encoding for MADS- and an SRP54-domain was expressed ubiquitously in the plant, albeit at a low level (*TaBS.8A*; tpm_max_3.12; Table S2, Figure S3).

**Figure 5.**
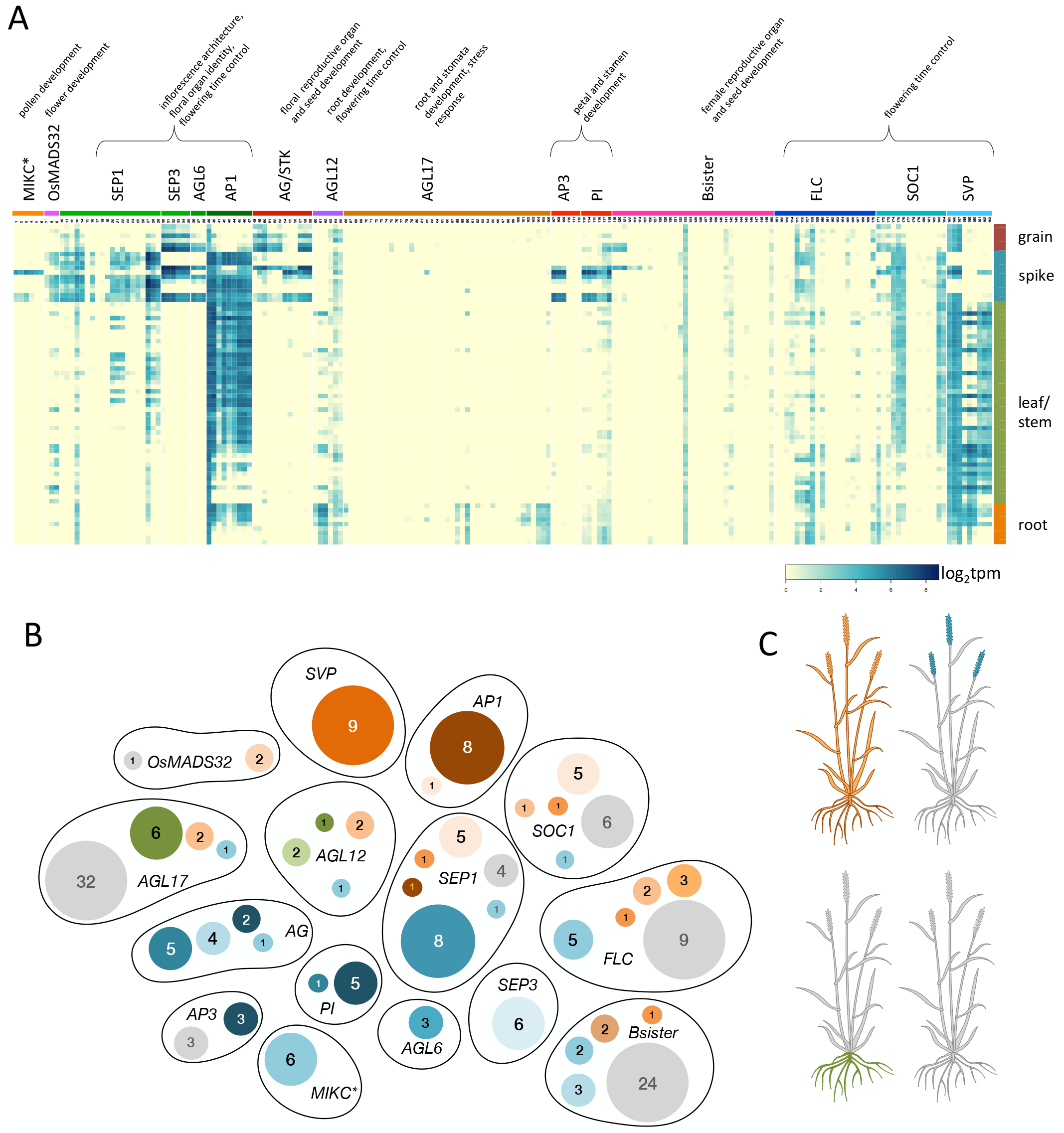
Expression of MIKC-type MADS-box genes during wheat development. Expression analysis was done for all subfamilies using RNA-seq data [8]. A heatmap shows expression level of all genes in different subfamilies (columns) and stages/tissues (rows) (A). Genes and tissues are listed in Tables S6 and S7, respectively. Genes were clustered into different modules according to their expression and mapped to subfamilies. Colors indicate different modules: shades of brown indicate ubiquitous expression, shades of green indicate expression in the root, shades of blue indicate expression in reproductive organs and grey indicates low or no expression during development (B). The digits inside the circles indicate the number of genes in the respective module. The clustering heatmap with all modules can be found in Figure S4. A schematic representation of a wheat plant depicts colors indicating different expression modules (C).

In general, MADS-box genes expression patterns are comparable with findings in rice [21, 48, 49] (Figure 5, Figure S4). MIKC*-type genes are expressed in anthers (Figure 5A, Table S7). Putative floral homeotic genes, such as *PI/AP3*-like and *AG/STK*-like genes, *OsMADS29*- and *OsMADS31*-like *Bsister* genes, *AGL6*-like genes as well as *SEP1* and *SEP3*-like genes are expressed specifically during flower and seed development (Figure 5). *AGL17*-like genes are expressed in roots. *AP1*-like, *SVP*-like and some *SEP1*-like genes are expressed ubiquitously throughout the plant life cycle in different tissues.

For further analysis, genes were hierarchically clustered according to expression similarities and then grouped into different expression modules (Figure 5B, Figure S4). This analysis showed that, genes from one subfamily could differ considerably in their expression pattern. For example, members of the *SEP1*-like gene subfamily are grouped into 6 different modules and *Bsister* genes are found in 5 different modules (Figure 5B). In contrast, *SVP-* and *AP1-* like genes show relatively little variation in their expression pattern (Figure 5B). It is also noteworthy that 79 genes including representatives from almost all subclades showed no expression or only low expression under very specific conditions during the developmental time course (41 % of genes; module 9; grey, Figure 5B).

### *AGL17*-like and *Bsister* genes are expressed in response to stress conditions

*AGL17-* and *OsMADS30*-like *Bsister* clades have been expanding during wheat evolution (Figure 1, Figure 2). Many of the genes from these clades are not expressed or expressed on a very low level during wheat development (Figure 5). However, some of these genes do show expression in response to distinct stress conditions (Figures 6, S4).

*Bsister* genes form three distinct clades of *OsMADS29*- *OsMADS30*- and *OsMADS31*-like genes in grasses [50]. Most *Bsister* genes have been described to be important for ovule and seed development with a specific expression pattern limited to female reproductive organs [51–53]. While *OsMADS29*-like and *OsMADS31*-like wheat *Bsister* genes follow this conserved expression pattern (Figure 6A, *TaBS.1* and *TaBS.2*), some *OsMADS30*-like wheat genes are expressed ubiquitously during the plant life cycle (Figure 6A, *TaBS.6B1 TaBS.5A4*). In contrast, five closely related *OsMADS30*-like genes showed low or no expression in any of the developmental stages. Instead, a specific upregulation in response to inoculation with the pathogen *Fusarium graminearum* was detected (Figure 6A, *TaBS.4*; Figure S5).

**Figure 6.**
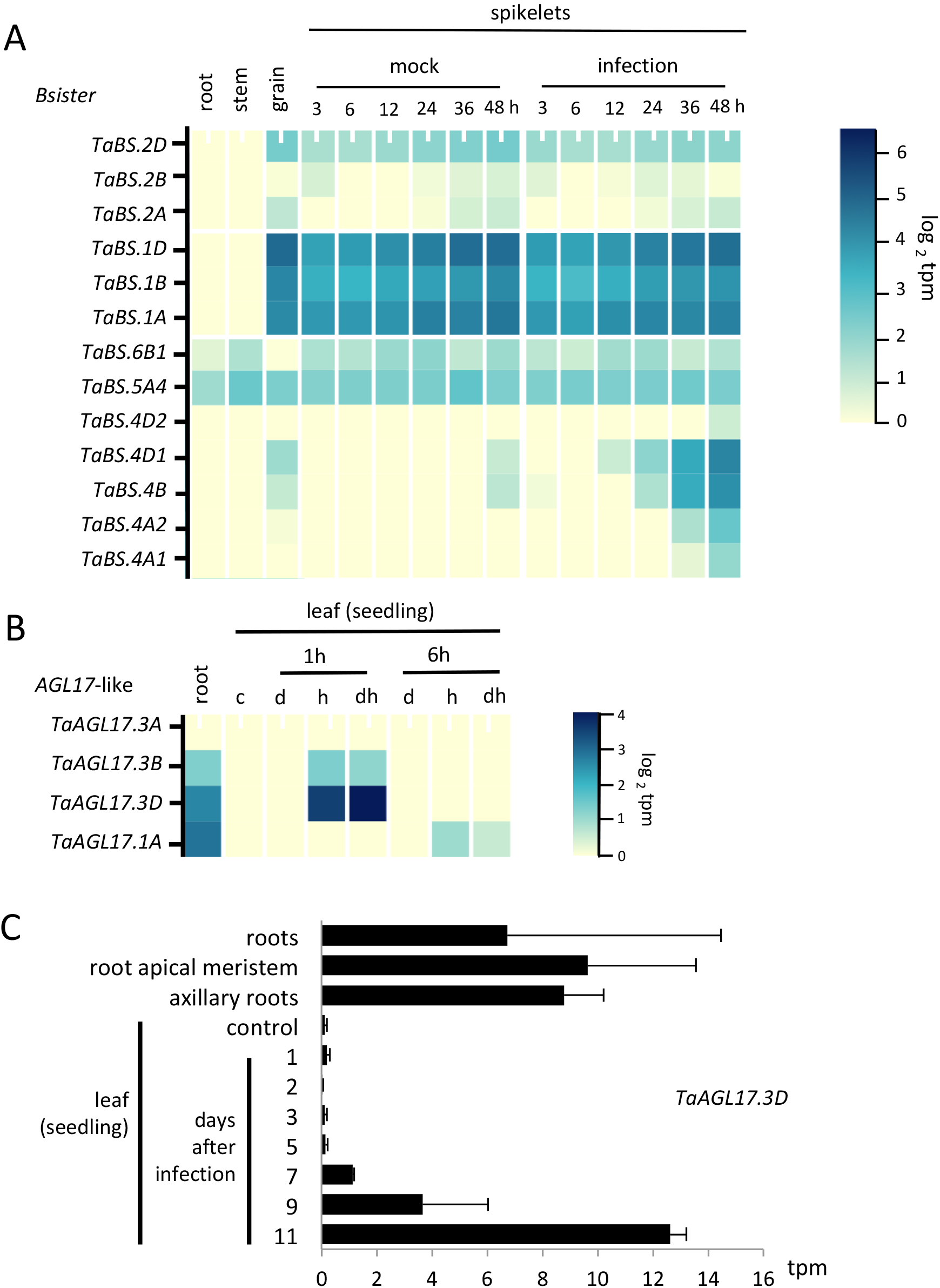
Expression of Bsister and AGL17-like genes in response to stress conditions. RNA-seq data was analyzed using wheatexpression.com [47]. *Bsister* gene expression in different tissues [8] and in response to infection with Fusarium head blight [93] (A). Expression of wheat *OsMADS31*-like genes (*TaBS.2*) *OsMADS29*-like genes (*TaBS.1*) and *OsMADS30*-like genes (*TaBS.4*, *TaBS.5A4*, *TaBS.6B1*) in root, stem and grain [8] as well as spikelets during mock inoculation and infection with Fusarium graminearum after different time points is shown as a heatmap. *AGL17*-like gene (1-4) expression in roots and seedling leaves under control conditions (c) and after 1 and 6 hours of drought (d), heat (h) and combined drought and heat stress (dh) [94] (B). Expression of *TaAGL17.3D* within different root tissues [8] and in leaves under control conditions and after infection with stripe rust (*Puccinia striiformis*) [95] is shown as a bar diagram. SEM is indicated as error bars. Data is shown as log2tpm (A, B) or as tpm (C).

*AGL17*-like genes are commonly expressed in roots and leaves [54]. They have also been described to be involved in osmotic and saline stress responses in rice [54]. Several wheat *AGL17*-like genes are not expressed at all during developmental stages (Figure 5, grey; Figure 6B). Two *AGL17*-like genes that are expressed in the root, but not in leaves under control conditions, show upregulation in leaf tissue after 1 hour of heat stress (Figure 6B, genes 2,3). However, gene expression is not detectable after 6 h and there is no specific response to drought stress (Figure 6B, genes 2,3). Another *AGL17*-like gene is upregulated after 6h of heat, but not drought stress (Figure 6B, gene 4). A fifth *AGL17*-like gene is expressed in the root, and additionally upregulated in leaves in response to infection with stripe rust 7 days after infection (Figure 6C).

## Discussion

### Many wheat MIKC-type MADS-box genes have evolutionarily conserved functions

MIKC-type MADS-box genes play a central role in plant development. They are therefore promising targets for crop breeding and improvement [55].

Here, we identified 201 wheat MIKC-type MADS-box genes, which we assigned to 15 conserved subfamilies (Table 1, Figure 1). At least 70 % of wheat MIKC-type MADS-box genes could be assigned to homoeologous groups with genes in every subgenome (Table 2). This is considerably above the average homoeologous retention rate in wheat (42 %, Table 2 [7]). Many MADS-domain proteins act in multi-protein complexes [56]. The composition of those protein complexes changes dynamically during development, with many proteins being part of more than one complex [57, 58]. Changes in the gene dosage may result in changes in the stoichiometry of the protein complexes which may in turn have detrimental phenotypic effects [59]. Thus, selection may act to retain homoeologs in all subgenomes.

We also found that the expression pattern of many wheat MADS-box genes is similar to that of close homologs in rice and other model plants, indicating that gene functions are broadly conserved between wheat and rice.

Together, the conservation of all major subclades, the high homoeolog retention rate and the conservation of expression patterns underline the high biological importance of the MIKC-type MADS-box gene family in general and the distinct subclades in particular. Hence, the rich knowledge about the developmental role of MIKC-type MADS-box genes from model plants, combined with a complete picture of gene number, expression data and phylogenetic analyses in wheat will aid wheat breeding and improvement.

### Subfamily-specific expansion of wheat MIKC-type MADS-box genes may contribute to the high adaptability of wheat

The hexaploid nature of wheat and the large size of the MADS-box gene family provide an ideal opportunity to study the evolutionary fate of genes after gene duplication and polyploidization.

With 201 genes, wheat has one of the largest MIKC-type MADS-box gene counts among flowering plants [27, 44, 45]. In total, wheat has about 3.1 times as many transcription factors as rice, which can generally be explained by its hexaploidy [60]. However, the number of MIKC-type MADS-box genes is more than 4.5 times higher in wheat than in rice (Figure 2A-D). The strikingly high number of MIKC-type MADS-box genes observed in this study is mainly due to – leaving hexaploidy aside – the significant expansion of four subclades: *SEP1*, *Bsister*, *AGL17* and *FLC* (Figure 2).

The expansion of MIKC-type subfamilies has been reported before in different plant species, such as for *SOC1*-like genes in *Eucalyptus* as well as *SVP*-like and *SOC1*-like genes in cotton [27, 45]. Intriguingly, genes from *FLC*-, *SVP*- and *SOC1-* subfamilies are involved in the control of flowering time [12, 61, 62]. It has been hypothesized that the expansion (and contraction) of developmental control genes, more specifically eudicot *FLC*-like genes, facilitate the rapid adaptation to changes in environmental factors such as temperature [63]. In a similar manner, duplications of wheat *FLC*-like genes might have enabled the adaptation of wheat to different climatic conditions, therefore contributing to its global distribution. It will be interesting to see whether copy number variations of *FLC*-like genes can be detected in different wheat varieties.

The expansion of *Bsister* and *AGL17*-like genes may similarly be explained with an adaptive advantage. However, in those cases neofunctionalization might be involved. *Bsister* like genes form three subclades of *OsMADS29*-, *OsMADS30-* and *OsMADS31*-like genes in grasses. Expression pattern and evolutionary analyses suggest that grass *OsMADS29*- and *OsMADS31*-like genes retain a conserved role in ovule and fruit development [48, 64], whereas *OsMADS30*-like genes may have functionally diverged [50]. For five wheat *OsMADS30*-like genes, upregulation was observed during infection with Fusarium head blight (*Fusarium graminearum*, Figure 6A). MIKC-type MADS-box genes are not typically associated with biotic stress responses and it remains speculative whether and how they might be involved in responding to a *Fusarium* infection. However, Fusarium head blight is a floral disease and *Bsister* genes are expressed in the flower. This may have facilitated a co-option of theses genes into a pathogen response network. Interestingly, the lack of synteny between *OsMADS30*-like genes might point towards transposable elements as a possible duplication mechanism, underlining the evolutionary importance of transposable elements [65].

An upregulation in response to stresses was also observed for some *AGL17*-like genes, which form the largest wheat MIKC-type subfamily. While many wheat *AGL17*-like genes are not expressed (Figure 5), a number of them are upregulated in late stages of stripe rust infection and in response to heat stress (Figure 6B, C). Another three *AGL17*-like genes were found to be upregulated in some stages of anther and grain development, a pattern unusual for *AGL17*-like genes (Figure 5A). This diversity of expression patterns and putative functions adds to the complex evolution of *AGL17*-like genes, as genes from this subfamily have been implicated in various different functions including root development, flowering time control, tillering, stomata development and stress response [54, 66–69].

*OsMADS30*- and *AGL17*-like genes might be involved in other stress responses as well and might be good candidates to investigate cultivar-specific resistance to biotic and abiotic stresses.

### Dynamic evolution of MIKC-type MADS-box genes in subtelomeric regions

The cause for the expansion of the *FLC*-, *AGL17*- *SEP1*- *OsMADS30*-like subfamilies might be the chromosomal position of their genes. Subtelomeric distal chromosome segments have been previously described as being targets of recombination events, and many fast evolving genes lie within these evolutionary hot spots [46, 70]. In wheat specifically, genes related to stress response and external stimuli, notably traits with a high requirement for adaptability, have been found to be located in distal chromosomal segments [7]. In contrast, genes related to photosynthesis, cell cycle or translation, e.g. genes with involved in highly conserved pathways are enriched in proximal chromosomal segments [7].

This notion is supported by our findings: genes of the larger wheat MIKC-type subclades tend to be located in subtelomeric segments (Figure 4, Figure 3). Remarkably, many of these expanded clades do control traits that are important for adaptation to different environments. For example, *AGL17*-like genes, are involved in root development and stress response and *FLC*-like genes, determine flowering time. On the other hand, smaller MIKC-type subclades involved in highly conserved developmental functions, such as *AP3/PI*- or *AG/STK*-like genes, which control reproductive organ identity, tend to be located more in central chromosomal segments (Figure 3). The higher prevalence of duplication events in subtelomeric chromosomal segments might thus have caused the expansion of certain subclades, possibly facilitating rapid adaptation to different environmental conditions. On the other hand, there might be an evolutionary advantage for wheat MIKC-type subclades with highly conserved functions to be located in proximal chromosomal positions: this way e.g. developmentally detrimental gene dosage variations are minimized.

The high prevalence of gene duplications in subtelomeric segments most likely also led to a higher proportion of truncated genes lacking either the MADS- or K-box (Figure 3A). This might render the encoded protein functionally impaired, as suggested earlier for SEP1.2A (WLHS1A), which lacks a K-domain and shows no protein-protein interactions *in vivo* [71] (Figure 1, Table 1). However, MIKC-type MADS-domain proteins without a K-domain might theoretically still be able to bind DNA [13] and compete with full-length MIKC-type proteins for target sites, thus functioning as transcriptional inhibitors. On the other hand, genes lacking the MADS-box but encoding a K-domain might act in a dominant-negative manner by binding and sequestering other MADS-domain proteins [38, 39]. Evidence that dominant-negative versions of transcriptions factors can be evolutionary and developmentally important comes from basic helix-loop-helix proteins [72, 73]. We found *OsMADS30*-like wheat genes, encoding for only MADS, only K and an unusual combination of MADS and SRP54 domain expressed ubiquitously (Figures 5A, 6A), hence deviating from canonical *Bsister* expression in the flower and again hinting towards a possible neofunctionalization of *OsMADS30*-like genes during wheat evolution.

## Conclusions

MIKC-type MADS-box genes are hugely important for wheat development and hence bear immense potential for the improvement of this economically highly relevant crop. Our data indicate that MADS-box gene duplications might have been crucial for increasing the adaptability of wheat to different environmental conditions as well as for fine-tuning quantitative traits by gene duplication. By thoroughly characterizing the entire complement of wheat MIKC-type MADS box genes, we provide the basis for the development of markers for future breeding efforts as well as for the identification of gene-editing targets to improve wheat performance. Further, we frequently observe possible neofunctionalization, a requirement for understanding the emergence of new traits during evolution.

## Methods

### Sequence search and annotation of MIKC-type MADS-box genes

Wheat coding sequence (CDSs) predictions, refmap comparison between IWGSC and The Genome Analysis Centre (TGAC) and functional annotations of both, high and low confidence (HC and LC) wheat genes, were downloaded from the IWGSC archive v1.0 (https://urgi.versailles.inra.fr/download/iwgsc/IWGSC_RefSeq_Annotations/v1.0/) [7] and the CSS - TGAC comparison was downloaded from https://opendata.earlham.ac.uk/opendata/data/Triticum_aestivum/TGAC/v1/annotation/ [74]. Functional annotations were filtered for PFAM identifiers of the MADS- and K-domain (PF00319 and PF01486), respectively [75]. A total of 439 sequences were identified (see a list of all gene IDs in Table S1). Of these, 188 sequences had a MADS-box and a K-box (181 HC plus 7 LC), 240 sequences (159 HC plus 71 LC) had only a MADS-box; and 21 had only a K- -box (16 HC plus 5 LC) (Table S1). Splice variants were excluded and only the first variant was kept for further analysis, with three exceptions (see Table S2).

All CDSs were translated into amino acid sequences and aligned with all MADS-domain protein sequences of rice (*Oryza sativa*, [21]) with MAFFT (L-INS-i algorithm) [76, 77] using only the MADS-domain of each sequence. Subsequently, a phylogeny was generated using IQTREE [78, 79]. This allowed distinguishing type I and type II (MIKC-type) MADS-domain proteins (Table S1). All type I MADS-box CDSs were excluded from subsequent analyses.

We evaluated the predicted gene structure of all genes, assuming a canonical M-I-K-C domain structure. In cases where either MADS- or K-box were absent from gene predictions, we compared the sequences with closely related rice and Arabidopsis genes, TGAC gene predictions and screened the surrounding genomic regions using the NCBI conserved domain database [80–83]. In 17 cases gene prediction was repeated with FGENESH+ or FGENESH_C [84] using rice MADS-box genes of the same clade (Table S4). In one case (*TaAGL17.2A1*, *TraesCSU01G209900*), the TGAC CDS was used instead of IWGSC prediction, because it encoded for a canonical MIKC structure as compared to the IWGSC prediction, which comprised a MADS-box. This approach yielded 193 wheat MIKC-type MADS-box sequences.

In parallel, a BLAST search was carried out by which we identified 8 additional MIKC-type genes (Table S3) (https://urgi.versailles.inra.fr/blast_iwgsc/) [7, 85].

Altogether, a total of 201 wheat MIKC-type MADS-box genes were identified (Table 1, Table S2).

### Maximum likelihood phylogeny of MIKC-type MADS-box genes

Based on the first phylogeny, MIKC-type sequences were sorted into the major grass MIKC-type subfamilies (Table S2) [40]. Afterwards, subfamily alignments of MIKC-type protein sequences were created using wheat, rice and Arabidopsis protein sequences [24, 42, 43] using MAFFT (E-INS-i algorithm) [76, 77]. Subfamily alignments were then merged using MAFFT (E-INS-i algorithm;) [76, 77]. The full-length alignment of all MIKC-type MADS-domain proteins was analyzed with Homo (Version 1.3) [86] to confirm that the sequences met the phylogenetic assumption of reversible conditions. Individual residues were subsequently masked with Alistat (Version 1.3) [87], leaving only sites with a completeness score (Cc, defined as Cc = Number of unambiguous characters in the column / number of sequences) above 0.5 (a total of 207 sites) (alignment in Supplementary Text 2).

Using the masked protein alignment, a phylogenetic tree was inferred under maximum likelihood with IQ-TREE [79]. The substitution model was calculated with ModelFinder (integrated in IQ-TREE; best-fit model: JTT+R5 chosen according to BIC) [78]. Consistency of the phylogenetic estimate was evaluated with Ultra Fast bootstraps as well as SH-aLRT test (1000 replicates each) [88–90]. The resulting treefile was visualized with Geneious version 11.1 (https://www.geneious.com) (Figure 1) and FigTree version 1.4.4 (http://tree.bio.ed.ac.uk/software/figtree/) (Figure S2).

### Naming of MIKC-type MADS-box genes

We suggest a consistent naming pattern for all MIKC-type wheat MADS-box genes, taking into account their subfamily association, phylogenetic relationships as well as their subgenome location (A, B or D). Each gene name starts with an abbreviation for the species name *Triticum aestivum* (*Ta*), followed by the name of the respective Arabidopsis subfamily (e.g. “*SEP1*” for *SEPALLATA1*-like genes, “*AGL6*” for *AGL6*-like genes or “*BS*” for *Bsister* genes). The gene names include an A, B or D, indicating the subgenome they are located in, e.g. *TaAGL6B*. If more than one triad of homoeologs was found in one subfamily, these were distinguished by a “.” and consecutive numbers, then followed by the subgenome (e.g. *TaAGL12.1A and TaAGL12.2A*). If more than one copy of a gene was present in one subgenome (inparalogs, e.g. due to tandem duplications or transposition), a number was added after the letter that indicates the subgenome (prevalent in *FLC*, *SEP1*, *AGL17* and *Bsister* subclades). Hence, the name of the gene with the ID TraesCS7B01G020900 is *TaSEP1.5B1* since it is a *SEP1*-like gene, and more precisely one of two inparalogs of the B genome (Figure 1, Table 1, all gene names listed in Table S2).

### Identification of homoeologs

Homoeologous genes were identified by phylogeny (Figure 1, Figure S3A-D). In some cases, where Ultra Fast bootstraps and SH-aLRT were not high enough to support a clade (above 90 and 75, respectively), synteny and previous classifications were considered [8]. 38 genes belonging to the *FLC*-, *Bsister* and *AGL17*-subclades were excluded from the analysis, because their homoeolog status could not definitely be determined (Table 1, Table 2, Figure S3).

### Plant cultivation, RNA isolation and RT-PCR

*Triticum turgidum* cv. Kronos (tetraploid) was used for RT-PCR analysis. Plants were grown in a greenhouse with no additional lighting at an ambient temperature of 20 to 24 °C until heading stage. RNA was extracted from the frozen flag leaf tip using QIAGEN^®^ RNeasy Mini Kit according to manufacturer’s instructions. cDNA was generated with Superscript IV^®^ Reverse Transcriptase Kit using an oligo dT primer. RT-PCR was carried out with Thermo Scientific™ Phusion High-Fidelity DNA Polymerase. Primer sequences are listed in Table S5.

### Expression analysis of MIKC-type MADS-box genes using RNA-seq

RNA-seq data of 193 wheat MADS-box genes was analyzed using www.wheatexpression.com and http://bar.utoronto.ca/efp_wheat/ [8]. For the remaining 8 genes, which were identified by BLAST, no expression data was available. Developmental stages depicted in Figure 5 refer to 70 tissues/time points from spring wheat cv. Azhurnaya [8]. Expression levels were downloaded from www.wheatexpression.com as log_2_tpm and a heatmap was generated with heatmapper (<u>heatmapper.ca/expression</u>) [91]. Clustering was performed with heatmapper using centroid linkage with euclidean distance measure. All genes, modules and tissues are listed in Tables S6 and S7.

## Declarations

### Ethics approval and consent to participate

Not applicable

### Consent for publication

Not applicable

### Availability of data and material

Not applicable

## Authors’ contributions

RM and SS developed the analysis approach. SS and AK analyzed the data with supervision from RM and LJ. SP performed the RT-PCR experiments. SS and RM wrote the manuscript. All authors read and approved the final manuscript.

## Acknowledgements

We are grateful for the support by the International Wheat Genome Sequencing Consortium and early access to the genome data.

## Supplementary Material

### Supplementary Figures

**Figure S1. RT-PCR of two FLC-like wheat genes.** Schematic representation of two FLC genes and primer binding sites (red arrowheads) (A). Intron size is indicated by dotted line, boxes indicate exons, lines indicates introns. Gel images of PCR products and expected fragment sizes (B). The sequence of the PCR products was verified using Sanger Sequencing.

**Figure S2. Maximum likelihood phylogeny of MIKC-type MADS-domain proteins from bread wheat, rice and Arabidopsis, branches not transformed.** SH-aLRT and Ultrafast bootstrap values are indicated on the branches in %, values equal to 100 are designated by a -. The tree is unrooted, the MIKC* subclade was set as the outgroup.

**Figure S3. Maximum likelihood phylogenies of four different MIKC-type subfamilies.** Subphylogenies of *SEPALLATA*- (A), *Bsister*- (B), *AGL17*- (C) and *FLC*- (D) like genes from Arabidopsis, rice and wheat were generated using protein alignments and IQ-TREE [78, 79]. *AGL6*- (A), *AP3/PI*-(B) and *OsMADS32*- (C, D)-like genes were used as an outgroup. Red and blue triangles indicate truncated genes encoding only for MADS- and K-domain, respectively. Black circle indicates gene encoding for a MADS- and SRP54-domain (PF00448). Groups excluded from homoeolog analysis are indicated by asterisks.

**Figure S4. Cluster expression analysis of wheat MIKC-type MADS-box genes during developmental stages.** Expression analysis was done for all wheat MIKC-type genes and tissue using RNA-seq data from wheatexpression.com [8, 47]. A heatmap shows expression levels of all genes in different subfamilies (rows) and stages/tissues (columns). Genes and tissues are listed in Tables S6 and S7, respectively. Heatmap and clustering was done with heatmapper.com and a cladogram showing the result of the clustering is shown on the left. Modules 1 to 17 were assigned and borders indicated by red lines (right). Values represent log_2_tpm.

**Figure S5. Expression of wheat *Bsister* genes during Fusarium head blight infection.** RNA-seq data was analyzed using wheatexpression.com [8, 47]. *Bsister* gene expression in the donor plant and 4 near-isogenic lines (NIL) under control conditions (c) and *Fusarium graminearum* inoculation (s) (30 and 50 h) is shown as a heatmap [96]. Values represent log_2_tpm.

### Supplementary Tables

**Table S1. List of all sequences identified by PFAM domain (type I and type II).**

**Table S2. List of all MIKC-type MADS-box genes in wheat.**

^1^ (+) indicates FLC-specific K-box (not PFAM PF01486); (-) TGAC version of this gene does have PF01486, IWGSC does not have PF01486

^2^ Protein length and molecular weight were determined with the protein molecular weight function on bioinformatics.org using conceptually translated CDS.

^3^ According to [97]

^4^ According to [8]

^5^ TGAC identifiers were inferred from refmap comparison between IWGSC and TGAC [7] (details see Methods)

^6^ identifiers were inferred from refmap comparison between CSS - TGAC [74] (details see Methods)

**Table S3. Average distance to the nearest downstream MIKC-type gene in kbp in different chromosomal segments.** Distances between start points of neighboring MIKC-type genes have been calculated and averaged over chromosomal segments.

**Table S4. List of all MIKC-type MADS-box genes predicted during this study.** Table includes gene name, IWGSC identifier (if available) and chromosome location, PFAM domain(s), predicted coding sequence and prediction algorithm and guiding sequence FGENESH+ and FGENESH_C predictions are guided by homolog protein or cDNA sequences, respectively.

**Table S5. Primer sequences for RT-PCR.**

**Table S6. Expression modules and gene numbers for Figure 5A and S3.**

**Table S7. Tissues for expression analysis for Figure 5A and S3.**

