## Supplemental Material for "Genome-wide analysis of MIKC-type MADS-box genes in wheat: pervasive duplications may have facilitated adaptation to different environmental conditions"

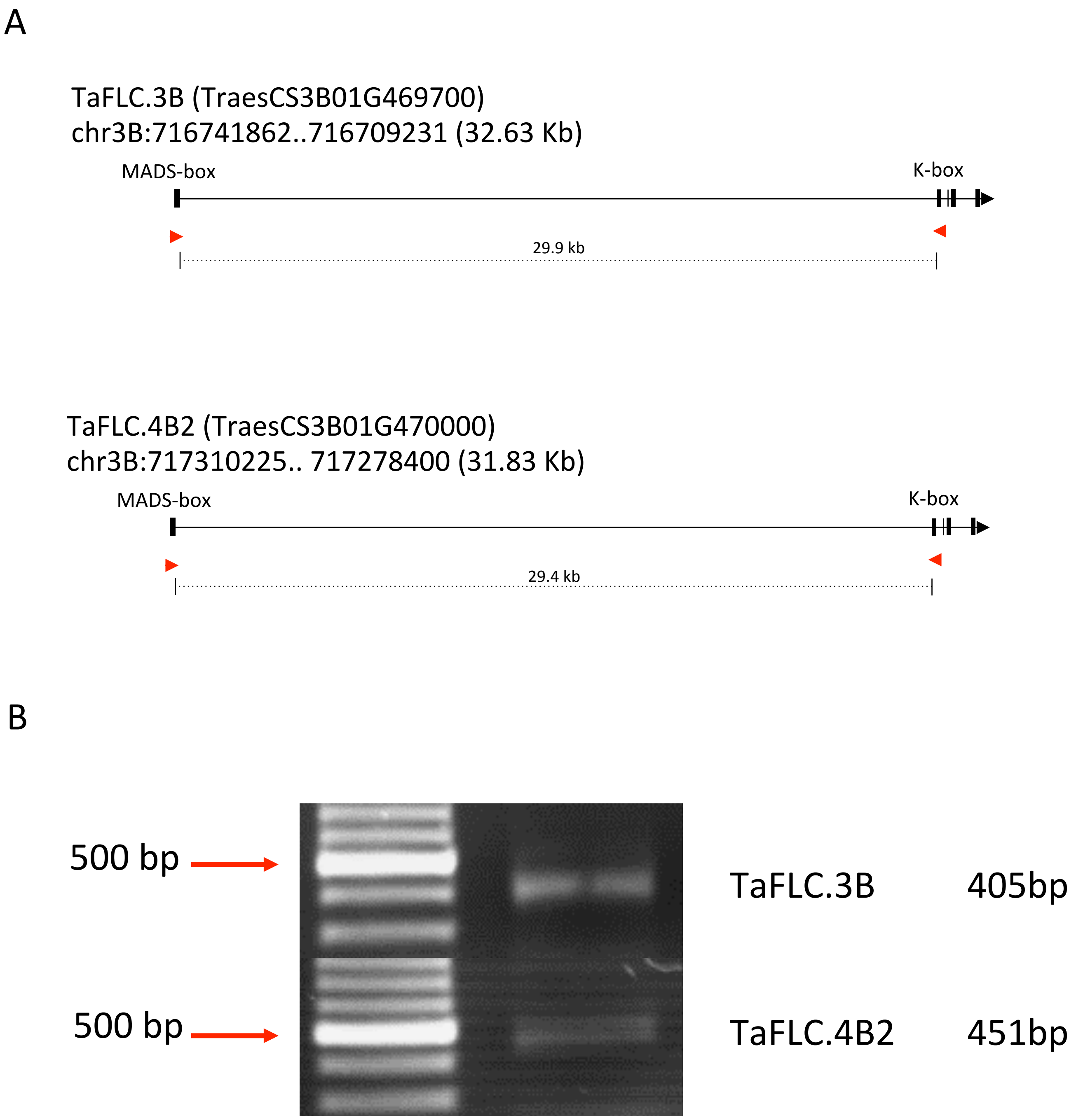


**Figure S1. RT-PCR of two FLC-like wheat genes.** Schematic representation of two FLC genes and primer binding sites (red arrowheads) (A). Intron size is indicated by dotted line, boxes indicate exons, lines indicates introns. Gel images of PCR products and expected fragment sizes (B). The sequence of the PCR products was verified using Sanger Sequencing.

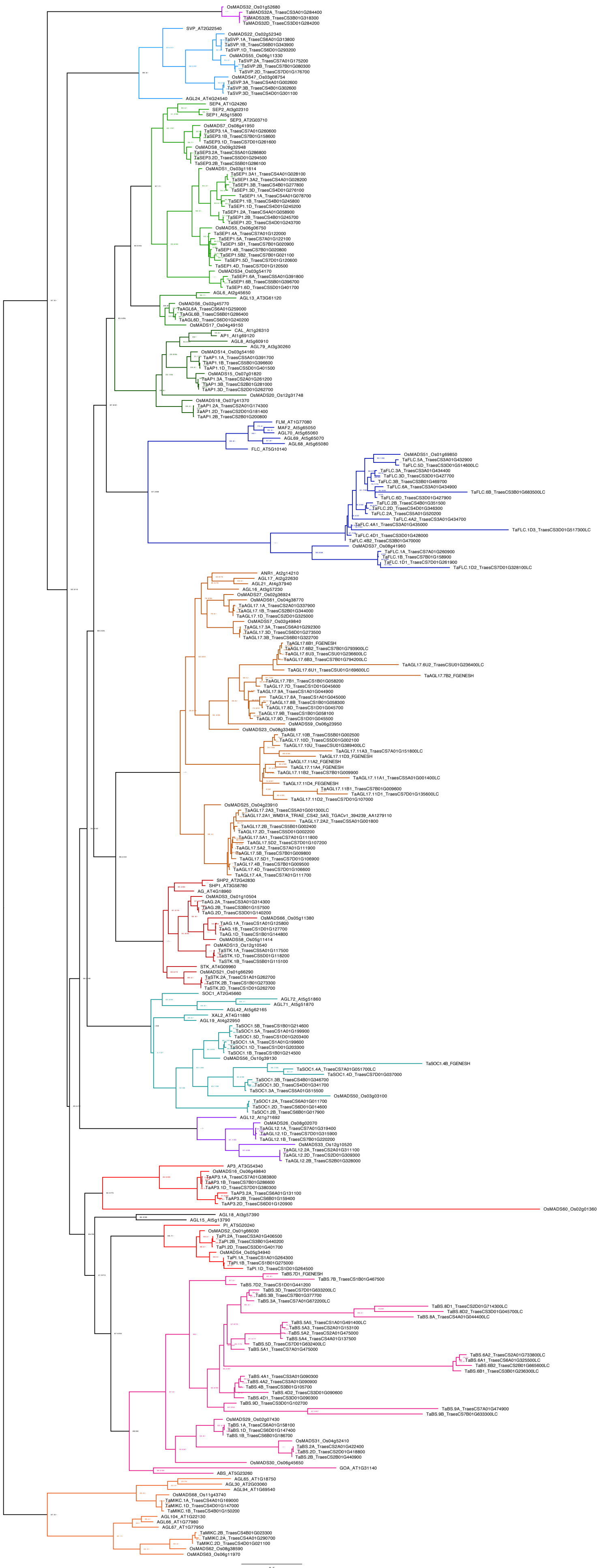

**Figure S2. Maximum likelihood phylogeny of MIKC-type MADS-domain proteins from bread wheat, rice and Arabidopsis, branches not transformed.** SH-aLRT and Ultrafast bootstrap values are indicated on the branches in %, values equal to 100 are designated by a -. The tree is unrooted, the MIKC* subclade was set as the outgroup.

A

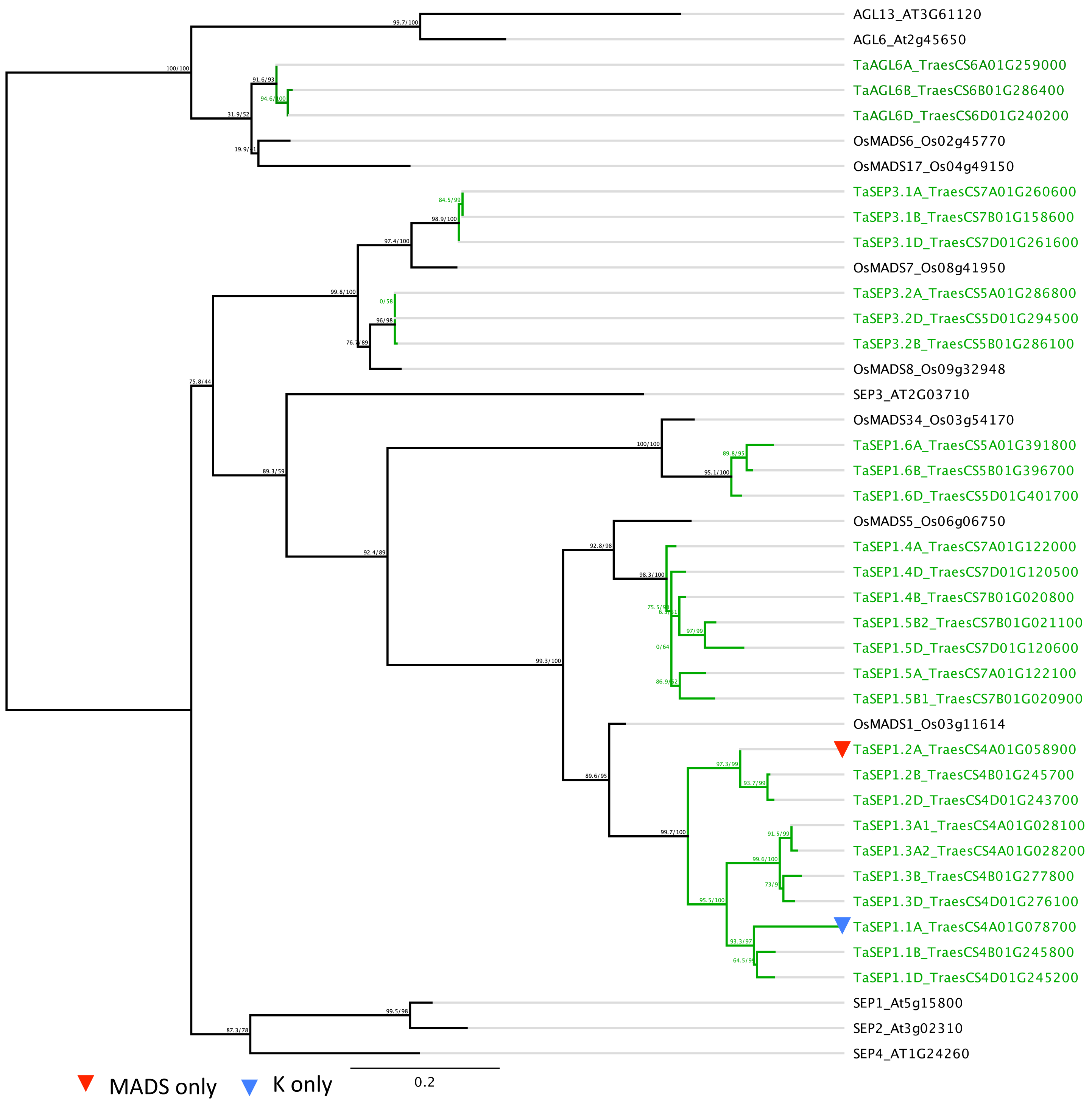
AGL13_AT3G61120
AGL6_At2g45650
TaAGL6A_TraesCS6A01G259000
TaAGL6B_TraesCS6B01G286400
TaAGL6D_TraesCS6D01G240200
OsMADS6_Os02g45770
OsMADS17_Os04g49150
TaSEP3.1A_TraesCS7A01G260600
TaSEP3.1B_TraesCS7B01G158600
TaSEP3.1D_TraesCS7D01G261600
OsMADS7_Os08g41950
TaSEP3.2A_TraesCS5A01G286800
TaSEP3.2D_TraesCS5D01G294500
TaSEP3.2B_TraesCS5B01G286100
OsMADS8_Os09g32948
SEP3_AT2G03710
OsMADS34_Os03g54170
TaSEP1.6A_TraesCS5A01G391800
TaSEP1.6B_TraesCS5B01G396700
TaSEP1.6D_TraesCS5D01G401700
OsMADS5_Os06g06750
TaSEP1.4A_TraesCS7A01G122000
TaSEP1.4D_TraesCS7D01G120500
TaSEP1.4B_TraesCS7B01G020800
TaSEP1.5B2_TraesCS7B01G021100
TaSEP1.5D_TraesCS7D01G120600
TaSEP1.5A_TraesCS7A01G122100
TaSEP1.5B1_TraesCS7B01G020900
OsMADS1_Os03g11614
TaSEP1.2A_TraesCS4A01G058900
TaSEP1.2B_TraesCS4B01G245700
TaSEP1.2D_TraesCS4D01G243700
TaSEP1.3A1_TraesCS4A01G028100
TaSEP1.3A2_TraesCS4A01G028200
TaSEP1.3B_TraesCS4B01G277800
TaSEP1.3D_TraesCS4D01G276100
TaSEP1.1A_TraesCS4A01G078700
TaSEP1.1B_TraesCS4B01G245800
TaSEP1.1D_TraesCS4D01G245200
SEP1_At5g15800
SEP2_At3g02310
SEP4_AT1G24260
MADS only
K only
0.2

B

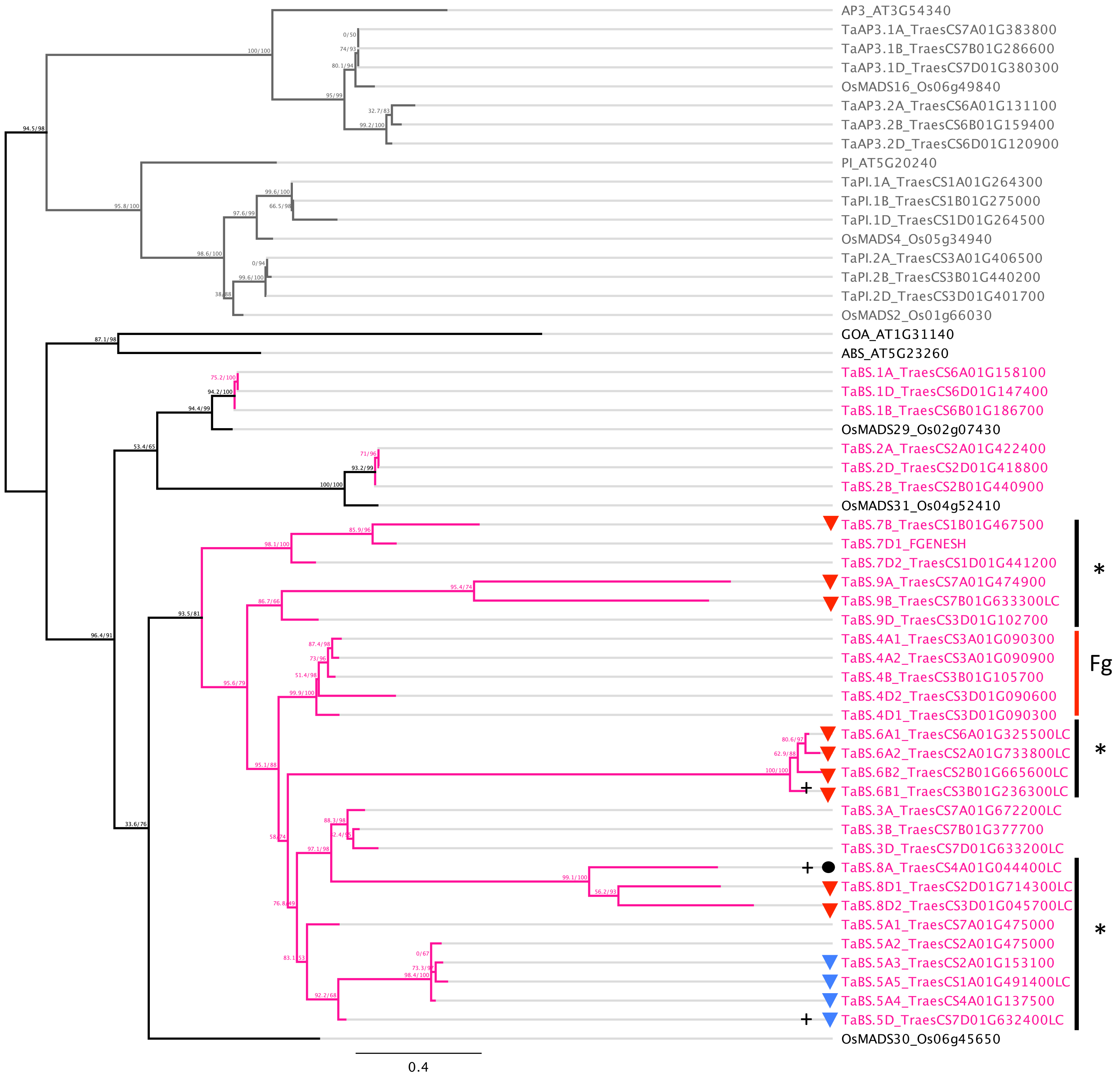


● SRP54 domain (PF00448)

▼ MADS only ▼ K only

* excluded from homoeolog analysis

Fg expressed during *F. graminearum* response

\+ expressed ubiquitiously

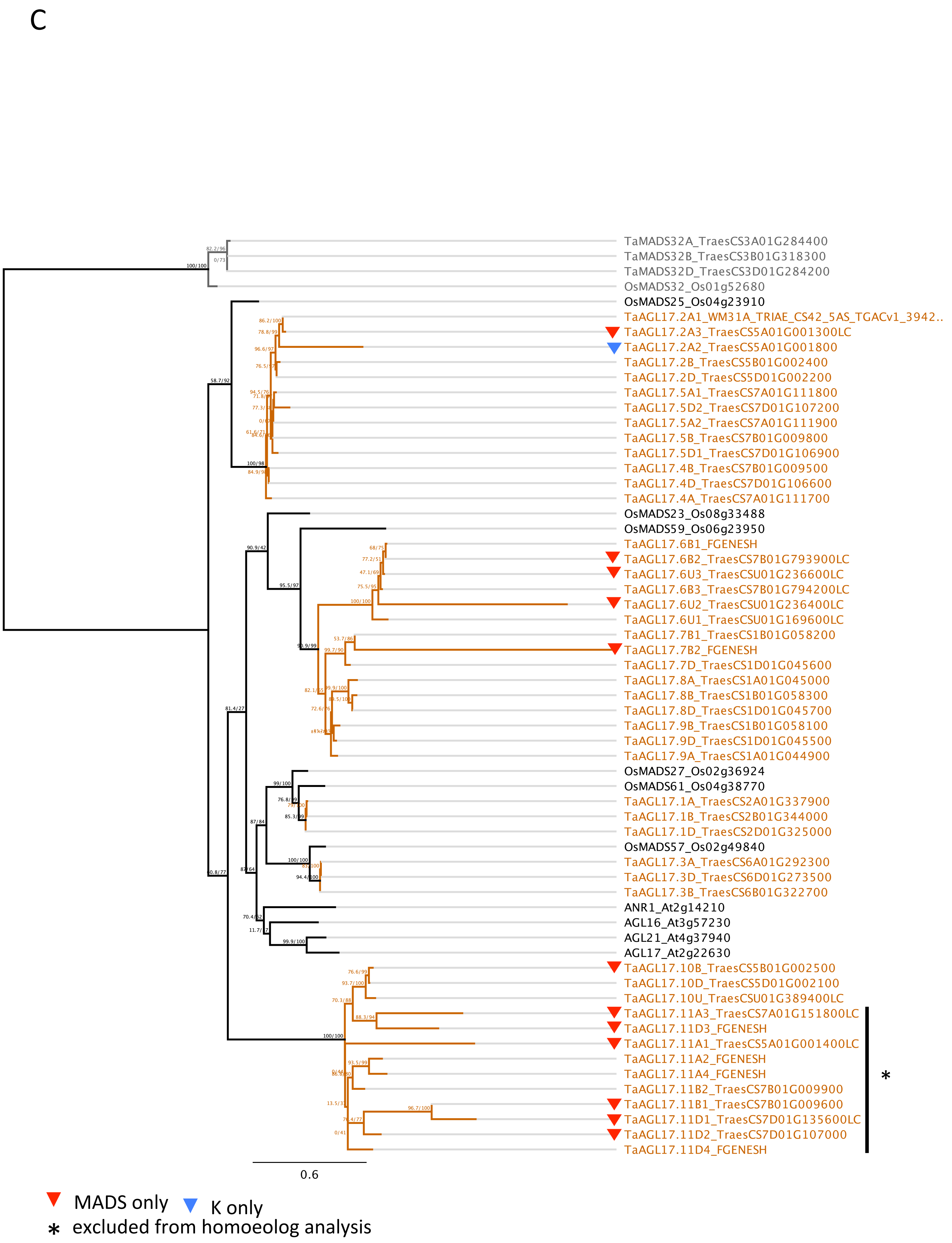
C
TaMADS32A_TraesCS3A01G284400
TaMADS32B_TraesCS3B01G318300
TaMADS32D_TraesCS3D01G284200
OsMADS32_Os01g52680
OsMADS25_Os04g23910
TaAGL17.2A1_WM31A_TRIAE_CS42_5AS_TGACv1_3942..
TaAGL17.2A3_TraesCS5A01G001300LC
TaAGL17.2A2_TraesCS5A01G001800
TaAGL17.2B_TraesCS5B01G002400
TaAGL17.2D_TraesCS5D01G002200
TaAGL17.5A1_TraesCS7A01G111800
TaAGL17.5D2_TraesCS7D01G107200
TaAGL17.5A2_TraesCS7A01G111900
TaAGL17.5B_TraesCS7B01G009800
TaAGL17.5D1_TraesCS7D01G106900
TaAGL17.4B_TraesCS7B01G009500
TaAGL17.4D_TraesCS7D01G106600
TaAGL17.4A_TraesCS7A01G111700
OsMADS23_Os08g33488
OsMADS59_Os06g23950
TaAGL17.6B1_FGENESH
TaAGL17.6B2_TraesCS7B01G793900LC
TaAGL17.6U3_TraesCSU01G236600LC
TaAGL17.6B3_TraesCS7B01G794200LC
TaAGL17.6U2_TraesCSU01G236400LC
TaAGL17.6U1_TraesCSU01G169600LC
TaAGL17.7B1_TraesCS1B01G058200
TaAGL17.7B2_FGENESH
TaAGL17.7D_TraesCS1D01G045600
TaAGL17.8A_TraesCS1A01G045000
TaAGL17.8B_TraesCS1B01G058300
TaAGL17.8D_TraesCS1D01G045700
TaAGL17.9B_TraesCS1B01G058100
TaAGL17.9D_TraesCS1D01G045500
TaAGL17.9A_TraesCS1A01G044900
OsMADS27_Os02g36924
OsMADS61_Os04g38770
TaAGL17.1A_TraesCS2A01G337900
TaAGL17.1B_TraesCS2B01G344000
TaAGL17.1D_TraesCS2D01G325000
OsMADS57_Os02g49840
TaAGL17.3A_TraesCS6A01G292300
TaAGL17.3D_TraesCS6D01G273500
TaAGL17.3B_TraesCS6B01G322700
ANR1_At2g14210
AGL16_At3g57230
AGL21_At4g37940
AGL17_At2g22630
TaAGL17.10B_TraesCS5B01G002500
TaAGL17.10D_TraesCS5D01G002100
TaAGL17.10U_TraesCSU01G389400LC
TaAGL17.11A3_TraesCS7A01G151800LC
TaAGL17.11D3_FGENESH
TaAGL17.11A1_TraesCS5A01G001400LC
TaAGL17.11A2_FGENESH
TaAGL17.11A4_FGENESH
TaAGL17.11B2_TraesCS7B01G009900
TaAGL17.11B1_TraesCS7B01G009600
TaAGL17.11D1_TraesCS7D01G135600LC
TaAGL17.11D2_TraesCS7D01G107000
TaAGL17.11D4_FGENESH
*
0.6
MADS only
K only
* excluded from homoeolog analysis

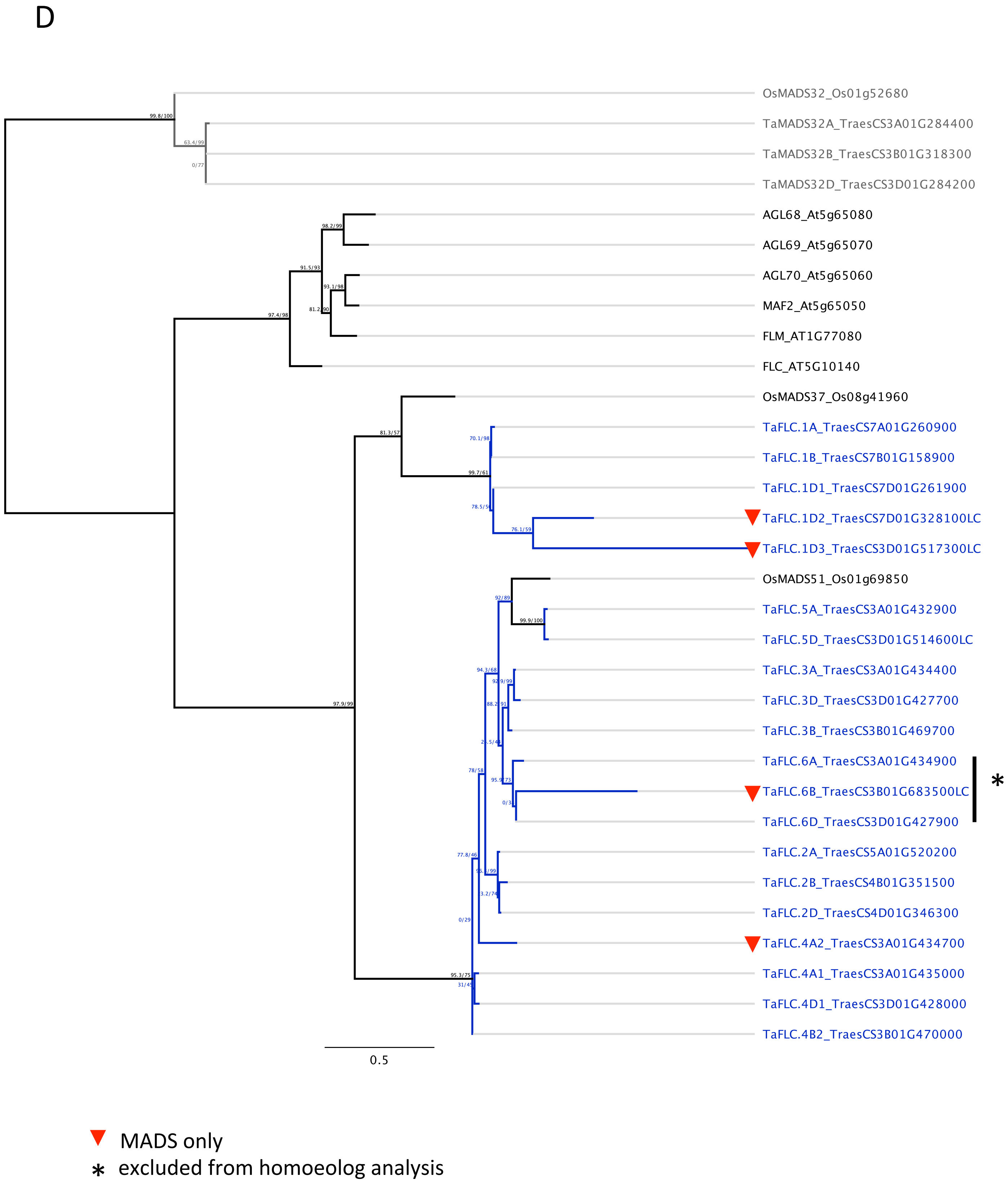


**Figure S3. Maximum likelihood phylogenies of four different MIKC-type subfamilies.** Subphylogenies of *SEPALLATA*- (A), *Bsister*- (B), *AGL17*- (C) and *FLC*- (D) like genes from Arabidopsis, rice and wheat were generated using protein alignments and IQ-TREE [78, 79]. *AGL6*- (A), *AP3/PI*-(B) and *OsMADS32*- (C, D)-like genes were used as an outgroup. Red and blue triangles indicate truncated genes encoding only for MADS- and K-domain, respectively. Black circle indicates gene encoding for a MADS- and SRP54-domain (PF00448). Groups excluded from homoeolog analysis are indicated by asterisks.

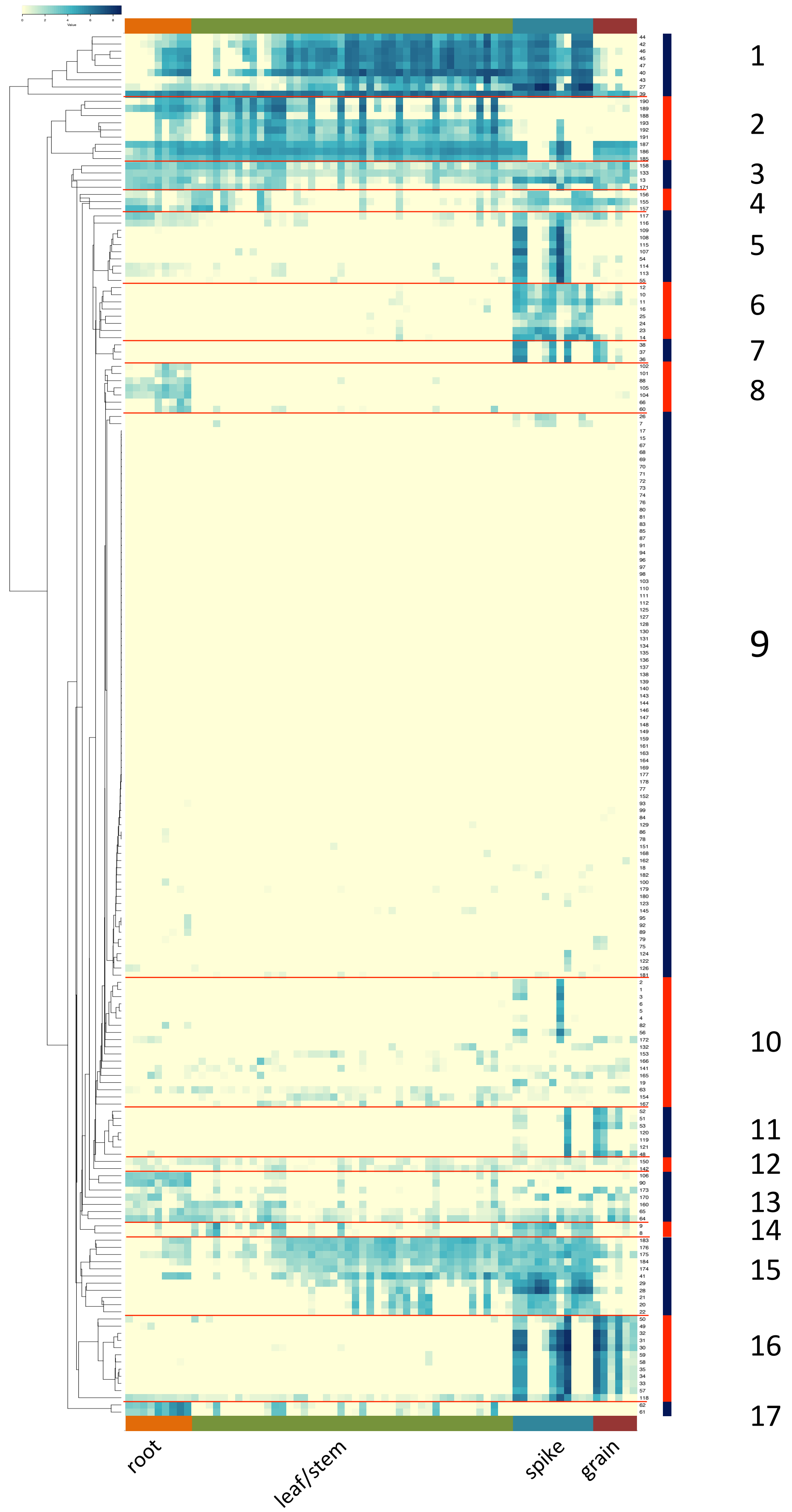


**Figure S4. Cluster expression analysis of wheat MIKC-type MADS-box genes during developmental stages.** Expression analysis was done for all wheat MIKC-type genes and tissue using RNA-seq data from wheatexpression.com [8, 47]. A heatmap shows expression levels of all genes in different subfamilies (rows) and stages/tissues (columns). Genes and tissues are listed in Tables S6 and S7, respectively. Heatmap and clustering was done with heatmapper.com and a cladogram showing the result of the clustering is shown on the left. Modules 1 to 17 were assigned and borders indicated by red lines (right). Values represent $log_2$tpm.

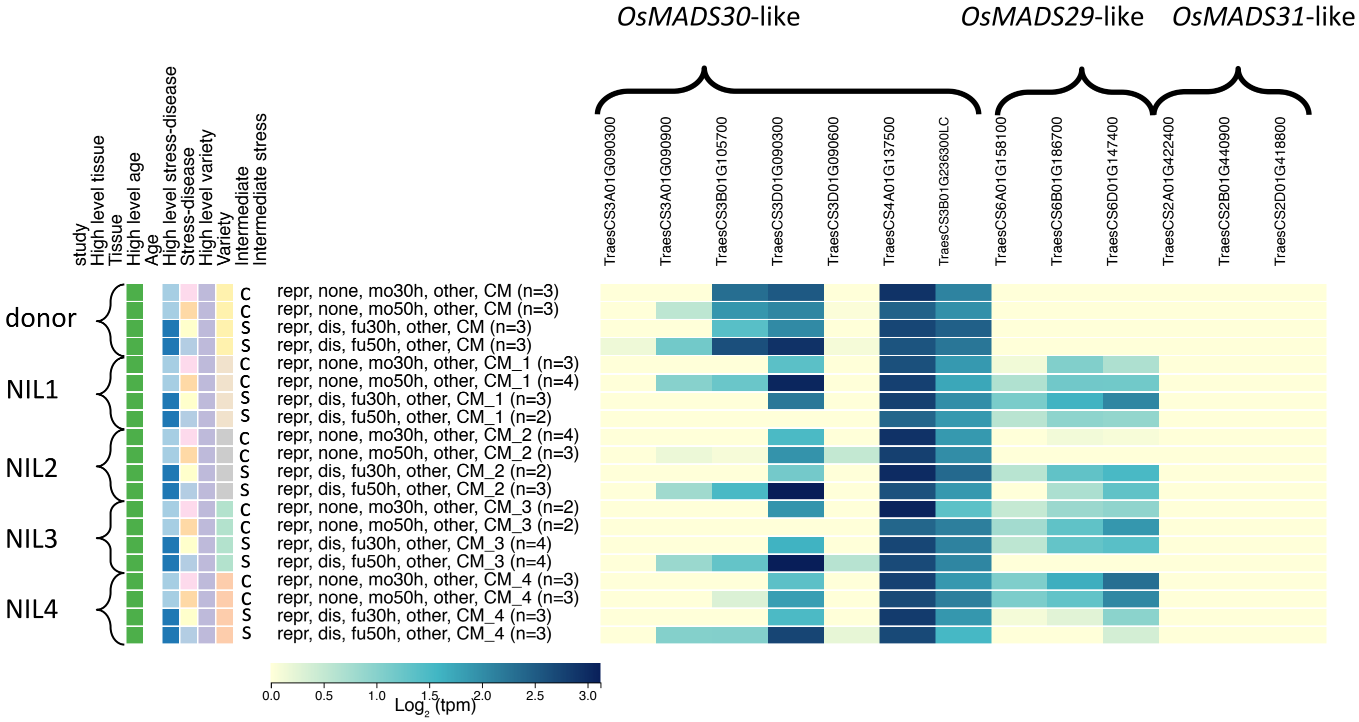


**Figure S5. Expression of wheat *Bsister* genes during Fusarium head blight infection.** RNA-seq data was analyzed using wheatexpression.com [8, 47]. *Bsister* gene expression in the donor plant and 4 near-isogenic lines (NIL) under control conditions (c) and *Fusarium graminearum* inoculation (s) (30 and 50 h) is shown as a heatmap [96]. Values represent $log_2$tpm.

| Gene-ID | Confidence | Pfam-IDs | typeI or type II MADS-box gene |
|---|---|---|---|
| TraesCS1A01G013100.1 | HC | PF00319 | 1 |
| TraesCS1A01G061800.1 | HC | PF00319 | 1 |
| TraesCS1A01G310800.1 | HC | PF00319 | 1 |
| TraesCS1B01G080200.1 | HC | PF00319 | 1 |
| TraesCS1B01G322300.1 | HC | PF00319 | 1 |
| TraesCS1D01G062500.1 | HC | PF00319 | 1 |
| TraesCS1D01G310700.1 | HC | PF00319 | 1 |
| TraesCS2A01G146400.1 | HC | PF00319 | 1 |
| TraesCS2A01G283500.1 | HC | PF00319 | 1 |
| TraesCS2A01G459200.1 | HC | PF00319 | 1 |
| TraesCS2A01G523900.1 | HC | PF00319 | 1 |
| TraesCS2B01G146100.1 | HC | PF00319 | 1 |
| TraesCS2B01G171700.1 | HC | PF00319 | 1 |
| TraesCS2B01G300500.1 | HC | PF00319 | 1 |
| TraesCS2D01G128000.1 | HC | PF00319 | 1 |
| TraesCS2D01G150900.1 | HC | PF00319 | 1 |
| TraesCS2D01G282200.1 | HC | PF00319 | 1 |
| TraesCS2D01G526000.1 | HC | PF00319 | 1 |
| TraesCS2D01G526200.1 | HC | PF00319 | 1 |
| TraesCS3A01G137500.1 | HC | PF00319 | 1 |
| TraesCS3A01G312600.1 | HC | PF00319 | 1 |
| TraesCS3A01G359300.1 | HC | PF00319 | 1 |
| TraesCS3A01G417800.1 | HC | PF00319 | 1 |
| TraesCS3A01G454500.1 | HC | PF00319 | 1 |
| TraesCS3A01G527400.1 | HC | PF00319 | 1 |
| TraesCS3A01G535200.1 | HC | PF00319 | 1 |
| TraesCS3B01G155200.1 | HC | PF00319 | 1 |
| TraesCS3B01G159600.1 | HC | PF00319 | 1 |
| TraesCS3B01G453000.1 | HC | PF00319 | 1 |
| TraesCS3B01G454700.1 | HC | PF00319 | 1 |
| TraesCS3B01G493100.1 | HC | PF00319 | 1 |
| TraesCS3B01G595700.1 | HC | PF00319 | 1 |
| TraesCS3B01G608600.1 | HC | PF00319 | 1 |
| TraesCS3B01G612300.1 | HC | PF00319 | 1 |
| TraesCS3B01G612400.1 | HC | PF00319 | 1 |
| TraesCS3B01G612500.1 | HC | PF00319 | 1 |
| TraesCS3B01G612600.1 | HC | PF00319 | 1 |
| TraesCS3B01G612700.1 | HC | PF00319 | 1 |
| TraesCS3B01G612800.1 | HC | PF00319 | 1 |
| TraesCS3B01G612900.1 | HC | PF00319 | 1 |
| TraesCS3D01G138400.1 | HC | PF00319 | 1 |
| TraesCS3D01G446900.1 | HC | PF00319 | 1 |
| TraesCS3D01G513800.1 | HC | PF00319 | 1 |
| TraesCS3D01G532900.1 | HC | PF00319 | 1 |
| TraesCS3D01G540700.1 | HC | PF00319 | 1 |
| TraesCS4A01G403600.1 | HC | PF00319 | 1 |
| TraesCS5A01G010400.1 | HC | PF00319 | 1 |
| TraesCS5B01G008000.1 | HC | PF00319 | 1 |
| TraesCS5B01G471900.1 | HC | PF00319 | 1 |
| TraesCS5D01G474300.1 | HC | PF00319 | 1 |
| TraesCS6A01G020500.1 | HC | PF00319 | 1 |
| TraesCS6A01G020900.1 | HC | PF00319 | 1 |
| TraesCS6A01G022700.1 | HC | PF00319 | 1 |
| TraesCS6A01G138300.1 | HC | PF00319 | 1 |
| TraesCS6A01G338400.1 | HC | PF00319 | 1 |
| TraesCS6A01G358900.1 | HC | PF00319 | 1 |
| TraesCS6A01G359100.1 | HC | PF00319 | 1 |
| TraesCS6A01G359700.1 | HC | PF00319 | 1 |
| TraesCS6B01G027900.1 | HC | PF00319 | 1 |
| TraesCS6B01G028100.1 | HC | PF00319 | 1 |
| TraesCS6B01G028300.1 | HC | PF00319 | 1 |
| TraesCS6B01G369000.1 | HC | PF00319 | 1 |
| TraesCS6B01G369100.1 | HC | PF00319 | 1 |
| TraesCS6B01G391600.1 | HC | PF00319 | 1 |
| TraesCS6B01G391700.1 | HC | PF00319 | 1 |
| TraesCS6B01G391800.1 | HC | PF00319 | 1 |
| TraesCS6B01G392600.1 | HC | PF00319 | 1 |
| TraesCS6D01G023800.1 | HC | PF00319 | 1 |
| TraesCS6D01G024000.1 | HC | PF00319 | 1 |
| TraesCS6D01G024300.1 | HC | PF00319 | 1 |
| TraesCS6D01G319100.1 | HC | PF00319 | 1 |
| TraesCS6D01G341700.1 | HC | PF00319 | 1 |
| TraesCS6D01G341800.1 | HC | PF00319 | 1 |
| TraesCS6D01G342000.1 | HC | PF00319 | 1 |
| TraesCS6D01G342100.1 | HC | PF00319 | 1 |
| TraesCS6D01G342700.1 | HC | PF00319 | 1 |
| TraesCS7A01G142600.1 | HC | PF00319 | 1 |
| TraesCS7A01G335700.1 | HC | PF00319 | 1 |
| TraesCS7A01G342300.1 | HC | PF00319 | 1 |
| TraesCS7A01G342600.1 | HC | PF00319 | 1 |
| TraesCS7A01G367400.1 | HC | PF00319 | 1 |
| TraesCS7A01G367500.1 | HC | PF00319 | 1 |
| TraesCS7A01G393100.1 | HC | PF00319 | 1 |
| TraesCS7A01G398800.1 | HC | PF00319 | 1 |
| TraesCS7A01G551000.1 | HC | PF00319 | 1 |
| TraesCS7A01G558000.1 | HC | PF00319 | 1 |
| TraesCS7A01G559500.1 | HC | PF00319 | 1 |
| TraesCS7A01G561000.1 | HC | PF00319 | 1 |
| TraesCS7B01G042900.1 | HC | PF00319 | 1 |
| TraesCS7B01G043000.1 | HC | PF00319 | 1 |
| TraesCS7B01G211300.1 | HC | PF00319 | 1 |
| TraesCS7B01G240400.1 | HC | PF00319 | 1 |
| TraesCS7B01G240500.1 | HC | PF00319 | 1 |
| TraesCS7B01G247300.1 | HC | PF00319 | 1 |
| TraesCS7B01G295100.1 | HC | PF00319 | 1 |
| TraesCS7B01G295200.1 | HC | PF00319 | 1 |
| TraesCS7B01G299400.1 | HC | PF00319 | 1 |
| TraesCS7B01G482700.1 | HC | PF00319 | 1 |
| TraesCS7B01G483300.1 | HC | PF00319 | 1 |
| TraesCS7B01G484000.1 | HC | PF00319 | 1 |
| TraesCS7B01G484100.1 | HC | PF00319 | 1 |
| TraesCS7B01G484500.1 | HC | PF00319 | 1 |
| TraesCS7B01G484600.1 | HC | PF00319 | 1 |
| TraesCS7B01G485400.1 | HC | PF00319 | 1 |
| TraesCS7D01G143800.1 | HC | PF00319 | 1 |
| TraesCS7D01G337000.1 | HC | PF00319 | 1 |
| TraesCS7D01G337200.1 | HC | PF00319 | 1 |
| TraesCS7D01G343400.1 | HC | PF00319 | 1 |
| TraesCS7D01G355300.1 | HC | PF00319 | 1 |
| TraesCS7D01G388600.1 | HC | PF00319 | 1 |
| TraesCS7D01G393000.1 | HC | PF00319 | 1 |
| TraesCS7D01G536700.1 | HC | PF00319 | 1 |
| TraesCS7D01G548500.1 | HC | PF00319 | 1 |
| TraesCS7D01G550500.1 | HC | PF00319 | 1 |
| TraesCS7D01G550600.1 | HC | PF00319 | 1 |
| TraesCS7D01G551000.1 | HC | PF00319 | 1 |
| TraesCS7D01G551900.1 | HC | PF00319 | 1 |
| TraesCS7D01G552000.1 | HC | PF00319 | 1 |
| TraesCS7D01G552400.1 | HC | PF00319 | 1 |
| TraesCSU01G235300.1 | HC | PF00319 | 1 |
| TraesCS1A01G085000LC.1 | LC | PF00319 | 1 |
| TraesCS1A01G101400LC.1 | LC | PF00319 | 1 |
| TraesCS1A01G456600LC.1 | LC | PF00319 | 1 |
| TraesCS1B01G555600LC.1 | LC | PF00319 | 1 |
| TraesCS1B01G555900LC.1 | LC | PF00319 | 1 |
| TraesCS2A01G128800LC.1 | LC | PF00319 | 1 |
| TraesCS2A01G149300LC.1 | LC | PF00319 | 1 |
| TraesCS2A01G685300LC.1 | LC | PF00319 | 1 |
| TraesCS2B01G239100LC.1 | LC | PF00319 | 1 |
| TraesCS2B01G825100LC.1 | LC | PF00319 | 1 |
| TraesCS2D01G166900LC.1 | LC | PF00319 | 1 |
| TraesCS2D01G692400LC.1 | LC | PF00319 | 1 |
| TraesCS3A01G498500LC.1 | LC | PF00319 | 1 |
| TraesCS3A01G686700LC.1 | LC | PF00319 | 1 |
| TraesCS3A01G732000LC.1 | LC | PF00319 | 1 |
| TraesCS3B01G579800LC.1 | LC | PF00319 | 1 |
| TraesCS3B01G658400LC.1 | LC | PF00319 | 1 |
| TraesCS3B01G906200LC.1 | LC | PF00319 | 1 |
| TraesCS3B01G906300LC.1 | LC | PF00319 | 1 |
| TraesCS3B01G906400LC.1 | LC | PF00319 | 1 |
| TraesCS3D01G159700LC.1 | LC | PF00319 | 1 |
| TraesCS3D01G434600LC.1 | LC | PF00319 | 1 |
| TraesCS3D01G499900LC.1 | LC | PF00319 | 1 |
| TraesCS3D01G653600LC.1 | LC | PF00319 | 1 |
| TraesCS5A01G024600LC.1 | LC | PF00319 | 1 |
| TraesCS5D01G018600LC.1 | LC | PF00319 | 1 |
| TraesCS6A01G559100LC.1 | LC | PF00319 | 1 |
| TraesCS6B01G253300LC.1 | LC | PF00319 | 1 |
| TraesCS6D01G154600LC.1 | LC | PF00319 | 1 |
| TraesCS7A01G142000LC.1 | LC | PF00319 | 1 |
| TraesCS7A01G190000LC.1 | LC | PF00319 | 1 |
| TraesCS7A01G190200LC.1 | LC | PF00319 | 1 |
| TraesCS7A01G553100LC.1 | LC | PF00319 | 1 |
| TraesCS7A01G555400LC.1 | LC | PF00319 | 1 |
| TraesCS7A01G618600LC.1 | LC | PF00319 | 1 |
| TraesCS7A01G796600LC.1 | LC | PF00319 | 1 |
| TraesCS7A01G798100LC.1 | LC | PF00319 | 1 |
| TraesCS7B01G453100LC.1 | LC | PF00319 | 1 |
| TraesCS7B01G809500LC.1 | LC | PF00319 | 1 |
| TraesCS7B01G809800LC.1 | LC | PF00319 | 1 |
| TraesCS7B01G810300LC.1 | LC | PF00319 | 1 |
| TraesCS7D01G125700LC.1 | LC | PF00319 | 1 |
| TraesCS7D01G524700LC.1 | LC | PF00319 | 1 |
| TraesCS7D01G525700LC.1 | LC | PF00319 | 1 |
| TraesCS7D01G581300LC.1 | LC | PF00319 | 1 |
| TraesCS7D01G742100LC.1 | LC | PF00319 | 1 |
| TraesCS7D01G742600LC.1 | LC | PF00319 | 1 |
| TraesCSU01G085400LC.1 | LC | PF00319 | 1 |
| TraesCSU01G249400LC.1 | LC | PF00319 | 1 |
| TraesCSU01G491900LC.1 | LC | PF00319 | 1 |
| TraesCSU01G494900LC.1 | LC | PF00319 | 1 |
| TraesCSU01G524600LC.1 | LC | PF00319 | 1 |
| TraesCSU01G526800LC.1 | LC | PF00319 | 1 |
| TraesCSU01G595200LC.1 | LC | PF00319 | 1 |
| TraesCS1B01G214600.1 | HC | PF00319 | 2 |
| TraesCS1B01G467500.1 | HC | PF00319 | 2 |
| TraesCS3A01G432900.1 | HC | PF00319 | 2 |
| TraesCS3A01G434400.1 | HC | PF00319 | 2 |
| TraesCS3A01G434700.1 | HC | PF00319 | 2 |
| TraesCS3A01G434900.1 | HC | PF00319 | 2 |
| TraesCS3A01G435000.1 | HC | PF00319 | 2 |
| TraesCS3B01G469700.1 | HC | PF00319 | 2 |
| TraesCS3B01G470000.1 | HC | PF00319 | 2 |

| | | | |
|---|---|---|---|
| TraesCS3D01G090600.1 | HC | PF00319 | 2 |
| TraesCS3D01G427700.1 | HC | PF00319 | 2 |
| TraesCS3D01G427900.1 | HC | PF00319 | 2 |
| TraesCS3D01G428000.1 | HC | PF00319 | 2 |
| TraesCS4A01G058900.1 | HC | PF00319 | 2 |
| TraesCS4A01G058900.2 | HC | PF00319 | 2 |
| TraesCS4A01G169000.1 | HC | PF00319 | 2 |
| TraesCS4A01G290700.1 | HC | PF00319 | 2 |
| TraesCS4B01G023300.1 | HC | PF00319 | 2 |
| TraesCS4B01G150200.1 | HC | PF00319 | 2 |
| TraesCS4B01G150200.2 | HC | PF00319 | 2 |
| TraesCS4B01G351500.1 | HC | PF00319 | 2 |
| TraesCS4D01G021100.1 | HC | PF00319 | 2 |
| TraesCS4D01G147000.1 | HC | PF00319 | 2 |
| TraesCS4D01G147000.2 | HC | PF00319 | 2 |
| TraesCS4D01G346300.1 | HC | PF00319 | 2 |
| TraesCS5A01G001800.1 | HC | PF00319 | 2 |
| TraesCS5A01G520200.1 | HC | PF00319 | 2 |
| TraesCS5B01G002500.1 | HC | PF00319 | 2 |
| TraesCS7A01G260900.1 | HC | PF00319 | 2 |
| TraesCS7A01G474900.1 | HC | PF00319 | 2 |
| TraesCS7B01G009600.1 | HC | PF00319 | 2 |
| TraesCS7B01G158900.1 | HC | PF00319 | 2 |
| TraesCS7D01G037000.1 | HC | PF00319 | 2 |
| TraesCS7D01G107000.1 | HC | PF00319 | 2 |
| TraesCS7D01G107200.1 | HC | PF00319 | 2 |
| TraesCS7D01G261900.1 | HC | PF00319 | 2 |
| TraesCSU01G209900.1 | HC | PF00319 | 2 |
| TraesCS2A01G733800LC.1 | LC | PF00319 | 2 |
| TraesCS2B01G665600LC.1 | LC | PF00319 | 2 |
| TraesCS2D01G714300LC.1 | LC | PF00319 | 2 |
| TraesCS3B01G236300LC.1 | LC | PF00319 | 2 |
| TraesCS3B01G683500LC.1 | LC | PF00319 | 2 |
| TraesCS3D01G045700LC.1 | LC | PF00319 | 2 |
| TraesCS3D01G514600LC.1 | LC | PF00319 | 2 |
| TraesCS3D01G517300LC.1 | LC | PF00319 | 2 |
| TraesCS5A01G001400LC.1 | LC | PF00319 | 2 |
| TraesCS6A01G325500LC.1 | LC | PF00319 | 2 |
| TraesCS7A01G151800LC.1 | LC | PF00319 | 2 |
| TraesCS7B01G633300LC.1 | LC | PF00319 | 2 |
| TraesCS7B01G793900LC.1 | LC | PF00319 | 2 |
| TraesCS7D01G135600LC.1 | LC | PF00319 | 2 |
| TraesCS7D01G328100LC.1 | LC | PF00319 | 2 |
| TraesCSU01G236400LC.1 | LC | PF00319 | 2 |
| TraesCSU01G236600LC.1 | LC | PF00319 | 2 |
| TraesCS7B01G020900.1 | HC | PF00319 | 2 |
| TraesCS7B01G021100.1 | HC | PF00319 | 2 |
| TraesCS4A01G044400LC.1 | LC | PF00319; PF00448 | 2 |
| TraesCS1A01G044900.1 | HC | PF00319; PF01486 | 2 |
| TraesCS1A01G045000.1 | HC | PF00319; PF01486 | 2 |
| TraesCS1A01G125800.1 | HC | PF00319; PF01486 | 2 |
| TraesCS1A01G125800.2 | HC | PF00319; PF01486 | 2 |
| TraesCS1A01G199600.1 | HC | PF00319; PF01486 | 2 |
| TraesCS1A01G199600.2 | HC | PF00319; PF01486 | 2 |
| TraesCS1A01G199900.1 | HC | PF00319; PF01486 | 2 |
| TraesCS1A01G262700.1 | HC | PF00319; PF01486 | 2 |
| TraesCS1A01G262700.2 | HC | PF00319; PF01486 | 2 |
| TraesCS1A01G264300.1 | HC | PF00319; PF01486 | 2 |
| TraesCS1B01G058100.1 | HC | PF00319; PF01486 | 2 |
| TraesCS1B01G058200.1 | HC | PF00319; PF01486 | 2 |
| TraesCS1B01G058300.1 | HC | PF00319; PF01486 | 2 |
| TraesCS1B01G144800.1 | HC | PF00319; PF01486 | 2 |
| TraesCS1B01G144800.3 | HC | PF00319; PF01486 | 2 |
| TraesCS1B01G214500.1 | HC | PF00319; PF01486 | 2 |
| TraesCS1B01G273300.1 | HC | PF00319; PF01486 | 2 |
| TraesCS1B01G275000.1 | HC | PF00319; PF01486 | 2 |
| TraesCS1D01G045500.1 | HC | PF00319; PF01486 | 2 |
| TraesCS1D01G045600.1 | HC | PF00319; PF01486 | 2 |
| TraesCS1D01G045700.1 | HC | PF00319; PF01486 | 2 |
| TraesCS1D01G127700.1 | HC | PF00319; PF01486 | 2 |
| TraesCS1D01G127700.2 | HC | PF00319; PF01486 | 2 |
| TraesCS1D01G203300.1 | HC | PF00319; PF01486 | 2 |
| TraesCS1D01G203300.2 | HC | PF00319; PF01486 | 2 |
| TraesCS1D01G203400.1 | HC | PF00319; PF01486 | 2 |
| TraesCS1D01G262700.1 | HC | PF00319; PF01486 | 2 |
| TraesCS1D01G264500.1 | HC | PF00319; PF01486 | 2 |
| TraesCS1D01G264500.2 | HC | PF00319; PF01486 | 2 |
| TraesCS1D01G441200.1 | HC | PF00319; PF01486 | 2 |
| TraesCS2A01G174300.3 | HC | PF00319; PF01486 | 2 |
| TraesCS2A01G261200.1 | HC | PF00319; PF01486 | 2 |
| TraesCS2A01G261200.2 | HC | PF00319; PF01486 | 2 |
| TraesCS2A01G311100.1 | HC | PF00319; PF01486 | 2 |
| TraesCS2A01G337900.1 | HC | PF00319; PF01486 | 2 |
| TraesCS2A01G422400.1 | HC | PF00319; PF01486 | 2 |
| TraesCS2A01G422400.2 | HC | PF00319; PF01486 | 2 |
| TraesCS2B01G200800.1 | HC | PF00319; PF01486 | 2 |
| TraesCS2B01G281000.1 | HC | PF00319; PF01486 | 2 |
| TraesCS2B01G328000.1 | HC | PF00319; PF01486 | 2 |
| TraesCS2B01G344000.1 | HC | PF00319; PF01486 | 2 |
| TraesCS2B01G440900.1 | HC | PF00319; PF01486 | 2 |
| TraesCS2D01G181400.3 | HC | PF00319; PF01486 | 2 |
| TraesCS2D01G262700.1 | HC | PF00319; PF01486 | 2 |
| TraesCS2D01G309300.1 | HC | PF00319; PF01486 | 2 |
| TraesCS2D01G325000.1 | HC | PF00319; PF01486 | 2 |
| TraesCS2D01G418800.1 | HC | PF00319; PF01486 | 2 |
| TraesCS2D01G418800.2 | HC | PF00319; PF01486 | 2 |
| TraesCS3A01G090300.1 | HC | PF00319; PF01486 | 2 |
| TraesCS3A01G090900.1 | HC | PF00319; PF01486 | 2 |
| TraesCS3A01G284400.1 | HC | PF00319; PF01486 | 2 |
| TraesCS3A01G314300.1 | HC | PF00319; PF01486 | 2 |
| TraesCS3A01G314300.2 | HC | PF00319; PF01486 | 2 |
| TraesCS3A01G314300.3 | HC | PF00319; PF01486 | 2 |
| TraesCS3A01G406500.1 | HC | PF00319; PF01486 | 2 |
| TraesCS3A01G406500.2 | HC | PF00319; PF01486 | 2 |
| TraesCS3B01G105700.1 | HC | PF00319; PF01486 | 2 |
| TraesCS3B01G105700.2 | HC | PF00319; PF01486 | 2 |
| TraesCS3B01G157500.1 | HC | PF00319; PF01486 | 2 |
| TraesCS3B01G157500.2 | HC | PF00319; PF01486 | 2 |
| TraesCS3B01G157500.3 | HC | PF00319; PF01486 | 2 |
| TraesCS3B01G157500.4 | HC | PF00319; PF01486 | 2 |
| TraesCS3B01G318300.1 | HC | PF00319; PF01486 | 2 |
| TraesCS3B01G440200.1 | HC | PF00319; PF01486 | 2 |
| TraesCS3B01G440200.2 | HC | PF00319; PF01486 | 2 |
| TraesCS3B01G440200.3 | HC | PF00319; PF01486 | 2 |
| TraesCS3D01G090300.1 | HC | PF00319; PF01486 | 2 |
| TraesCS3D01G102700.1 | HC | PF00319; PF01486 | 2 |
| TraesCS3D01G140200.1 | HC | PF00319; PF01486 | 2 |
| TraesCS3D01G140200.2 | HC | PF00319; PF01486 | 2 |
| TraesCS3D01G140200.3 | HC | PF00319; PF01486 | 2 |
| TraesCS3D01G140200.4 | HC | PF00319; PF01486 | 2 |
| TraesCS3D01G284200.1 | HC | PF00319; PF01486 | 2 |
| TraesCS3D01G401700.1 | HC | PF00319; PF01486 | 2 |
| TraesCS3D01G401700.2 | HC | PF00319; PF01486 | 2 |
| TraesCS4A01G028100.1 | HC | PF00319; PF01486 | 2 |
| TraesCS4A01G028200.1 | HC | PF00319; PF01486 | 2 |
| TraesCS4B01G245700.1 | HC | PF00319; PF01486 | 2 |
| TraesCS4B01G245800.1 | HC | PF00319; PF01486 | 2 |
| TraesCS4B01G277800.1 | HC | PF00319; PF01486 | 2 |
| TraesCS4B01G302600.1 | HC | PF00319; PF01486 | 2 |
| TraesCS4B01G302600.2 | HC | PF00319; PF01486 | 2 |
| TraesCS4B01G302600.3 | HC | PF00319; PF01486 | 2 |
| TraesCS4B01G346700.1 | HC | PF00319; PF01486 | 2 |
| TraesCS4D01G243700.1 | HC | PF00319; PF01486 | 2 |
| TraesCS4D01G245200.1 | HC | PF00319; PF01486 | 2 |
| TraesCS4D01G276100.1 | HC | PF00319; PF01486 | 2 |
| TraesCS4D01G301100.1 | HC | PF00319; PF01486 | 2 |
| TraesCS4D01G341700.1 | HC | PF00319; PF01486 | 2 |
| TraesCS5A01G117500.1 | HC | PF00319; PF01486 | 2 |
| TraesCS5A01G117500.2 | HC | PF00319; PF01486 | 2 |
| TraesCS5A01G286800.1 | HC | PF00319; PF01486 | 2 |
| TraesCS5A01G391700.1 | HC | PF00319; PF01486 | 2 |
| TraesCS5A01G391700.2 | HC | PF00319; PF01486 | 2 |
| TraesCS5A01G391800.1 | HC | PF00319; PF01486 | 2 |
| TraesCS5A01G515500.1 | HC | PF00319; PF01486 | 2 |
| TraesCS5B01G002400.1 | HC | PF00319; PF01486 | 2 |
| TraesCS5B01G115100.1 | HC | PF00319; PF01486 | 2 |
| TraesCS5B01G115100.2 | HC | PF00319; PF01486 | 2 |
| TraesCS5B01G286100.1 | HC | PF00319; PF01486 | 2 |
| TraesCS5B01G286100.2 | HC | PF00319; PF01486 | 2 |
| TraesCS5B01G396600.1 | HC | PF00319; PF01486 | 2 |
| TraesCS5B01G396600.2 | HC | PF00319; PF01486 | 2 |
| TraesCS5B01G396700.1 | HC | PF00319; PF01486 | 2 |
| TraesCS5D01G002100.1 | HC | PF00319; PF01486 | 2 |
| TraesCS5D01G002200.1 | HC | PF00319; PF01486 | 2 |
| TraesCS5D01G002200.2 | HC | PF00319; PF01486 | 2 |
| TraesCS5D01G118200.1 | HC | PF00319; PF01486 | 2 |
| TraesCS5D01G118200.2 | HC | PF00319; PF01486 | 2 |
| TraesCS5D01G294500.1 | HC | PF00319; PF01486 | 2 |
| TraesCS5D01G401500.1 | HC | PF00319; PF01486 | 2 |
| TraesCS5D01G401500.2 | HC | PF00319; PF01486 | 2 |
| TraesCS5D01G401500.3 | HC | PF00319; PF01486 | 2 |
| TraesCS5D01G401700.1 | HC | PF00319; PF01486 | 2 |
| TraesCS6A01G011700.1 | HC | PF00319; PF01486 | 2 |
| TraesCS6A01G011700.2 | HC | PF00319; PF01486 | 2 |
| TraesCS6A01G131100.1 | HC | PF00319; PF01486 | 2 |
| TraesCS6A01G158100.1 | HC | PF00319; PF01486 | 2 |
| TraesCS6A01G259000.1 | HC | PF00319; PF01486 | 2 |
| TraesCS6A01G259000.2 | HC | PF00319; PF01486 | 2 |
| TraesCS6A01G259000.3 | HC | PF00319; PF01486 | 2 |
| TraesCS6A01G292300.1 | HC | PF00319; PF01486 | 2 |
| TraesCS6A01G292300.2 | HC | PF00319; PF01486 | 2 |
| TraesCS6A01G313800.1 | HC | PF00319; PF01486 | 2 |
| TraesCS6B01G017900.1 | HC | PF00319; PF01486 | 2 |
| TraesCS6B01G159400.1 | HC | PF00319; PF01486 | 2 |
| TraesCS6B01G186700.1 | HC | PF00319; PF01486 | 2 |
| TraesCS6B01G286400.1 | HC | PF00319; PF01486 | 2 |
| TraesCS6B01G286400.2 | HC | PF00319; PF01486 | 2 |
| TraesCS6B01G286400.3 | HC | PF00319; PF01486 | 2 |
| TraesCS6B01G322700.1 | HC | PF00319; PF01486 | 2 |
| TraesCS6B01G322700.2 | HC | PF00319; PF01486 | 2 |
| TraesCS6B01G343900.1 | HC | PF00319; PF01486 | 2 |
| TraesCS6D01G014600.1 | HC | PF00319; PF01486 | 2 |
| TraesCS6D01G120900.1 | HC | PF00319; PF01486 | 2 |
| TraesCS6D01G147400.1 | HC | PF00319; PF01486 | 2 |

| | | | |
|---|---|---|---|
| TraesCS6D01G240200.1 | HC | PF00319; PF01486 | 2 |
| TraesCS6D01G240200.2 | HC | PF00319; PF01486 | 2 |
| TraesCS6D01G240200.3 | HC | PF00319; PF01486 | 2 |
| TraesCS6D01G273500.1 | HC | PF00319; PF01486 | 2 |
| TraesCS6D01G273500.2 | HC | PF00319; PF01486 | 2 |
| TraesCS6D01G293200.1 | HC | PF00319; PF01486 | 2 |
| TraesCS7A01G111700.1 | HC | PF00319; PF01486 | 2 |
| TraesCS7A01G111700.2 | HC | PF00319; PF01486 | 2 |
| TraesCS7A01G111800.1 | HC | PF00319; PF01486 | 2 |
| TraesCS7A01G111900.1 | HC | PF00319; PF01486 | 2 |
| TraesCS7A01G122000.1 | HC | PF00319; PF01486 | 2 |
| TraesCS7A01G122100.1 | HC | PF00319; PF01486 | 2 |
| TraesCS7A01G122100.2 | HC | PF00319; PF01486 | 2 |
| TraesCS7A01G175200.1 | HC | PF00319; PF01486 | 2 |
| TraesCS7A01G175200.2 | HC | PF00319; PF01486 | 2 |
| TraesCS7A01G260600.1 | HC | PF00319; PF01486 | 2 |
| TraesCS7A01G260600.2 | HC | PF00319; PF01486 | 2 |
| TraesCS7A01G319400.1 | HC | PF00319; PF01486 | 2 |
| TraesCS7A01G319400.2 | HC | PF00319; PF01486 | 2 |
| TraesCS7A01G383800.1 | HC | PF00319; PF01486 | 2 |
| TraesCS7A01G475000.1 | HC | PF00319; PF01486 | 2 |
| TraesCS7B01G009500.1 | HC | PF00319; PF01486 | 2 |
| TraesCS7B01G009800.1 | HC | PF00319; PF01486 | 2 |
| TraesCS7B01G009900.1 | HC | PF00319; PF01486 | 2 |
| TraesCS7B01G009900.2 | HC | PF00319; PF01486 | 2 |
| TraesCS7B01G020800.1 | HC | PF00319; PF01486 | 2 |
| TraesCS7B01G080300.1 | HC | PF00319; PF01486 | 2 |
| TraesCS7B01G080300.2 | HC | PF00319; PF01486 | 2 |
| TraesCS7B01G158600.1 | HC | PF00319; PF01486 | 2 |
| TraesCS7B01G220200.1 | HC | PF00319; PF01486 | 2 |
| TraesCS7B01G220200.2 | HC | PF00319; PF01486 | 2 |
| TraesCS7B01G286600.1 | HC | PF00319; PF01486 | 2 |
| TraesCS7B01G286600.2 | HC | PF00319; PF01486 | 2 |
| TraesCS7B01G377700.1 | HC | PF00319; PF01486 | 2 |
| TraesCS7D01G106600.1 | HC | PF00319; PF01486 | 2 |
| TraesCS7D01G106900.1 | HC | PF00319; PF01486 | 2 |
| TraesCS7D01G120500.1 | HC | PF00319; PF01486 | 2 |
| TraesCS7D01G120600.1 | HC | PF00319; PF01486 | 2 |
| TraesCS7D01G176700.1 | HC | PF00319; PF01486 | 2 |
| TraesCS7D01G261600.1 | HC | PF00319; PF01486 | 2 |
| TraesCS7D01G261600.2 | HC | PF00319; PF01486 | 2 |
| TraesCS7D01G315900.1 | HC | PF00319; PF01486 | 2 |
| TraesCS7D01G315900.2 | HC | PF00319; PF01486 | 2 |
| TraesCS7D01G315900.3 | HC | PF00319; PF01486 | 2 |
| TraesCS7D01G380300.1 | HC | PF00319; PF01486 | 2 |
| TraesCS7A01G051700LC.1 | LC | PF00319; PF01486 | 2 |
| TraesCS7A01G672200LC.1 | LC | PF00319; PF01486 | 2 |
| TraesCS7B01G794200LC.1 | LC | PF00319; PF01486 | 2 |
| TraesCS7D01G632400LC.1 | LC | PF00319; PF01486 | 2 |
| TraesCSU01G169600LC.1 | LC | PF00319; PF01486 | 2 |
| TraesCSU01G389400LC.1 | LC | PF00319; PF01486 | 2 |
| TraesCS1B01G144800.2 | HC | PF01486 | 2 |
| TraesCS2A01G153100.1 | HC | PF01486 | 2 |
| TraesCS2A01G174300.1 | HC | PF01486 | 2 |
| TraesCS2A01G174300.2 | HC | PF01486 | 2 |
| TraesCS2A01G475000.1 | HC | PF01486 | 2 |
| TraesCS2D01G181400.1 | HC | PF01486 | 2 |
| TraesCS2D01G181400.2 | HC | PF01486 | 2 |
| TraesCS2D01G262700.2 | HC | PF01486 | 2 |
| TraesCS4A01G002600.1 | HC | PF01486 | 2 |
| TraesCS4A01G078700.1 | HC | PF01486 | 2 |
| TraesCS4A01G078700.2 | HC | PF01486 | 2 |
| TraesCS4A01G137500.1 | HC | PF01486 | 2 |
| TraesCS4D01G243700.2 | HC | PF01486 | 2 |
| TraesCS6B01G343900.2 | HC | PF01486 | 2 |
| TraesCS6D01G293200.2 | HC | PF01486 | 2 |
| TraesCS7B01G021000.1 | HC | PF01486 | 2 |
| TraesCS1A01G491400LC.1 | LC | PF01486 | 2 |
| TraesCS1B01G406400LC.1 | LC | PF01486 | 2 |
| TraesCS5A01G001300LC.1 | LC | PF01486 | 2 |
| TraesCS5B01G002300LC.1 | LC | PF01486 | 2 |
| TraesCS7B01G032300LC.1 | LC | PF01486 | 2 |

| Clade | Gene name | Con | IWGSC ID, Ref seq 1.0 | splice variant | number of splice variants | rice ortholog | PFAM ID | M | K[1] | protein length (aa)[2] | protein weight (kDa)[2] | RNA seq[3] | tpm [4] | TGAC ID[5] | CSS IWGSS ID[6] | chr | start | stop | Distance to next MIKC in kb | gene len | Chr region | gene prediction by | Ma 2017 | Paolacci 2007 | Shi 2016 | NCBI genbank | Accession numb | GenBank Name |
|---|---|---|---|---|---|---|---|---|---|---|---|---|---|---|---|---|---|---|---|---|---|---|---|---|---|---|---|---|
| SEP1 | TaSEP1.1A | HC | TraesCS4A01G078700 | 1 | 2 | OsMADS1 | PF01486 | - | + | 219 | 25.19 | y | 29.12 | TRIAE_CS42_4AS_TGACv1_306423_AA1007920 | Traes_4AS_9A9C84490.1, Traes_4AS_BE | chr4A | 79051380 | 79054826 | 117728.23 | 3446 | R2a | IWGSC | TaMADS76, TaMADS79 | TaSEP-1 | | | | |
| SEP1 | TaSEP1.1B | HC | TraesCS4B01G245800 | 1 | 1 | OsMADS1 | PF00319; PF01486 | + | + | 247 | 28.28 | y | 25.65 | TRIAE_CS42_4BL_TGACv1_321103_AA1055390 | Traes_4BS_59C6DEC88.1 | chr4B | 508424397 | 508438351 | 51209.114 | 13954 | R2b | IWGSC | TaMADS86 | TaSEP-1 | | | | |
| SEP1 | TaSEP1.1D | HC | TraesCS4D01G245200 | 1 | 1 | OsMADS1 | PF00319; PF01486 | + | + | 248 | 28.51 | y | 23.08 | TRIAE_CS42_4DL_TGACv1_344241_AA1145340 | Traes_4DL_008F8BBFA.1, Traes_4DL_541 | chr4D | 410402499 | 410413305 | 36869.962 | 10806 | R2b | IWGSC | TaMADS88 | TaSEP-1 | | | AM502866 | |
| SEP1 | TaSEP1.2A | HC | TraesCS4A01G058900 | 1 | 2 | OsMADS1-1 | PF00319 | + | - | 170 | 19.21 | y | 35.59 | TRIAE_CS42_4AS_TGACv1_306359_AA1007020 | | chr4A | 52605554 | 52617528 | 26433.852 | 11974 | R2a | IWGSC | TaMADS89 | TaSEP-2A | TaMADWLHS1-A | WM4 | AM502867 | AB295662,AB295 |
| SEP1 | TaSEP1.2B | HC | TraesCS4B01G245700 | 1 | 1 | OsMADS1-1 | PF00319; PF01486 | + | + | 216 | 24.96 | y | 0.51 | TRIAE_CS42_4BL_TGACv1_322135_AA1068830 | | chr4B | 507964484 | 507982935 | 441.462 | 18451 | R2b | IWGSC | TaMADS80 | TaSEP-2B | TaMADAGL24 | WM5A | AM502868 | |
| SEP1 | TaSEP1.2D | HC | TraesCS4D01G243700 | 1 | 2 | OsMADS1-1 | PF00319; PF01486 | + | + | 225 | 26.01 | y | 49.22 | TRIAE_CS42_4DL_TGACv1_343498_AA1135400 | | chr4D | 407116087 | 407128655 | 3273.844 | 12568 | R2b | IWGSC | | | | | | |
| SEP1 | TaSEP1.3A1 | HC | TraesCS4A01G028100 | 1 | 1 | OsMADS1-2 | PF00319; PF01486 | + | + | 242 | 27.90 | y | 9.13 | TRIAE_CS42_4AS_TGACv1_307811_AA1023620 | | chr4A | 19945645 | 19958329 | 32.989 | 12684 | R1 | IWGSC | | | | WM5B | | |
| SEP1 | TaSEP1.3A2 | HC | TraesCS4A01G028200 | 1 | 1 | OsMADS1-2 | PF00319; PF01486 | + | + | 232 | 26.86 | n | 0.50 | TRIAE_CS42_4AS_TGACv1_308208_AA1026620 | | chr4A | 19991318 | 20007018 | 18730.418 | 15700 | R1 | IWGSC | | | | | | |
| SEP1 | TaSEP1.3B | HC | TraesCS4B01G277800 | 1 | 1 | OsMADS1-2 | PF00319; PF01486 | + | + | 247 | 28.36 | y | 1.58 | TRIAE_CS42_4BL_TGACv1_320477_AA1040830 | | chr4B | 559647465 | 559661486 | 29518.523 | 14021 | R3 | IWGSC | | | | | | |
| SEP1 | TaSEP1.3D | HC | TraesCS4D01G276100 | 1 | 1 | OsMADS1-2 | PF00319; PF01486 | + | + | 241 | 27.87 | y | 18.81 | TRIAE_CS42_4DL_TGACv1_343863_AA1140660 | Traes_4DL_964466BEC.1 | chr4D | 447283267 | 447293095 | 22011.106 | 9828 | R3 | IWGSC | TaMADS92 | | | | | |
| SEP1 | TaSEP1.4A | HC | TraesCS7A01G122000 | 1 | 1 | OsMADS5-1 | PF00319; PF01486 | + | + | 227 | 26.39 | y | 171.32 | TRIAE_CS42_7AS_TGACv1_571056_AA1843970 | Traes_7AS_ADA694FCE.1 | chr7A | 78511991 | 78520747 | 47.737 | 8756 | R1 | IWGSC | TaMADS146 | | | | | DQ512362 |
| SEP1 | TaSEP1.4B | HC | TraesCS7B01G020800 | 1 | 1 | OsMADS5-1 | PF00319; PF01486 | + | + | 227 | 26.20 | y | 104.03 | TRIAE_CS42_7BS_TGACv1_592464_AA1938550 | Traes_7BS_592EC3AB1.1, Traes_7BS_7C | chr7B | 18207990 | 18218252 | 279.997 | 10262 | R2a | IWGSC | TaMADS158, TaMADS159 | | | | | |
| SEP1 | TaSEP1.4D | HC | TraesCS7D01G120500 | 1 | 1 | OsMADS5-1 | PF00319; PF01486 | + | + | 227 | 26.21 | y | 334.98 | TRIAE_CS42_7DS_TGACv1_621517_AA2017990 | Traes_7DS_68DE33D2A.1 | chr7D | 74600771 | 74607936 | 339.236 | 7165 | R1 | IWGSC | TaMADS173 | TaSEP-6 | TaMADAGL8 | WM20 | AM502886 | DQ512359 |
| SEP1 | TaSEP1.5A | HC | TraesCS7A01G122100 | 1 | 2 | OsMADS5 | PF00319; PF01486 | + | + | 227 | 26.29 | y | 37.09 | TRIAE_CS42_7AS_TGACv1_570718_AA1839760 | Traes_7AS_C25A349A9.1 | chr7A | 78568484 | 78575570 | 50250.667 | 7086 | R1 | IWGSC | TaMADS148 | | TaMADAGL40 | | DQ512370 | |
| SEP1 | TaSEP1.5B1 | HC | TraesCS7B01G020900 | 1 | 1 | OsMADS5 | PF00319; PF01486 | + | + | 193 | 21.91 | y | 6.29 | TRIAE_CS42_7BS_TGACv1_592410_AA1937560 | Traes_7BS_F4CFCDF52.1 | chr7B | 18498249 | 18506298 | 605.534 | 8049 | R2a | FGENESH+ | TaMADS162 | | | | | |
| SEP1 | TaSEP1.5B2 | HC | TraesCS7B01G021100 | 1 | 1 | OsMADS5 | PF00319; PF01486 | + | + | 216 | 25.03 | n | 5.20 | TRIAE_CS42_U_TGACv1_640724_AA2071080 | | chr7B | 19111832 | 19141655 | 71047.784 | 29823 | R2a | FGENESH+ | | | TaMADAGL34 | | | |
| SEP1 | TaSEP1.5D | HC | TraesCS7D01G120600 | 1 | 1 | OsMADS5 | PF00319; PF01486 | + | + | 227 | 26.44 | n | 11.29 | TRIAE_CS42_7DS_TGACv1_623155_AA2050210 | Traes_7DS_D435DF316.1 | chr7D | 74947172 | 74965204 | 53947.038 | 18032 | R1 | IWGSC | TaMADS179 | | | | | |
| SEP1 | TaSEP1.6A | HC | TraesCS5A01G391800 | 1 | 1 | OsMADS34 | PF00319; PF01486 | + | + | 238 | 27.49 | y | 22.83 | TRIAE_CS42_5AS_TGACv1_393214_AA1269860 | Traes_5AS_7BFB385EF.1 | chr5A | 588017766 | 588024080 | 90907.445 | 6314 | R3 | IWGSC | TaMADS103 | TaSEP-5A | TaMADWM19A | WM19A | AM502884 | |
| SEP1 | TaSEP1.6B | HC | TraesCS5B01G396700 | 1 | 1 | OsMADS34 | PF00319; PF01486 | + | + | 236 | 27.29 | y | 134.98 | TRIAE_CS42_5DL_TGACv1_433206_AA1405530 | | chr5B | 467417868 | 467424909 | 4427.309 | 7041 | R3 | IWGSC | | | | | | |
| SEP1 | TaSEP1.6D | HC | TraesCS5D01G401700 | 1 | 1 | OsMADS34 | PF00319; PF01486 | + | + | 236 | 27.22 | y | 19.33 | TRIAE_CS42_5BL_TGACv1_405406_AA1326650 | Traes_5BL_4CA71C036.1 | chr5D | 574055475 | 574061439 | NA | 5964 | R3 | IWGSC | TaMADS106 | | | WM19B | | |
| SEP3 | TaSEP3.1A | HC | TraesCS7A01G260600 | 1 | 2 | OsMADS7 | PF00319; PF01486 | + | + | 248 | 28.74 | y | 424.22 | TRIAE_CS42_7AS_TGACv1_569904_AA1826720 | | chr7A | 253427732 | 253433200 | 956.781 | 5468 | C | IWGSC | TaMADS142 | | | | | |
| SEP3 | TaSEP3.1B | HC | TraesCS7B01G158600 | 1 | 1 | OsMADS7 | PF00319; PF01486 | + | + | 246 | 28.50 | y | 297.55 | TRIAE_CS42_7BS_TGACv1_592319_AA1935630 | | chr7B | 215000979 | 215006859 | 925.268 | 5880 | C | IWGSC | | TaSEP-4 | | WM11 | AM502877 | |
| SEP3 | TaSEP3.1D | HC | TraesCS7D01G261600 | 1 | 2 | OsMADS7 | PF00319; PF01486 | + | + | 248 | 28.75 | y | 323.93 | TRIAE_CS42_7DS_TGACv1_621785_AA2026060 | Traes_7DS_4AF5C695F.1, Traes_7DS_C8 | chr7D | 237609830 | 237619034 | 1354.456 | 9204 | C | IWGSC | TaMADS170, TaMADS178 | | | | | |
| SEP3 | TaSEP3.2A | HC | TraesCS5A01G286800 | 1 | 1 | OsMADS8 | PF00319; PF01486 | + | + | 252 | 29.05 | y | 180.98 | TRIAE_CS42_5AL_TGACv1_374543_AA1202840 | Traes_5AL_EBF32FE10.1 | chr5A | 494862838 | 494871925 | 92539.526 | 9087 | R3 | IWGSC | TaMADS100 | | TaMADWM10A | | AM502875 | |
| SEP3 | TaSEP3.2B | HC | TraesCS5B01G286100 | 1 | 2 | OsMADS8 | PF00319; PF01486 | + | + | 249 | 28.73 | y | 164.88 | TRIAE_CS42_5BL_TGACv1_406187_AA1341980 | Traes_5BL_9627436AE.1 | chr5B | 471852218 | 471860116 | 101942.76 | 7898 | R3 | IWGSC | TaMADS109 | TaSEP-3A | | | AM502875 | |
| SEP3 | TaSEP3.2D | HC | TraesCS5D01G294500 | 1 | 1 | OsMADS8 | PF00319; PF01486 | + | + | 252 | 29.05 | y | 139.26 | TRIAE_CS42_5DL_TGACv1_435792_AA1454980 | Traes_5DL_8C647BFE2.1 | chr5D | 391729992 | 391737307 | 75439.3 | 7315 | R3 | IWGSC | TaMADS113 | TaSEP-3B | TaMADMADS | WM10C | AM502876 | |
| AG/STK | TaAG.1A | HC | TraesCS1A01G125800 | 1 | 2 | OsMADS58 | PF00319; PF01486 | + | + | 269 | 30.38 | y | 184.08 | TRIAE_CS42_1AS_TGACv1_018970_AA0057130 | Traes_1AS_985B833A1.1 | chr1A | 150449685 | 150456774 | 208304.583 | 7089 | R2a | IWGSC | TaMADS8 | | | | | |
| AG/STK | TaAG.1B | HC | TraesCS1D01G127700 | 1 | 2 | OsMADS58 | PF00319; PF01486 | + | + | 269 | 30.43 | y | 182.84 | TRIAE_CS42_1DS_TGACv1_080255_AA0244360 | Traes_1DS_F22A3DB6A.1 | chr1D | 140519067 | 140526895 | 146929.95 | 7828 | C | IWGSC | TaMADS22 | TaAG-1 | | WM2 | AM502863 | |
| AG/STK | TaAG.1D | HC | TraesCS1B01G144800 | 1 | 3 | OsMADS58 | PF00319; PF01486 | + | + | 269 | 30.42 | y | 198.92 | TRIAE_CS42_1BS_TGACv1_049537_AA0156150 | Traes_1BS_1202C8C0D.1, Traes_1BS_40F | chr1B | 198145270 | 198154010 | 191058.922 | 8740 | C | IWGSC | TaMADS12,TaMADS13 | | | | | |
| AG/STK | TaAG.2A | HC | TraesCS3A01G314300 | 1 | 3 | OsMADS3 | PF00319; PF01486 | + | + | 273 | 30.92 | y | 123.42 | TRIAE_CS42_3AS_TGACv1_211344_AA0688860 | Traes_3AS_55E9080C2.1, Traes_4AS_A3 | chr3A | 555035146 | 555042843 | 96539.472 | 7697 | R2b | IWGSC | TaMADS50, TaMADS77 | TaAG-2A | TaMADAGL39 | WM29A | AM502898 | |
| AG/STK | TaAG.2B | HC | TraesCS3B01G157500 | 1 | 4 | OsMADS3 | PF00319; PF01486 | + | + | 276 | 31.37 | y | 139.04 | TRIAE_CS42_3B_TGACv1_220593_AA0710240 | | chr3B | 151443942 | 151451682 | 361802.32 | 7740 | R2a | IWGSC | TaMADS63 | TaAG-2B | | WM29B | AM502899 | |
| AG/STK | TaAG.2D | HC | TraesCS3D01G140200 | 1 | 4 | OsMADS3 | PF00319; PF01486 | + | + | 273 | 31.04 | y | 56.85 | TRIAE_CS42_3DS_TGACv1_272039_AA0913620 | Traes_3DS_51A589227.1 | chr3D | 100852964 | 100860875 | 292663.6 | 7911 | R2a | IWGSC | TaMADS71 | | | | | |
| AG/STK | TaSTK.1A | HC | TraesCS5A01G117500 | 1 | 2 | OsMADS13 | PF00319; PF01486 | + | + | 254 | 28.51 | y | 72.13 | TRIAE_CS42_5AS_TGACv1_393214_AA1269860 | | chr5A | 239118848 | 239124243 | 255738.595 | 5395 | C | IWGSC | | TaAG-3B | TaMADAGL2 | WM3B | AM502865 | |
| AG/STK | TaSTK.1B | HC | TraesCS5B01G115100 | 1 | 2 | OsMADS13 | PF00319; PF01486 | + | + | 252 | 28.40 | y | 43.36 | TRIAE_CS42_5BS_TGACv1_423180_AA1369410 | Traes_5BS_A19FD8E34.1 | chr5B | 191029558 | 191038231 | 276379.637 | 8673 | C | IWGSC | TaMADS111 | | | | | |
| AG/STK | TaSTK.1D | HC | TraesCS5D01G118200 | 1 | 2 | OsMADS13 | PF00319; PF01486 | + | + | 252 | 28.26 | y | 119.79 | TRIAE_CS42_5DS_TGACv1_457116_AA1482580 | Traes_5DS_16243E52C.1 | chr5D | 156574030 | 156579651 | 235150.341 | 5621 | C | IWGSC | TaMADS115 | TaAG-3A | | WM3A | AM502864 | DQ512364 |
| AG/STK | TaSTK.2A | HC | TraesCS1A01G262700 | 1 | 2 | OsMADS21 | PF00319; PF01486 | + | + | 275 | 30.30 | y | 28.56 | TRIAE_CS42_1AL_TGACv1_002892_AA0045960 | Traes_1AL_5F5A87122.1 | chr1A | 458216619 | 458220982 | 1725.858 | 4363 | R2b | IWGSC | TaMADS2 | TaAG-4A | TaMADWM27A | WM27A | AM502894 | |
| AG/STK | TaSTK.2B | HC | TraesCS1B01G273300 | 1 | 1 | OsMADS21 | PF00319; PF01486 | + | + | 255 | 28.29 | y | 26.12 | TRIAE_CS42_1BL_TGACv1_033317_AA0138690 | Traes_1BL_B44C0D37C.1 | chr1B | 480437844 | 480442278 | 2008.406 | 4434 | R2b | IWGSC | TaMADS9 | TaAG-4B | TaMADWM27B | WM27B | AM502895 | |
| AG/STK | TaSTK.2D | HC | TraesCS1D01G262700 | 1 | 1 | OsMADS21 | PF00319; PF01486 | + | + | 250 | 27.65 | y | 33.44 | TRIAE_CS42_1DL_TGACv1_063298_AA0226000 | Traes_1DL_6DA0DFC5B.1 | chr1D | 356928483 | 356933138 | 3453.44 | 4655 | R2b | IWGSC | TaMADS17 | | | | | |
| AGL12 | TaAGL12.1A | HC | TraesCS7A01G319400 | 1 | 2 | OsMADS26 | PF00319; PF01486 | + | + | 224 | 25.22 | y | 9.52 | TRIAE_CS42_7AL_TGACv1_557712_AA1785060 | | chr7A | 459626073 | 459628590 | 99571.149 | 2517 | R2b | IWGSC | | | | | | |
| AGL12 | TaAGL12.1B | HC | TraesCS7B01G220200 | 1 | 2 | OsMADS26 | PF00319; PF01486 | + | + | 229 | 25.80 | y | 60.67 | TRIAE_CS42_7BL_TGACv1_579758_AA1909910 | Traes_7BL_9BCF391CF.1 | chr7B | 411493387 | 411495817 | 110577.229 | 2430 | C | IWGSC | TaMADS153 | | TaMADAGL32 | | | |
| AGL12 | TaAGL12.1D | HC | TraesCS7D01G315900 | 1 | 3 | OsMADS26 | PF00319; PF01486 | + | + | 232 | 26.18 | y | 56.58 | TRIAE_CS42_7DL_TGACv1_604914_AA2002950 | Traes_7DL_CAF83263E.1 | chr7D | 401836566 | 401839012 | 91005.033 | 2446 | R2b | IWGSC | TaMADS166 | TaAGL12-1 | | WM12 | AM502878 | |
| AGL12 | TaAGL12.2A | HC | TraesCS2A01G311100 | 1 | 1 | OsMADS33 | PF00319; PF01486 | + | + | 202 | 23.40 | y | 20.67 | TRIAE_CS42_2AL_TGACv1_094081_AA0292300 | | chr2A | 534625654 | 534630242 | 37424.556 | 4588 | R2b | IWGSC | | | | | | |
| AGL12 | TaAGL12.2B | HC | TraesCS2B01G328000 | 1 | 1 | OsMADS33 | PF00319; PF01486 | + | + | 202 | 23.52 | y | 8.31 | TRIAE_CS42_2BL_TGACv1_129629_AA0391000 | Traes_2BL_E0978B1BC.1 | chr2B | 470059642 | 470062563 | 20256.506 | 2921 | R2b | IWGSC | TaMADS35 | | | | | |
| AGL12 | TaAGL12.2D | HC | TraesCS2D01G309300 | 1 | 1 | OsMADS33 | PF00319; PF01486 | + | + | 202 | 23.40 | y | 3.73 | TRIAE_CS42_2DL_TGACv1_161097_AA0556680 | Traes_2DL_71F120931.1 | chr2D | 396662325 | 396667410 | 21376.997 | 5085 | R2b | IWGSC | TaMADS41 | | | | | |
| AGL17 | TaAGL17.10B | HC | TraesCS5B01G002500 | 1 | 1 | none | PF00319 | + | - | 151 | 17.41 | n | 0.95 | TRIAE_CS42_5BS_TGACv1_423187_AA1369730 | Traes_5BS_284476236.1 | chr5B | 3866145 | 3867846 | 187161.712 | 1701 | R1 | FGENESH+ | TaMADS110 | | | | | |
| AGL17 | TaAGL17.10D | HC | TraesCS5D01G002100 | 1 | 1 | none | PF00319; PF01486 | + | + | 225 | 26.09 | n | 0.82 | TRIAE_CS42_5DS_TGACv1_457275_AA1484570 | | chr5D | 2286406 | 2288880 | 2.806 | 2474 | R1 | IWGSC | | | | | | |
| AGL17 | TaAGL17.10U | LC | TraesCSU01G389400LC | 1 | 1 | none | PF00319; PF01486 | + | + | 160 | 18.59 | y | 11.33 | TRIAE_CS42_5AS_TGACv1_394239_AA1279100 | Traes_5AS_E9E60BA43.1 | chrUn | 300170542 | 300171819 | NA | 1277 | NA | IWGSC | TaMADS105 | | | | | |
| AGL17 | TaAGL17.11A1 | LC | TraesCS5A01G001400LC | 1 | 1 | none | PF00319 | + | - | 92 | 10.31 | n | 0.22 | | | chr5A | 1297025 | 1297304 | 5.339 | 279 | R1 | IWGSC | | | | | | |
| AGL17 | TaAGL17.11A2 | NA | no gene predicted at site | 1 | 1 | none | PF00319; PF01486 | + | + | 158 | 18.35 | n | NA | | | chr7A | 68342914 | 68344465 | 396.003 | 1551 | R1 | FGENESH+ | | | | | | |
| AGL17 | TaAGL17.11A3 | LC | TraesCS7A01G151800LC | 1 | 1 | none | PF00319 | + | - | 126 | 14.25 | n | 0.14 | TRIAE_CS42_7AS_TGACv1_570873_AA1841890 | Traes_7AS_EFE436F1D.1 | chr7A | 68762176 | 68770437 | 1.785 | 8261 | R1 | IWGSC | TaMADS150 | | | | | |
| AGL17 | TaAGL17.11A4 | NA | no gene predicted at site | 1 | 1 | none | PF00319; PF01486 | + | + | 176 | 20.69 | n | NA | | | chr7A | 68772222 | 68779045 | 9732.946 | 6823 | R1 | FGENESH+ | | | | | | |
| AGL17 | TaAGL17.11B1 | HC | TraesCS7B01G009600 | 1 | 1 | none | PF00319 | + | - | 67 | 7.55 | n | 0.24 | | | chr7B | 6002981 | 6003351 | 62.782 | 370 | R1 | IWGSC | | | | | | |
| AGL17 | TaAGL17.11B2 | HC | TraesCS7B01G009900 | 1 | 2 | none | PF00319; PF01486 | + | + | 187 | 21.90 | y | 0.18 | TRIAE_CS42_7BS_TGACv1_593230_AA1949580 | Traes_7BS_209950516.1 | chr7B | 6081190 | 6088818 | 12119.172 | 7628 | R1 | IWGSC | TaMADS156 | | | | | |
| AGL17 | TaAGL17.11D1 | LC | TraesCS7D01G135600LC | 1 | 1 | none | PF00319 | + | - | 339 | 36.81 | | 0.00 | TRIAE_CS42_7DS_TGACv1_621619_AA2021270 | Traes_7DS_D9008CC09.1 | chr7D | 64450486 | 64453017 | 18.466 | 2531 | R1 | IWGSC | TaMADS180 | | | | | |
| AGL17 | TaAGL17.11D2 | HC | TraesCS7D01G107000 | 1 | 1 | none | PF00319 | + | - | 71 | 8.19 | n | 0.00 | TRIAE_CS42_7DS_TGACv1_621619_AA2021290 | | chr7D | 64496913 | 64497302 | 18.043 | 389 | R1 | IWGSC | | | | | | |
| AGL17 | TaAGL17.11D3 | NA | no gene predicted at site | 1 | 1 | none | PF00319 | + | - | 150 | 17.05 | n | NA | | | chr7D | 64490707 | 64494277 | 2.636 | 3570 | R1 | FGENESH+ | | | | | | |
| AGL17 | TaAGL17.11D4 | NA | no gene predicted at site | 1 | 1 | none | PF00319; PF01486 | + | + | 313 | 36.58 | n | NA | | | chr7D | 64531588 | 64533657 | 10067.114 | 2069 | R1 | FGENESH+ | TaMADS168 | | | | | |
| AGL17 | TaAGL17.1A | HC | TraesCS2A01G337900 | 1 | 1 | OsMADS61 | PF00319; PF01486 | + | + | 240 | 27.52 | y | 14.20 | TRIAE_CS42_2AL_TGACv1_093384_AA0278940 | Traes_2AL_267502097.1 | chr2A | 572054798 | 572068820 | 105412.955 | 14022 | R2b | IWGSC | TaMADS26 | TaAGL17-1 | TaMADWM30 | WM30 | AM502900 | |
| AGL17 | TaAGL17.1B | HC | TraesCS2B01G344000 | 1 | 1 | OsMADS61 | PF00319; PF01486 | + | + | 240 | 27.49 | y | 12.80 | TRIAE_CS42_2BL_TGACv1_130458_AA0411830 | Traes_2BL_3E613DE21.1 | chr2B | 490319069 | 490333877 | 138614.981 | 14808 | R2b | IWGSC | TaMADS32 | | | | | |
| AGL17 | TaAGL17.1D | HC | TraesCS2D01G325000 | 1 | 1 | OsMADS61 | PF00319; PF01486 | + | + | 240 | 27.50 | y | 12.05 | TRIAE_CS42_2DL_TGACv1_159992_AA0545320 | | chr2D | 418044407 | 418060526 | 115001.507 | 16119 | R2b | IWGSC | TaMADS38 | | | | | |
| AGL17 | TaAGL17.2A1 | HC | TraesCSU01G209900 | 1 | 1 | OsMADS25-1 | PF00319; (PF01486)* | + | (-) | 232 | 26.47 | y | 3.13 | TRIAE_CS42_5AS_TGACv1_394239_AA1279110 | Traes_5AS_029A65B0A.1 | chrUn | 311790493 | 311791081 | NA | 588 | NA | TGAC | TaMADS101 | TaAGL17-2A | TaMADWM31A | WM31A | AM502901 | |
| AGL17 | TaAGL17.2A2 | HC | TraesCS5A01G001800 | 1 | 1 | OsMADS25-1 | PF00319 | + | - | 59 | 6.65 | n | 0.84 | | | chr5A | 1302643 | 1302822 | 237816.026 | 179 | R1 | IWGSC | | | | | | |
| AGL17 | TaAGL17.2A3 | LC | TraesCS5A01G001300LC | 1 | 1 | OsMADS25-1 | PF01486 | - | + | 148 | 17.06 | y | 2.40 | TRIAE_CS42_5AS_TGACv1_394239_AA1279110 | Traes_1BS_B03D4CD04.1, Traes_3DS_B0 | chr5A | 1266561 | 1268289 | 28.736 | 1728 | R1 | IWGSC | TaMADS15, TaMADS73, TaMADS104, TaMADS133 | | | | | |
| AGL17 | TaAGL17.2B | HC | TraesCS5B01G002400 | 1 | 1 | OsMADS25-1 | PF00319; PF01486 | + | + | 232 | 26.38 | y | 1.05 | TRIAE_CS42_5BS_TGACv1_423187_AA1369720 | Traes_5BS_BC15EF87A.1 | chr5B | 3849483 | 3862668 | 3.477 | 13185 | R1 | IWGSC | TaMADS112 | TaAGL17-2B | TaMADWM31B | WM31B | AM502902 | |
| AGL17 | TaAGL17.2D | HC | TraesCS5D01G002200 | 1 | 2 | OsMADS25-1 | PF00319; PF01486 | + | + | 190 | 21.88 | y | 1.42 | TRIAE_CS42_5DS_TGACv1_457275_AA1484560 | Traes_5DS_3E8E121C7.1 | chr5D | 2291686 | 2303227 | 154270.803 | 11541 | R1 | IWGSC | TaMADS116 | | | | | |
| AGL17 | TaAGL17.3A | HC | TraesCS6A01G292300 | 1 | 2 | OsMADS57 | PF00319; PF01486 | + | + | 235 | 26.48 | y | 2.31 | TRIAE_CS42_6AL_TGACv1_471339_AA1507320 | Traes_6AL_B5D4C3A49.1 | chr6A | 524304592 | 524312866 | 26324.245 | 8274 | R2b | IWGSC | TaMADS121 | TaAGL17-3A | TaMADWM32B | WM32A | AM502904, AM502905 | |
| AGL17 | TaAGL17.3B | HC | TraesCS6B01G322700 | 1 | 2 | OsMADS57 | PF00319; PF01486 | + | + | 235 | 26.42 | y | 6.30 | TRIAE_CS42_6BL_TGACv1_500233_AA1601850 | | chr6B | 571729444 | 571737072 | 34058.05 | 7628 | R2b | IWGSC | | TaAGL17-3B | | WM32B | AM502905 | |
| AGL17 | TaAGL17.3D | HC | TraesCS6D01G273500 | 1 | 2 | OsMADS57 | PF00319; PF01486 | + | + | 235 | 26.46 | y | 15.05 | TRIAE_CS42_6DL_TGACv1_527898_AA1709630 | Traes_6DL_D1C1DBD34.1 | chr6D | 382344991 | 382353112 | 21616.618 | 8121 | R2b | IWGSC | TaMADS134 | | | | | |
| AGL17 | TaAGL17.4A | HC | TraesCS7A01G111700 | 1 | 2 | OsMADS25-2 | PF00319; PF01486 | + | + | 238 | 27.01 | y | 0.65 | TRIAE_CS42_7AS_TGACv1_570106_AA1830540 | Traes_7AS_8123257BA.1 | chr7A | 68227268 | 68236661 | 87.995 | 9393 | R1 | IWGSC | TaMADS145 | | | | | |
| AGL17 | TaAGL17.4B | HC | TraesCS7B01G009500 | 1 | 1 | OsMADS25-2 | PF00319; PF01486 | + | + | 230 | 26.09 | y | 0.44 | TRIAE_CS42_7BS_TGACv1_592058_AA1929510 | Traes_7BS_DC9822CEF.1 | chr7B | 5981410 | 5999811 | 3.17 | 18401 | R1 | IWGSC | TaMADS161 | | | | | |
| AGL17 | TaAGL17.4D | HC | TraesCS7D01G106600 | 1 | 1 | OsMADS25-2 | PF00319; PF01486 | + | + | 230 | 26.12 | y | 8.67 | TRIAE_CS42_7DS_TGACv1_621619_AA2021270 | Traes_7DS_7A30E1397.1 | chr7D | 64436534 | 64446655 | 3.831 | 10121 | R1 | IWGSC | TaMADS174 | TaAGL17-2C | TaMADWM31C | WM31C | AM502903 | |
| AGL17 | TaAGL17.5A1 | HC | TraesCS7A01G111800 | 1 | 1 | OsMADS25-3 | PF00319; PF01486 | + | + | 230 | 26.22 | y | 0.08 | TRIAE_CS42_7AS_TGACv1_570454_AA1835900 | Traes_7AS_360247894.1 | chr7A | 68324656 | 68338010 | 4.904 | 13354 | R1 | IWGSC | TaMADS143 | | | | | |
| AGL17 | TaAGL17.5A2 | HC | TraesCS7A01G111900 | 1 | 1 | OsMADS25-3 | PF00319; PF01486 | + | + | 230 | 25.95 | y | 1.21 | TRIAE_CS42_7AS_TGACv1_570873_AA1841880 | Traes_2DL_6A10DD109.1, Traes_7AS_37 | chr7A | 68740468 | 68762474 | 0 | 22006 | R1 | IWGSC | TaMADS39, TaMADS144, TaMADS147 | | | | | |
| AGL17 | TaAGL17.5B | HC | TraesCS7B01G009800 | 1 | 1 | OsMADS25-3 | PF00319; PF01486 | + | + | 230 | 25.92 | y | 0.16 | TRIAE_CS42_7BS_TGACv1_593230_AA1949590 | Traes_7BS_9D42F9BEA.1 | chr7B | 6066133 | 6078759 | 2.431 | 12626 | R1 | IWGSC | TaMADS160 | | | | | |
| AGL17 | TaAGL17.5D1 | HC | TraesCS7D01G106900 | 1 | 1 | OsMADS25-3 | PF00319; PF01486 | + | + | 230 | 25.99 | y | 1.44 | TRIAE_CS42_7DS_TGACv1_621619_AA2021300 | Traes_7DS_59CBF5647.1 | chr7D | 64471483 | 64488459 | 2.248 | 16976 | R1 | IWGSC | TaMADS172 | | | | | |
| AGL17 | TaAGL17.5D2 | HC | TraesCS7D01G107200 | 1 | 1 | OsMADS25-3 | PF00319; PF01486 | + | + | 197 | 22.56 | n | 0.15 | TRIAE_CS42_7DS_TGACv1_621619_AA2021310 | Traes_7DS_7F8C88C92.1 | chr7D | 64515345 | 64528197 | 3.391 | 12852 | R1 | FGENESH+ | TaMADS176 | | TaMADAGL6 | | DQ512329 | |
| AGL17 | TaAGL17.6B1 | NA | no gene predicted at site | 1 | 1 | OsMADS59-1 | PF00319; PF01486 | + | + | 115 | 13.19 | n | NA | | | chr7B | 731976814 | 731982042 | 56.992 | 5228 | R3 | FGENESH+ | | | | | | |
| AGL17 | TaAGL17.6B2 | LC | TraesCS7B01G793900LC | 1 | 1 | OsMADS59-1 | PF00319 | + | - | 109 | 12.58 | n | 0.13 | | | chr7B | 732039034 | 732043532 | 56.788 | 4498 | R3 | IWGSC | | | | | | |
| AGL17 | TaAGL17.6B3 | LC | TraesCS7B01G794200LC | 1 | 1 | OsMADS59-1 | PF00319; PF01486 | + | + | 232 | 26.58 | y | 2.76 | | | chr7B | 732100320 | 732107408 | NA | 7088 | R3 | IWGSC | | | | | | |
| AGL17 | TaAGL17.6U1 | LC | TraesCSU01G169600LC | 1 | 1 | OsMADS59-1 | PF00319; PF01486 | + | + | 211 | 24.35 | n | 1.15 | NA | Traes_7BL_F5B6736D0.1 | chrUn | 88448060 | 88454722 | NA | 6662 | NA | IWGSC | TaMADS154 | | | | | |
| AGL17 | TaAGL17.6U2 | LC | TraesCSU01G236400LC | 1 | 1 | OsMADS59-1 | PF00319 | + | - | 97 | 11.08 | n | 0.98 | | | chrUn | 160334913 | 160335206 | NA | 293 | NA | IWGSC | | | | | | |
| AGL17 | TaAGL17.6U3 | LC | TraesCSU01G236600LC | 1 | 1 | OsMADS59-1 | PF00319 | + | - | 109 | 12.46 | y | 2.67 | TRIAE_CS42_7BL_TGACv1_584691_AA1919350 | Traes_4AL_72A03AD23.1 | chrUn | 160396770 | 160401475 | NA | 4705 | NA | IWGSC | TaMADS74 | | | | | |
| AGL17 | TaAGL17.7B1 | HC | TraesCS1B01G058200 | 1 | 1 | OsMADS59-2 | PF00319; PF01486 | + | + | 223 | 25.62 | n | 0.15 | TRIAE_CS42_1BS_TGACv1_049691_AA0159690 | | chr1B | 40330406 | 40347378 | 8.306 | 16972 | R1 | IWGSC | | | | | | |
| AGL17 | TaAGL17.7B2 | NA | no gene predicted at site | 1 | 1 | OsMADS59-2 | PF00319 | + | - | 168 | 18.73 | n | NA | TRIAE_CS42_1BS_TGACv1_049691_AA0159690 | | chr1B | 40355684 | 40362159 | 24.514 | 6475 | R1 | FGENESH+ | | | | | | |
| AGL17 | TaAGL17.7D | HC | TraesCS1D01G045600 | 1 | 1 | OsMADS59-2 | PF00319; PF01486 | + | + | 234 | 26.82 | n | 0.45 | TRIAE_CS42_1DS_TGACv1_080122_AA0240280 | | chr1D | 24929697 | 24936216 | 165.079 | 6519 | R1 | IWGSC | | | | | | |
| AGL17 | TaAGL17.8A | HC | TraesCS1A01G045000 | 1 | 1 | OsMADS59-3 | PF00319; PF01486 | + | + | 198 | 22.84 | n | 0.44 | TRIAE_CS42_1AS_TGACv1_019741_AA0070900 | Traes_1AS_24511D656.1 | chr1A | 25791227 | 25795729 | 124653.956 | 4502 | R1 | IWGSC | TaMADS6 | | | | | |
| AGL17 | TaAGL17.8B | HC | TraesCS1B01G058300 | 1 | 1 | OsMADS59-3 | PF00319; PF01486 | + | + | 236 | 26.52 | y | 1.14 | TRIAE_CS42_1BS_TGACv1_049691_AA0159710 | | chr1B | 40386673 | 40391043 | 157754.227 | 4370 | R1 | IWGSC | | | | | | |
| AGL17 | TaAGL17.8D | HC | TraesCS1D01G045700 | 1 | 1 | OsMADS59-3 | PF00319; PF01486 | + | + | 236 | 26.59 | y | 2.15 | TRIAE_CS42_U_TGACv1_642436_AA2117570 | Traes_1DS_A0312C264.1 | chr1D | 25101295 | 25105690 | 115413.377 | 4395 | R1 | IWGSC | TaMADS21 | | | | | |
| AGL17 | TaAGL17.9A | HC | TraesCS1A01G044900 | 1 | 1 | OsMADS59-4 | PF00319; PF01486 | + | + | 243 | 27.70 | n | 0.14 | TRIAE_CS42_1AS_TGACv1_020392_AA0077140 | Traes_1AS_3232D5589.1 | chr1A | 25710162 | 25714282 | 76.945 | 4120 | R1 | IWGSC | TaMADS7 | | | | | |
| AGL17 | TaAGL17.9B | HC | TraesCS1B01G058100 | 1 | 1 | OsMADS59-4 | PF00319; PF01486 | + | + | 243 | 27.67 | y | 7.35 | TRIAE_CS42_1BS_TGACv1_050417_AA0171830 | Traes_1BS_7366111C0.1 | chr1B | 40195364 | 40199852 | 130.554 | 4488 | R1 | IWGSC | TaMADS14 | | | | | |
| AGL17 | TaAGL17.9D | HC | TraesCS1D01G045500 | 1 | 1 | OsMADS59-4 | PF00319; PF01486 | + | + | 233 | 26.72 | y | 8.92 | TRIAE_CS42_1DS_TGACv1_080895_AA0255360 | Traes_1DS_F70AAB507.1 | chr1D | 24787220 | 24791488 | 138.209 | 4268 | R1 | IWGSC | TaMADS23 | | | | | |
| AGL6 | TaAGL6A | HC | TraesCS6A01G259000 | 1 | 3 | OsMADS6/Os | PF00319; PF01486 | + | + | 259 | 29.47 | y | 54.21 | TRIAE_CS42_U_TGACv1_641552_AA2097800 | Traes_6AL_1F7DAC5FA.1 | chr6A | 481736953 | 481744180 | 42560.412 | 7227 | R2b | IWGSC | TaMADS119 | TaAGL6-1A | | WM9A | AM502872 | TaMADS#12 |
| AGL6 | TaAGL6B | HC | TraesCS6B01G286400 | 1 | 3 | OsMADS6/Os | PF00319; PF01486 | + | + | 259 | 29.40 | y | 34.95 | TRIAE_CS42_6BL_TGACv1_500609_AA1607090 | Traes_6BL_7C6B17284.1 | chr6B | 516074914 | 516082298 | 55647.146 | 7384 | R2b | IWGSC | TaMADS126 | TaAGL6-1B | TaMADS12 | WM9B | AM502873 | |
| AGL6 | TaAGL6D | HC | TraesCS6D01G240200 | 1 | 3 | OsMADS6/Os | PF00319; PF01486 | + | + | 260 | 29.57 | y | 45.01 | TRIAE_CS42_6DL_TGACv1_526443_AA1683650 | Traes_6DL_5B1D4DBF5.1 | chr6D | 341645092 | 341652716 | 40692.275 | 7624 | R2b | IWGSC | TaMADS131 | TaAGL6-1C | | WM9D | AM502874 | |
| AP1 | TaAP1.1A | HC | TraesCS5A01G391700 | 1 | 2 | OsMADS14 | PF00319; PF01486 | + | + | 244 | 27.85 | y | 129.15 | TRIAE_CS42_5DS_TGACv1_457465_AA1486810 | Traes_5AL_13E2DEC48.1, Traes_5DS_86 | chr5A | 587411451 | 587423425 | 594.341 | 11974 | R3 | IWGSC | TaMADS98, TaMADS117 | TaAP1-1 | | WM6 | AM502869 | |
| AP1 | TaAP1.1B | HC | TraesCS5B01G396600 | 1 | 2 | OsMADS14 | PF00319; PF01486 | + | + | 244 | 27.84 | y | 22.18 | TRIAE_CS42_5BL_TGACv1_405351_AA1325650 | Traes_5BL_5D2D22E67.1, Traes_5BL_89 | chr5B | 573802876 | 573816110 | NA | 13234 | R3 | IWGSC | TaMADS107, TaMADS108 | | | | | |
| AP1 | TaAP1.1D | HC | TraesCS5D01G401500 | 1 | 3 | OsMADS14 | PF00319; PF01486 | + | + | 254 | 28.78 | y | 50.23 | TRIAE_CS42_5DL_TGACv1_434261_AA1432830 | Traes_5DL_9CC4EC839.1 | chr5D | 467176607 | 467184508 | 106870.967 | 7901 | R3 | IWGSC | TaMADS114 | | | | | |
| AP1 | TaAP1.2A | HC | TraesCS2A01G174300 | 3 | 3 | OsMADS18 | PF00319; PF01486 | + | + | 267 | 30.06 | y | 141.88 | TRIAE_CS42_2AS_TGACv1_114196_AA0364910 | Traes_2AS_E2C631DBE.1, Traes_2AS_F7 | chr2A | 132859446 | 132876629 | 279066.563 | 17183 | R2a | IWGSC | TaMADS29, TaMADS30, TaMADS36 | | | WM7 | | |
| AP1 | TaAP1.2B | HC | TraesCS2B01G200800 | 1 | 1 | OsMADS18 | PF00319; PF01486 | + | + | 267 | 30.01 | y | 124.09 | TRIAE_CS42_2BS_TGACv1_148746_AA0494980 | | chr2B | 180040807 | 180060077 | 207638.399 | 19270 | R2a | IWGSC | | TaAP1-2 | TaMADAGL10 | WM7 | AM502870 | TaMADS2 |
| AP1 | TaAP1.2D | HC | TraesCS2D01G181400 | 3 | 3 | OsMADS18 | PF00319; PF01486 | + | + | 267 | 30.03 | n | 132.43 | TRIAE_CS42_2DS_TGACv1_177164_AA0567240 | Traes_2DS_4F6BA4A13.1, Traes_2DS_F2 | chr2D | 126124904 | 126149761 | 192760.495 | 24857 | R2a | IWGSC | TaMADS43, TaMADS44 | | | | | |
| AP1 | TaAP1.3A | HC | TraesCS2A01G261200 | 1 | 2 | OsMADS15 | PF00319; PF01486 | + | + | 273 | 31.22 | y | 154.83 | TRIAE_CS42_2AL_TGACv1_092934_AA0267480 | Traes_2AL_20C2D79E1.1 | chr2A | 411943192 | 411950517 | 122675.137 | 7325 | R2b | IWGSC | TaMADS25 | | | | | |
| AP1 | TaAP1.3B | HC | TraesCS2B01G281000 | 1 | 1 | OsMADS15 | PF00319; PF01486 | + | + | 274 | 31.32 | y | 162.85 | TRIAE_CS42_2BL_TGACv1_130188_AA0405830 | Traes_2BL_26F24E716.1 | chr2B | 387698476 | 387705714 | 82353.928 | 7238 | C | IWGSC | TaMADS31 | TaAP1-3 | TaMADAGL29 | WM8 | AM502871 | |
| AP1 | TaAP1.3D | HC | TraesCS2D01G262700 | 1 | 2 | OsMADS15 | PF00319; PF01486 | + | + | 271 | 31.01 | y | 107.31 | TRIAE_CS42_2DL_TGACv1_158019_AA0506180 | Traes_2DL_903A29CBA.1 | chr2D | 318910256 | 318917796 | 77744.529 | 7540 | C | IWGSC | TaMADS42 | | | | | |
| AP3 | TaAP3.1A | HC | TraesCS7A01G383800 | 1 | 1 | OsMADS16 | PF00319; PF01486 | + | + | 237 | 26.73 | y | 169.42 | TRIAE_CS42_U_TGACv1_640856_AA2077490 | Traes_7AL_1C76E543C.1 | chr7A | 559199739 | 559206787 | 110968.362 | 7048 | R2b | IWGSC | TaMADS140 | | | | | |
| AP3 | TaAP3.1B | HC | TraesCS7B01G286600 | 1 | 2 | OsMADS16 | PF00319; PF01486 | + | + | 227 | 25.71 | y | 142.78 | TRIAE_CS42_7BL_TGACv1_578096_AA1889410 | Traes_7BL_7F4124E70.1, Traes_7BL_F62 | chr7B | 522073046 | 522079520 | 119605.228 | 6474 | R2b | IWGSC | TaMADS152, TaMADS155 | | TaMADS51 | WM13 | | |
| AP3 | TaAP3.1D | HC | TraesCS7D01G380300 | 1 | 1 | OsMADS16 | PF00319; PF01486 | + | + | 227 | 25.71 | y | 126.62 | TRIAE_CS42_7DL_TGACv1_603230_AA1978820 | Traes_7DL_DDCC09B24.1 | chr7D | 492844045 | 492850569 | 85285.357 | 6524 | R2b | IWGSC | TaMADS167 | TaAP3-1 | | WM13 | AM502879 | |
| AP3 | TaAP3.2A | HC | TraesCS6A01G131100 | 1 | 1 | OsMADS16-2 | PF00319; PF01486 | + | + | 239 | 27.13 | n | 0.73 | TRIAE_CS42_6AS_TGACv1_486056_AA1556010 | Traes_6AS_9AA76345D.1 | chr6A | 103268581 | 103270379 | 44977.59 | 1798 | R2a | IWGSC | TaMADS123 | | | | | |
| AP3 | TaAP3.2B | HC | TraesCS6B01G159400 | 1 | 1 | OsMADS16-2 | PF00319; PF01486 | + | + | 231 | 26.16 | n | 0.23 | TRIAE_CS42_6BS_TGACv1_514454_AA1659920 | Traes_6BS_8F1EC63B9.1 | chr6B | 162566837 | 162569327 | 50398.461 | 2490 | R2a | IWGSC | TaMADS130 | | | | | |
| AP3 | TaAP3.2D | HC | TraesCS6D01G120900 | 1 | 1 | OsMADS16-2 | PF00319; PF01486 | + | + | 239 | 27.16 | n | 0.33 | TRIAE_CS42_6DS_TGACv1_543151_AA1736280 | | chr6D | 85911043 | 85913195 | 34081.907 | 2152 | R2a | IWGSC | | | | | | |
| Bsister | TaBS.1A | HC | TraesCS6A01G158100 | 1 | 1 | OsMADS29 | PF00319; PF01486 | + | + | 252 | 28.26 | y | 20.9 | TRIAE_CS42_6AS_TGACv1_485254_AA1541650 | Traes_6AS_57E50EE92.1, Traes_6AS_B4 | chr6A | 148247969 | 148251460 | 159641.755 | 3491 | R2a | IWGSC | TaMADS122, TaMADS124 | | | | | |
| Bsister | TaBS.1B | HC | TraesCS6B01G186700 | 1 | 1 | OsMADS29 | PF00319; PF01486 | + | + | 253 | 28.58 | y | 17.98 | TRIAE_CS42_6BS_TGACv1_514879_AA1665310 | Traes_6BS_5789476CB.1 | chr6B | 212967788 | 212970624 | 303104.29 | 2836 | R2a | IWGSC | TaMADS129 | TaGGM13 | | WM25 | AM502893 | |
| Bsister | TaBS.1D | HC | TraesCS6D01G147400 | 1 | 1 | OsMADS29 | PF00319; PF01486 | + | + | 252 | 28.26 | y | 41.78 | TRIAE_CS42_6DS_TGACv1_544258_AA1747470 | Traes_6DS_9BBDCC9F7.1 | chr6D | 119995102 | 119998077 | 221647.015 | 2975 | R2a | IWGSC | TaMADS138 | | | | | |
| Bsister | TaBS.2A | HC | TraesCS2A01G422400 | 1 | 2 | OsMADS31 | PF00319; PF01486 | + | + | 240 | 27.91 | y | 4.18 | TRIAE_CS42_2AL_TGACv1_096227_AA0317980 | Traes_2AL_8DF89AA72.1 | chr2A | 677481775 | 677484114 | 38341.194 | 2339 | R3 | IWGSC | TaMADS28 | | | | | |
| Bsister | TaBS.2B | HC | TraesCS2B01G440900 | 1 | 1 | OsMADS31 | PF00319; PF01486 | + | + | 253 | 29.33 | y | 2.21 | TRIAE_CS42_2BL_TGACv1_129478_AA0385490 | | chr2B | 632623323 | 632626011 | NA | 2688 | R2b | IWGSC | | | | | | |
| Bsister | TaBS.2D | HC | TraesCS2D01G418800 | 1 | 2 | OsMADS31 | PF00319; PF01486 | + | + | 240 | 27.89 | y | 7.69 | TRIAE_CS42_2DL_TGACv1_159022_AA0530840 | Traes_2DL_6CD5A5CD9.1 | chr2D | 533062033 | 533064922 | 109285.544 | 2889 | R3 | IWGSC | TaMADS40 | | | | | |
| Bsister | TaBS.3A | LC | TraesCS7A01G672200LC | 1 | 1 | OsMADS30-1 | PF00319; PF01486 | + | + | 256 | 29.13 | y | 0.41 | TRIAE_CS42_2BS_TGACv1_147496_AA0483920 | | chr7A | 670566150 | 670568642 | NA | 2492 | R3 | IWGSC | | | | | | |
| Bsister | TaBS.3B | HC | TraesCS7B01G377700 | 1 | 1 | OsMADS30-1 | PF00319; PF01486 | + | + | 226 | 25.41 | y | 2.91 | TRIAE_CS42_7BL_TGACv1_577720_AA1881830 | | chr7B | 642269804 | 642272289 | 89704.525 | 2485 | R2b | IWGSC | | | | | | |
| Bsister | TaBS.3D | NA | TraesCS7D01G633200LC | 1 | 1 | OsMADS30-1 | PF00319; PF01486 | + | + | 214 | 23.95 | n | 0.11 | TRIAE_CS42_7DL_TGACv1_603225_AA1978720 | | chr7D | 578483710 | 578486060 | NA | 2350 | R3 | FGENESH_C | | | | | | |
| Bsister | TaBS.4A1 | HC | TraesCS3A01G090300 | 1 | 1 | OsMADS30-2 | PF00319; PF01486 | + | + | 225 | 25.61 | y | 0.31 | TRIAE_CS42_3AS_TGACv1_211758_AA0693780 | Traes_3AS_B4247C855.1 | chr3A | 57481565 | 57483399 | 232.404 | 1834 | R1 | IWGSC | TaMADS51 | | | | | |
| Bsister | TaBS.4A2 | HC | TraesCS3A01G090900 | 1 | 1 | OsMADS30-2 | PF00319; PF01486 | + | + | 227 | 25.95 | y | 1.28 | TRIAE_CS42_3AS_TGACv1_212037_AA0697010 | | chr3A | 57715803 | 57717750 | 455711.034 | 1947 | R1 | IWGSC | | | | | | |
| Bsister | TaBS.4B | HC | TraesCS3B01G105700 | 1 | 2 | OsMADS30-2 | PF00319; PF01486 | + | + | 226 | 25.71 | y | 0.43 | TRIAE_CS42_3B_TGACv1_222256_AA0760560 | TRAES3BF068500020CFD_c1 | chr3B | 71553642 | 71555571 | 79888.371 | 1929 | R2a | IWGSC | TaMADS64 | | | | | |
| Bsister | TaBS.4D1 | HC | TraesCS3D01G090300 | 1 | 1 | OsMADS30-2 | PF00319; PF01486 | + | + | 219 | 24.80 | y | 5.92 | TRIAE_CS42_3DS_TGACv1_271826_AA0908850 | | chr3D | 45922048 | 45923939 | 28.954 | 1891 | R1 | IWGSC | | | | | | |
| Bsister | TaBS.4D2 | HC | TraesCS3D01G090600 | 1 | 1 | OsMADS30-2 | PF00319; PF01486 | + | + | 184 | 21.10 | n | 0.62 | TRIAE_CS42_3DS_TGACv1_271826_AA0908900 | | chr3D | 45952893 | 45954851 | 9264.685 | 1958 | R1 | FGENESH+ | | | | | | |
| Bsister | TaBS.5A1 | HC | TraesCS7A01G475000 | 1 | 1 | OsMADS30-3 | PF00319; PF01486 | + | + | 225 | 25.48 | n | 0.33 | TRIAE_CS42_7AL_TGACv1_557405_AA1780900 | | chr7A | 670236002 | 670243109 | 323.041 | 7107 | R3 | IWGSC | | | | | | |
| Bsister | TaBS.5A2 | HC | TraesCS2A01G475000 | 1 | 1 | OsMADS30-3 | PF01486 | - | + | 100 | 11.52 | y | 0.54 | TRIAE_CS42_2AL_TGACv1_096226_AA0317960 | | chr2A | 715825308 | 715829370 | 50171.823 | 4062 | R3 | IWGSC | | | | | | |
| Bsister | TaBS.5A3 | HC | TraesCS2A01G153100 | 1 | 1 | OsMADS30-3 | PF01486 | - | + | 100 | 11.51 | y | 0.26 | | | chr2A | 101127995 | 101129131 | 31730.315 | 1136 | R2a | IWGSC | | | | | | |
| Bsister | TaBS.5A4 | HC | TraesCS4A01G137500 | 1 | 1 | OsMADS30-3 | PF01486 | - | + | 100 | 11.43 | y | 11.68 | TRIAE_CS42_4AS_TGACv1_306839_AA1013980 | | chr4A | 196783056 | 196785986 | 227105.213 | 2930 | C | IWGSC | | | | | | |
| Bsister | TaBS.5A5 | LC | TraesCS1A01G491400LC | 1 | 1 | OsMADS30-3 | PF01486 | - | + | 124 | 14.34 | n | 0.00 | TRIAE_CS42_1AL_TGACv1_000468_AA0012840 | | chr1A | 525835187 | 525838285 | NA | 3098 | R3 | IWGSC | | | | | | |
| Bsister | TaBS.5D | LC | TraesCS7D01G632400LC | 1 | 1 | OsMADS30-3 | PF00319; PF01486 | + | + | 176 | 19.21 | y | 0.63 | TRIAE_CS42_7DL_TGACv1_603629_AA1986570 | | chr7D | 578135926 | 578137455 | 346.255 | 1529 | R3 | IWGSC | | | | | | |
| Bsister | TaBS.6A1 | LC | TraesCS6A01G325500LC | 1 | 1 | OsMADS30-4 | PF00319 | + | - | 151 | 16.45 | y | 3.11 | TRIAE_CS42_6AL_TGACv1_471937_AA1516060 | | chr6A | 307893215 | 307899958 | 173836.995 | 6743 | C | IWGSC | | | | | | |
| Bsister | TaBS.6A2 | LC | TraesCS2A01G733800LC | 1 | 1 | OsMADS30-4 | PF00319 | + | - | 134 | 14.74 | n | 0.23 | | | chr2A | 766001193 | 766001597 | NA | 404 | R3 | IWGSC | | | | | | |
| Bsister | TaBS.6B1 | LC | TraesCS3B01G236300LC | 1 | 1 | OsMADS30-4 | PF00319 | + | - | 135 | 14.80 | y | 4.61 | TRIAE_CS42_3B_TGACv1_220599_AA0710540 | | chr2B | 174983258 | 174984136 | 5056.671 | 878 | R2a | IWGSC | | | | | | |
| Bsister | TaBS.6B2 | LC | TraesCS2B01G665600LC | 1 | 1 | OsMADS30-4 | PF00319 | + | - | 116 | 12.53 | y | 0.45 | TRIAE_CS42_2BL_TGACv1_130059_AA0402620 | | chr2B | 628948858 | 628949830 | 3673.493 | 972 | R2b | IWGSC | | | | | | |
| Bsister | TaBS.7B | HC | TraesCS1B01G467500 | 1 | 1 | OsMADS30-5 | PF00319 | + | - | 98 | 10.89 | n | 0.27 | TRIAE_CS42_1BL_TGACv1_030365_AA0088090 | | chr1B | 677273338 | 677274112 | NA | 774 | R3 | IWGSC | | | | | | |
| Bsister | TaBS.7D1 | NA | no gene predicted at site | 1 | 1 | OsMADS30-5 | PF00319; PF01486 | + | + | 503 | 54.83 | n | NA | | | chr1D | 486299358 | 486304128 | NA | 4770 | R3 | FGENESH+ | | | | | | |
| Bsister | TaBS.7D2 | HC | TraesCS1D01G441200 | 1 | 1 | OsMADS30-5 | PF00319; PF01486 | + | + | 187 | 20.95 | n | 0.00 | TRIAE_CS42_1DL_TGACv1_061515_AA0197450 | | chr1D | 486249793 | 486251742 | 47.616 | 1949 | R3 | IWGSC | | | | | | |
| Bsister | TaBS.8A | LC | TraesCS4A01G044400LC | 1 | 1 | OsMADS30-6 | PF00319; PF00448 | + | - | 214 | 23.96 | y | 3.12 | TRIAE_CS42_4AS_TGACv1_306506_AA1009250 | | chr4A | 38737436 | 38747176 | 13858.378 | 9740 | R1 | IWGSC | | | | | | |
| Bsister | TaBS.8D1 | LC | TraesCS2D01G714300LC | 1 | 1 | OsMADS30-6 | PF00319 | + | - | 87 | 9.48 | n | 0.43 | | | chr2D | 642350466 | 642350729 | NA | 263 | R3 | IWGSC | | | | | | |
| Bsister | TaBS.8D2 | LC | TraesCS3D01G045700LC | 1 | 1 | OsMADS30-6 | PF00319 | + | - | 94 | 10.51 | n | 0.75 | | | chr3D | 21007224 | 21007508 | 24914.54 | 284 | R1 | IWGSC | | | | | | |
| Bsister | TaBS.9A | HC | TraesCS7A01G474900 | 1 | 1 | OsMADS30-5 | PF00319 | + | - | 157 | 17.89 | n | 0.34 | TRIAE_CS42_7AL_TGACv1_557712_AA1785060 | Traes_7AL_67921A952.1 | chr7A | 670175149 | 670181113 | 54.889 | 5964 | R3 | IWGSC | TaMADS141 | | | | | |
| Bsister | TaBS.9B | LC | TraesCS7B01G633300LC | 1 | 1 | OsMADS30-5 | PF00319 | + | - | 146 | 16.56 | n | 0.38 | TRIAE_CS42_7BL_TGACv1_577784_AA1883230 | | chr7B | 641684748 | 641685559 | 584.245 | 811 | R2b | IWGSC | | | | | | |
| Bsister | TaBS.9D | HC | TraesCS3D01G102700 | 1 | 1 | OsMADS30-5 | PF00319; PF01486 | + | + | 229 | 26.27 | n | 1.68 | TRIAE_CS42_3DS_TGACv1_274303_AA0935370 | | chr3D | 55219536 | 55221713 | 45631.251 | 2177 | R2a | IWGSC | | | | | | |
| FLC | TaFLC.1A | HC | TraesCS7A01G260900 | 1 | 1 | OsMADS37 | PF00319 | + | (+) | 205 | 23.31 | n | 4.77 | TRIAE_CS42_7AS_TGACv1_569081_AA1806710 | | chr7A | 254389981 | 254432283 | 205193.79 | 42302 | C | IWGSC | TaMADS151 | | | | | |
| FLC | TaFLC.1B | HC | TraesCS7B01G158900 | 1 | 1 | OsMADS37 | PF00319 | + | (+) | 217 | 24.64 | n | 5.2 | TRIAE_CS42_7BS_TGACv1_592051_AA1929310 | | chr7B | 215932127 | 215969346 | 195524.041 | 37219 | C | IWGSC | TaMADS157 | | | | | |
| FLC | TaFLC.1D1 | HC | TraesCS7D01G261900 | 1 | 1 | OsMADS37 | PF00319 | + | (+) | 219 | 24.80 | y | 1.56 | TRIAE_CS42_7DS_TGACv1_621557_AA2019260 | | chr7D | 238973490 | 239005877 | 345.951 | 32387 | C | IWGSC | | | | | | |
| FLC | TaFLC.1D2 | LC | TraesCS7D01G328100LC | 1 | 1 | OsMADS37 | PF00319 | + | - | 103 | 11.09 | n | 1.11 | TRIAE_CS42_7DS_TGACv1_622625_AA2042820 | | chr7D | 239351828 | 239385957 | 162450.609 | 34129 | C | IWGSC | TaMADS171 | | | | | |
| FLC | TaFLC.1D3 | LC | TraesCS3D01G517300LC | 1 | 1 | OsMADS51-4 | PF00319 | + | - | 81 | 9.58 | n | 0.10 | | | chr3D | 541298581 | 541299906 | 71.313 | 1325 | R2b | IWGSC | | | | | | |
| FLC | TaFLC.2A | HC | TraesCS5A01G520200 | 2 | 1 | OsMADS51-2 | PF00319 | + | (+) | 202 | 22.73 | y | 25.39 | TRIAE_CS42_5AL_TGACv1_376789_AA1240970 | | chr5A | 681001866 | 681004489 | NA | 2623 | R3 | IWGSC | | | | | | |
| FLC | TaFLC.2B | HC | TraesCS4B01G351500 | 1 | 1 | OsMADS51-2 | PF00319 | + | (+) | 164 | 18.15 | n | 16.42 | TRIAE_CS42_4BL_TGACv1_320827_AA1049610 | | chr4B | 643648399 | 643651612 | 4445.216 | 3213 | R3 | IWGSC | | | | | | |
| FLC | TaFLC.2D | HC | TraesCS4D01G346300 | 1 | 1 | OsMADS51-2 | PF00319 | + | (+) | 207 | 22.80 | y | 14.69 | TRIAE_CS42_4DL_TGACv1_344001_AA1142360 | | chr4D | 500311335 | 500314011 | NA | 2676 | R3 | IWGSC | | | | | | |
| FLC | TaFLC.3A | HC | TraesCS3A01G434400 | 1 | 1 | OsMADS51-3 | PF00319 | + | (+) | 167 | 18.42 | y | 0.6 | TRIAE_CS42_3AL_TGACv1_195330_AA0648110 | | chr3A | 676207398 | 676244073 | 605.412 | 36675 | R3 | IWGSC | | | | | | |
| FLC | TaFLC.3B | HC | TraesCS3B01G469700 | 1 | 1 | OsMADS51-3 | PF00319 | + | (+) | 167 | 18.52 | y | 9.65 | TRIAE_CS42_3B_TGACv1_221184_AA0733140 | | chr3B | 716709509 | 716741965 | 224.657 | 32456 | R2b | IWGSC | TaMADS58 | | | | | |
| FLC | TaFLC.3D | HC | TraesCS3D01G427700 | 1 | 1 | OsMADS51-3 | PF00319 | + | (+) | 166 | 18.14 | y | 19.06 | TRIAE_CS42_3DL_TGACv1_249684_AA0854270 | | chr3D | 541165330 | 541197913 | 100.668 | 32583 | R2b | IWGSC | | | | | | |
| FLC | TaFLC.4A1 | HC | TraesCS3A01G435000 | 1 | 1 | OsMADS51-4 | PF00319 | + | (+) | 167 | 18.55 | y | 0.57 | TRIAE_CS42_3AL_TGACv1_195926_AA0655730 | | chr3A | 677141526 | 677172275 | NA | 30749 | R3 | IWGSC | | | | | | |
| FLC | TaFLC.4A2 | HC | TraesCS3A01G434700 | 1 | 1 | OsMADS51-4 | PF00319 | + | - | 102 | 11.66 | n | 0.62 | | | chr3A | 676849485 | 676853285 | 32.338 | 3800 | R3 | FGENESH_C | | | | | | |
| FLC | TaFLC.4B2 | HC | TraesCS3B01G470000 | 1 | 1 | OsMADS51-4 | PF00319 | + | (+) | 167 | 18.48 | n | 13.78 | TRIAE_CS42_3B_TGACv1_220587_AA0709710 | | chr3B | 717278544 | 717310236 | NA | 31692 | R2b | IWGSC | | | | | | |
| FLC | TaFLC.4D1 | HC | TraesCS3D01G428000 | 1 | 1 | OsMADS51-4 | PF00319 | + | (+) | 159 | 17.57 | y | 5.18 | TRIAE_CS42_3DL_TGACv1_251054_AA0876770 | | chr3D | 541598279 | 541632777 | NA | 34498 | R2b | IWGSC | | | | | | |
| FLC | TaFLC.5A | HC | TraesCS3A01G432900 | 1 | 1 | OsMADS51-4 | PF00319 | + | (+) | 155 | 17.28 | y | 8.26 | TRIAE_CS42_3AL_TGACv1_194409_AA0632550 | | chr3A | 674255517 | 674279708 | 1927.69 | 24191 | R3 | IWGSC | | | | | | |
| FLC | TaFLC.5D | LC | TraesCS3D01G514600LC | 1 | 1 | OsMADS51-4 | PF00319 | + | (+) | 164 | 18.12 | n | 10.31 | TRIAE_CS42_3DL_TGACv1_249535_AA0851090 | | chr3D | 538867195 | 538901906 | 2263.424 | 34711 | R2b | IWGSC | | | | | | |
| FLC | TaFLC.6A | HC | TraesCS3A01G434900 | 1 | 1 | OsMADS51-4 | PF00319 | + | (+) | 167 | 18.40 | y | 1.86 | TRIAE_CS42_3AL_TGACv1_194694_AA0637970 | | chr3A | 676885623 | 676906103 | 235.423 | 20480 | R3 | IWGSC | | | | | | |
| FLC | TaFLC.6B | LC | TraesCS3B01G683500LC | 1 | 1 | OsMADS51-4 | PF00319 | + | - | 78 | 8.92 | n | 0.16 | | | chr3B | 716966622 | 716966858 | 311.686 | 236 | R2b | IWGSC | | | | | | |
| FLC | TaFLC.6D | HC | TraesCS3D01G427900 | 1 | 1 | OsMADS51-4 | PF00319 | + | (+) | 167 | 18.37 | n | 1.68 | TRIAE_CS42_3DL_TGACv1_249571_AA0851680 | | chr3D | 541371219 | 541403858 | 194.421 | 32639 | R2b | IWGSC | | | | | | |
| MIKC* | TaMIKC.1A | HC | TraesCS4A01G169000 | 1 | 1 | OsMADS68 | PF00319; PF01486 | + | + | 387 | 42.98 | y | 33.74 | TRIAE_CS42_4AL_TGACv1_288685_AA0955560 | | chr4A | 423891199 | 423894626 | 170454.479 | 3427 | R2b | IWGSC | | | | | | |
| MIKC* | TaMIKC.1B | HC | TraesCS4B01G150200 | 1 | 2 | OsMADS68 | PF00319; PF01486 | + | + | 389 | 43.30 | y | 24.38 | TRIAE_CS42_4BS_TGACv1_328206_AA1084610 | | chr4B | 230008306 | 230011748 | 277952.736 | 3442 | C | IWGSC | | | | | | |
| MIKC* | TaMIKC.1D | HC | TraesCS4D01G147000 | 1 | 2 | OsMADS68 | PF00319; PF01486 | + | + | 389 | 43.21 | y | 20.08 | TRIAE_CS42_4DS_TGACv1_361328_AA1165800 | | chr4D | 137764701 | 137768122 | 269347.965 | 3421 | C | IWGSC | TaMADS95 | | | | | |
| MIKC* | TaMIKC.2A | HC | TraesCS4A01G290700 | 1 | 1 | OsMADS62 | PF00319; PF01486 | + | + | 391 | 42.91 | y | 35.26 | TRIAE_CS42_4AL_TGACv1_289782_AA0976740 | | chr4A | 594349105 | 594351879 | NA | 2774 | R3 | FGENESH+ | | | | | | |
| MIKC* | TaMIKC.2B | HC | TraesCS4B01G023300 | 1 | 1 | OsMADS62 | PF00319; PF01486 | + | + | 362 | 39.65 | y | 21.52 | TRIAE_CS42_4BS_TGACv1_328777_AA1093510 | | chr4B | 17017110 | 17019148 | 212989.158 | 2038 | R1 | FGENESH+ | | | | | | |
| MIKC* | TaMIKC.2D | HC | TraesCS4D01G021100 | 1 | 1 | OsMADS62 | PF00319; PF01486 | + | + | 353 | 38.67 | y | 53.23 | TRIAE_CS42_4DS_TGACv1_361789_AA1172760 | | chr4D | 9069291 | 9071423 | 128693.278 | 2132 | R1 | IWGSC | TaMADS96 | | | | | |
| OsMADS32 | TaMADS32A | HC | TraesCS3A01G284400 | 1 | 1 | OsMADS32 | PF00319; PF01486 | + | + | 213 | 24.38 | y | 17.19 | TRIAE_CS42_3AL_TGACv1_197941_AA0667660 | Traes_3AL_219064574.1 | chr3A | 513428784 | 513431087 | 41604.059 | 2303 | R2b | IWGSC | TaMADS46 | TaWM16 | | WM16 | AM502882 | |
| OsMADS32 | TaMADS32B | HC | TraesCS3B01G318300 | 1 | 1 | OsMADS32 | PF00319; PF01486 | + | + | 196 | 22.32 | y | 3.30 | TRIAE_CS42_3B_TGACv1_220686_AA0715430 | TRAES3BF048900050CFD_c1 | chr3B | 513254002 | 513256449 | 166386.453 | 2447 | R2b | IWGSC | TaMADS62 | TaWM16 | | WM16 | AM502882 | |
| OsMADS32 | TaMADS32D | HC | TraesCS3D01G284200 | 1 | 1 | OsMADS32 | PF00319; PF01486 | + | + | 196 | 22.33 | y | 35.62 | | | chr3D | 393524475 | 393526973 | 122855.097 | 2498 | R2b | IWGSC | | TaWM16 | | WM16 | AM502882 | |
| PI | TaPI.1A | HC | TraesCS1A01G264300 | 1 | 1 | OsMADS4 | PF00319; PF01486 | + | + | 208 | 24.34 | y | 135.40 | TRIAE_CS42_1AL_TGACv1_003698_AA0051100 | Traes_1AL_1B5F51626.1 | chr1A | 459946840 | 459948904 | 65886.283 | 2064 | R2b | IWGSC | TaMADS1 | | | | | |
| PI | TaPI.1B | HC | TraesCS1B01G275000 | 1 | 1 | OsMADS4 | PF00319; PF01486 | + | + | 208 | 24.37 | y | 26.84 | TRIAE_CS42_1BL_TGACv1_032898_AA0135200 | | chr1B | 482450684 | 482455256 | 194818.082 | 4572 | R2b | IWGSC | | | | WM14 | | |
| PI | TaPI.1D | HC | TraesCS1D01G264500 | 1 | 2 | OsMADS4 | PF00319; PF01486 | + | + | 218 | 25.24 | y | 18.17 | TRIAE_CS42_1DL_TGACv1_062601_AA0217110 | Traes_1DL_D25CDC57D.1 | chr1D | 360386578 | 360391321 | 125858.472 | 4743 | R2b | IWGSC | TaMADS19 | TaPI-1 | | WM14 | AM502880 | |
| PI | TaPI.2A | HC | TraesCS3A01G406500 | 1 | 2 | OsMADS2 | PF00319; PF01486 | + | + | 207 | 23.81 | y | 162.15 | TRIAE_CS42_U_TGACv1_641952_AA2108230 | Traes_3AL_01EE581F9.1 | chr3A | 651582315 | 651587619 | 22667.898 | 5304 | R2b | IWGSC | TaMADS45 | TaPI-2 | TaMADWPI2 | WM15 | AM502881 | |
| PI | TaPI.2B | HC | TraesCS3B01G440200 | 1 | 3 | OsMADS2 | PF00319; PF01486 | + | + | 141 | 16.06 | y | 153.78 | TRIAE_CS42_3B_TGACv1_224968_AA0803750 | TRAES3BF021600020CFD_c1 | chr3B | 679642902 | 679645420 | 37064.089 | 2518 | R2b | IWGSC | TaMADS60 | TaPI-2 | | | AM502881 | |
| PI | TaPI.2D | HC | TraesCS3D01G401700 | 1 | 2 | OsMADS2 | PF00319; PF01486 | + | + | 209 | 24.09 | y | 141.27 | TRIAE_CS42_3DL_TGACv1_249089_AA0837300 | Traes_3DL_0DEA285BD.1 | chr3D | 516382070 | 516384654 | 22482.541 | 2584 | R2b | IWGSC | TaMADS69 | | | | | |
| SOC1 | TaSOC1.1A | HC | TraesCS1A01G199600 | 1 | 2 | OsMADS56-1 | PF00319; PF01486 | + | + | 229 | 25.74 | y | 25.52 | TRIAE_CS42_1AL_TGACv1_001072_AA0024490 | Traes_1AL_6B108514B.1 | chr1A | 358761357 | 358772027 | 456.518 | 10670 | R2b | IWGSC | TaMADS3 | TaSOC1-1A | | WM1A | AM502861 | |
| SOC1 | TaSOC1.1B | HC | TraesCS1B01G214500 | 1 | 1 | OsMADS56-1 | PF00319; PF01486 | + | + | 218 | 24.59 | y | 1.87 | TRIAE_CS42_1BL_TGACv1_031173_AA0109140 | Traes_1BL_F1D5BF5F8.1 | chr1B | 389212932 | 389224699 | 327.586 | 11767 | R2b | IWGSC | TaMADS10 | TaSOC1-1B | | WM1B | AM505862 | |
| SOC1 | TaSOC1.1D | HC | TraesCS1D01G203300 | 1 | 2 | OsMADS56-1 | PF00319; PF01486 | + | + | 229 | 25.80 | y | 18.29 | TRIAE_CS42_1DL_TGACv1_061229_AA0189260 | | chr1D | 287456845 | 287468338 | 614.078 | 11493 | R2b | IWGSC | | | TaMADAGL7 | | | DQ512363 |
| SOC1 | TaSOC1.2A | HC | TraesCS6A01G011700 | 1 | 2 | NA | PF00319; PF01486 | + | + | 262 | 28.77 | y | 12.57 | TRIAE_CS42_6AS_TGACv1_486857_AA1565890 | Traes_6AS_D6ABA1D79.1 | chr6A | 5358050 | 5364845 | 97903.736 | 6795 | R1 | IWGSC | TaMADS125 | | TaMADAGL23 | | | |
| SOC1 | TaSOC1.2B | HC | TraesCS6B01G017900 | 1 | 1 | NA | PF00319; PF01486 | + | + | 258 | 28.26 | y | 22.80 | TRIAE_CS42_6BS_TGACv1_513226_AA1635080 | Traes_6BS_43B59D772.1 | chr6B | 11317455 | 11326092 | 151240.745 | 8637 | R1 | IWGSC | TaMADS128 | | TaMADAGL1 | | DQ512330 | |
| SOC1 | TaSOC1.2D | HC | TraesCS6D01G014600 | 1 | 1 | NA | PF00319; PF01486 | + | + | 259 | 28.50 | y | 33.56 | TRIAE_CS42_6DS_TGACv1_542755_AA1729670 | Traes_6DS_D50A0F246.1 | chr6D | 6012829 | 6020283 | 79890.76 | 7454 | R1 | IWGSC | TaMADS139 | TaSOC1-2 | | WM18 | AM502883 | |
| SOC1 | TaSOC1.3A | HC | TraesCS5A01G515500 | 1 | 1 | OsMADS50 | PF00319; PF01486 | + | + | 220 | 24.57 | y | 20.56 | TRIAE_CS42_5AL_TGACv1_374176_AA1192420 | Traes_5AL_21C395CA8.1 | chr5A | 678931525 | 678945164 | 2056.702 | 13639 | R3 | IWGSC | TaMADS99 | | | | | |
| SOC1 | TaSOC1.3B | HC | TraesCS4B01G346700 | 1 | 1 | OsMADS50 | PF00319; PF01486 | + | + | 222 | 24.97 | y | 17.96 | TRIAE_CS42_4BL_TGACv1_320854_AA1050320 | Traes_4BL_B075EFE84.1 | chr4B | 640289448 | 640312189 | 3336.21 | 22741 | R3 | IWGSC | TaMADS84 | TaSOC1-3B | | WM21B | AM502888 | DQ512354 |
| SOC1 | TaSOC1.3D | HC | TraesCS4D01G341700 | 1 | 1 | OsMADS50 | PF00319; PF01486 | + | + | 222 | 24.80 | y | 30.14 | TRIAE_CS42_4DL_TGACv1_343157_AA1130760 | Traes_4DL_5633C0561.1 | chr4D | 498384057 | 498398218 | 1913.117 | 14161 | R3 | IWGSC | TaMADS90 | TaSOC1-3A | TaMADS-box | WM21D | AM502887 | AB281427 |
| SOC1 | TaSOC1.4A | LC | TraesCS7A01G051700LC | 1 | 1 | OsMADS50-2 | PF00319; PF01486 | + | + | 194 | 22.28 | n | 0.00 | TRIAE_CS42_7AS_TGACv1_571927_AA1850810 | | chr7A | 19183367 | 19186523 | 49040.745 | 3156 | R1 | IWGSC | | | | | | |
| SOC1 | TaSOC1.4B | NA | no gene predicted at site | 1 | 1 | OsMADS50-2 | PF00319 | + | - | 119 | 13.33 | n | NA | | | chr4B | 648096828 | 648098700 | NA | 1872 | R3 | FGENESH+ | | | | | | |
| SOC1 | TaSOC1.4D | HC | TraesCS7D01G037000 | 1 | 1 | OsMADS50-2 | PF00319 | + | - | 61 | 6.72 | n | 0.00 | TRIAE_CS42_7DS_TGACv1_621953_AA2030230 | | chr7D | 18857672 | 18873778 | 45562.756 | 16106 | R1 | IWGSC | | | | | | |
| SOC1 | TaSOC1.5A | HC | TraesCS1A01G199900 | 1 | 1 | OsMADS56 | PF00319; PF01486 | + | + | 228 | 25.68 | n | 1.60 | TRIAE_CS42_1AL_TGACv1_000686_AA0017040 | Traes_1AL_6F5982F88.1 | chr1A | 359228545 | 359245835 | 98970.784 | 17290 | R2b | IWGSC | TaMADS4 | | | | | |
| SOC1 | TaSOC1.5B | HC | TraesCS1B01G214600 | 1 | 1 | OsMADS56 | PF00319; PF01486 | + | + | 223 | 24.97 | y | 2.09 | TRIAE_CS42_1BL_TGACv1_030680_AA0097800 | | chr1B | 389552285 | 389557966 | 90879.878 | 5681 | R2b | FGENESH+ | | | | | | |
| SOC1 | TaSOC1.5D | HC | TraesCS1D01G203400 | 1 | 1 | OsMADS56 | PF00319; PF01486 | + | + | 231 | 25.88 | y | 1.71 | TRIAE_CS42_1DL_TGACv1_061144_AA0186880 | Traes_1DL_D5BBCA2D0.1 | chr1D | 288082416 | 288103808 | 68824.675 | 21392 | R2b | IWGSC | TaMADS20 | | | | | |
| SVP | TaSVP.1A | HC | TraesCS6A01G313800 | 1 | 1 | OsMADS22 | PF00319; PF01486 | + | + | 228 | 25.44 | y | 38.07 | TRIAE_CS42_6AL_TGACv1_473223_AA1529480 | Traes_6AL_A93C6F2FC.1 | chr6A | 550637111 | 550643948 | NA | 6837 | R2b | IWGSC | TaMADS120 | TaSVP-1A | | WM22A | AM502889 | |
| SVP | TaSVP.1B | HC | TraesCS6B01G343900 | 1 | 2 | OsMADS22 | PF00319; PF01486 | + | + | 228 | 25.46 | y | 84.81 | TRIAE_CS42_6BL_TGACv1_500914_AA1611650 | Traes_6BL_E1793636C.1 | chr6B | 605795122 | 605801747 | NA | 6625 | R2b | IWGSC | TaMADS127 | TaSVP-1B | | WM22B | AM502890 | |
| SVP | TaSVP.1D | HC | TraesCS6D01G293200 | 1 | 2 | OsMADS22 | PF00319; PF01486 | + | + | 228 | 25.43 | y | 80.13 | TRIAE_CS42_6DL_TGACv1_527648_AA1706840 | Traes_6DL_D8C9E421C.1 | chr6D | 403969730 | 403976356 | NA | 6626 | R2b | IWGSC | TaMADS135 | | TaMADAGL11 | | | |
| SVP | TaSVP.2A | HC | TraesCS7A01G175200 | 1 | 2 | OsMADS55 | PF00319; PF01486 | + | + | 226 | 25.20 | y | 32.92 | TRIAE_CS42_7AS_TGACv1_571446_AA1847780 | Traes_7AS_CA6E66D75.1 | chr7A | 128826237 | 128833032 | 124594.7 | 6795 | R2a | IWGSC | TaMADS149 | TaSVP-2B | | WM24 | AM502892 | |
| SVP | TaSVP.2B | HC | TraesCS7B01G080300 | 1 | 2 | OsMADS55 | PF00319; PF01486 | + | + | 228 | 25.47 | y | 98.45 | TRIAE_CS42_7BS_TGACv1_592420_AA1937780 | | chr7B | 90189439 | 90195309 | 124805.67 | 5870 | R2a | IWGSC | | TaSVP-2B | | WM24 | AM502892 | |
| SVP | TaSVP.2D | HC | TraesCS7D01G176700 | 1 | 1 | OsMADS55 | PF00319; PF01486 | + | + | 226 | 25.22 | y | 57.29 | TRIAE_CS42_7DS_TGACv1_622344_AA2038090 | Traes_7DS_366A869CF.1, Traes_7DS_90 | chr7D | 128912242 | 128919539 | 108690.291 | 7297 | R2a | IWGSC | TaMADS169, TaMADS177 | | | | | |
| SVP | TaSVP.3A | HC | TraesCS4A01G002600 | 1 | 1 | OsMADS47 | PF01486 | - | + | 163 | 18.48 | y | 51.22 | TRIAE_CS42_4AS_TGACv1_307592_AA1021790 | Traes_4AS_BA5BB8032.1 | chr4A | 2402442 | 2412252 | 17533.393 | 9810 | R1 | IWGSC | TaMADS78 | TaSVP-3A | | WM28A | AM502896 | |
| SVP | TaSVP.3B | HC | TraesCS4B01G302600 | 1 | 3 | OsMADS47 | PF00319; PF01486 | + | + | 226 | 25.40 | y | 124.20 | TRIAE_CS42_4BL_TGACv1_321995_AA1067550 | Traes_4BL_410DEBFD3.1 | chr4B | 589180009 | 589195455 | 51093.993 | 15446 | R3 | IWGSC | TaMADS82 | TaSVP-3B | TaMADAGL13 | | AM502897 | |
| SVP | TaSVP.3D | HC | TraesCS4D01G301100 | 1 | 1 | OsMADS47 | PF00319; PF01486 | + | + | 226 | 25.37 | y | 85.03 | TRIAE_CS42_4DL_TGACv1_344084_AA1143460 | Traes_4DL_67EFB6303.1, Traes_4DL_AC | chr4D | 469304201 | 469319980 | 29064.077 | 15779 | R3 | IWGSC | TaMADS91, TaMADS93 | | | | | |

| NCBI GenBank | TC sequence(TaG | Corresponding E | NCBI ID | NCBI ID | NCBI GenBank G | Shitsukawa 2007 | Hirabayashi and Murai 2( | Murai et al. 1998 | Zhao 2006a | ZhaoT 2006b | Wang 2015 | Mizumoto 2009 | Shitsukawa 2007 | Kane 2005 | Fu 2005 | Ciaffi 2005 | Hama 2004 | Meguro 2003 | Ruelens 2013 | Yamada 2009 |
|---|---|---|---|---|---|---|---|---|---|---|---|---|---|---|---|---|---|---|---|---|
| | TC267375 | BJ276769 | AM502866 | CAM59044 | WM4 | | | | | | | | | | | | | | | |
| 664 | TC267376 | BJ255649 | DQ512341 | ABF57925 | TaAGL24 | | | | | TaAGL24 | | | WLHS-A | | | | | | | |
| | TC267376 | BJ255649 | DQ512356 | | | | | | | | | | WLHS-B | | | | | | | |
| | | | DQ534492 | ABG21011 | TaMADS7 | | | | | | | | WLHS-D | | | | | | | |
| | | | | | | | | | | TaAGL4 | | | | | | | | | | |
| ABF57946 | | | DQ534491 | | TaMADS3 | | | | | TaAGL5 | | | | | | | | | | |
| ABF57943 | TC253709 | CJ513681 | AM502886 | CAM59064 | WM20 | | | | | TaAGL8 | | | | | | | | | | |
| | | | DQ512370 | ABF57954 | TaAGL40 | | | | | TaAGL40 | | | | | | | | | | |
| | | | DQ512351,DQ51 | ABF57935,ABF57 | TaAGL34,TaAGL3 | | | | | TaAGL3, TaAGL34 | | | | | | | | | | |
| | TC250645 | CJ574682 | | | | | | | | TaAGL27 | | | | | | | | | | |
| | | | | | | | | | | next to TaVRT1 | | | | | | | | | | |
| | | | DQ512334 | ABF57918 | TaAGL16 | | | | | TaAGL16 | | | WSEP-A | | | | | | | |
| | TC250641 | DY761322 | DQ512345 | ABF57929 | TaAGL28 | | | | | TaAGL6 | | | WSEP-B | | | | | | | |
| | | | DQ512348 | ABF57932 | TaAGL30 | | | | | TaAGL30 | | | WSEP-D | | | | | | | |
| | TC250643 | CJ687184 | | | | | | | TaMADS1 | | | | | | | | | | | |
| | TC250643 | CJ687184 | | | | | | | TaMADS1 | | | | | | | | | | | |
| | | | | | | | WAG-1 | | | | | | | | | | | | | |
| | TC251484 | DY761328 | | CAM59041 | | | | | | | | | | | | WAG | | WAG | | |
| | TC240430 | CJ579736 | AB465688 | | | | | | | TaAGL39 | | WAG-2 (BAH56656) | | | | | | | | |
| | TC240430 | CJ579736 | | | | | | | | TaAGL39 | WAG-2o (ALM58844) | | | | | | | | | |
| | | | | | | | WAG-2 | | | | WAG-2m (ALM58842) | | | | | | | | | |
| | TC233894 | CJ680037 | DQ512337 | ABF57921 | TaAGL2 | | | | | TaAGL2 | | | | | | | | | | WSTK |
| | | | | | | | | | | | | | | | | | | | | WSTK |
| ABF57948 | TC233894 | CJ680037 | DQ512349 | ABF57933 | TaAGL31 | | | | | TaAGL9 | | | | | | | | | | WSTK |
| | TC259764 | CJ686261 | | | | | | | | | | | | | | | | | | |
| | TC259764 | CJ686261 | | | | | | | | | | | | | | | | | | |
| | | | DQ512350 | ABF57934 | TaAGL32 | | | | | | | | | | | | | | | |
| | TC270320 | DR741518 | | CAM59056 | | | | | | TaAGL32 | | | | | | | | | | |
| | | CJ686932 | | CAM59078 | | | | | | | | | | | | | | | | |
| | TC245817 | BE445262 | | | | | | | | TaAGL6 | | | | | | | | | | |
| | TC245817 | BE445262 | DQ512329 | | TaAGL6 | | | | | TaAGL6 | | | | | | | | | | |
| | | CJ722516 | | | | | | | | | | | | | | | | | | |
| | | CJ722516 | | | | | | | | | | | | | | | | | | |
| | | CJ722516 | DQ512368 | ABF57952 | TaAGL17 | | | | | TaAGL17 | | | | | | | | | | |
| | TC245817 | BE445262 | | | | | | | | TaAGL6 | | | | | | | | | | |
| AB007505 | TC268200 | CK210628 | DQ512353 | ABF57937 | TaAGL37 | | | | | TaAGL37 | | | | | | | | | | |
| | TC268200 | CK210628 | AB007505 | BAA33458 | TaMADS#12 | | | TaMADS#12 | | | | | | | | | | | | |
| | TC268200 | CK210628 | | | | | | TaMADS12 | | TaAGL37 | | | | | | | | | | |
| | TC235598 | CD936729 | DQ512342 | ABF57926 | TaAGL25 | | | | | TaAGL25 | | | | | VRN-A1 | | | | | |
| | | | AB007504 | BAA33457 | TaMADS#11 | | | | | | | | | | | | | | | |
| | DQ512344 | ABF57928 | AB007504 | BAA33457 | TaMADS#11 | | | | | TaAGL27 | | | | | | | | | | |
| | | | | | | | | | | TaAGL10 | | | | | | | | | | |
| DQ534490 | TC270111 | CJ654733 | DQ512331 | ABF57915 | | | | | | TaAGL10 | | | | | | | | | | |
| | | | DQ512346 | ABF57930 | TaAGL29 | | | | | | | | | | | | | | | |
| | TC236763 | BE498620 | | CAM59049 | | | | | | TaAGL29 | | | | | | | | | | |
| | | | AB007506 | BAA33459 | TaMADS#51 | | | | | | | | | | | | TaMADS#51; TaAP3-like | | | |
| | TC238728 | DY761315 | AB107993 | | TaMADS#82 | | | TaMADS51 | | | | | | | | | TaMADS#82; TaAP3-like, | | | |
| | | | | | | | | | | | | | | | | | | | | WBSIS |
| | TC241889 | CD890142 | DQ512369 | ABF57953 | TaAGL35 | | | | | TaAGL35 | | | | | | | | | | WBSIS |
| | | | | | | | | | | | | | | | | | | | | WBSIS |
| | | | DQ512367 | ABF57951 | TaAGL12 | | | | | | | | | | | | | | TaAGL12 | |
| | | | | | | | | | | | | | | | | | | | TaAGL12 | |
| | | | DQ512367 | ABF57951 | TaAGL12 | | | | | | | | | | | | | | TaAGL12 | |
| | | | DQ512357 | ABF57941 | | | | | | TaAGL41 | | | | | | | | | | |
| | | | DQ512365 | ABF57949 | TaAGL33, TaAGL22 | | | | | | | | | | | | | | | |
| | | | DQ512366 | ABF57950 | TaAGL33, TaAGL22 | | | | | TaAGL33 | | | | | | | | | | |
| | | | DQ512365 | ABF57949 | TaAGL33, TaAGL22 | | | | | | | | | | | | | | | |
| | | | DQ512358 | ABF57942 | TaAGL42 | | | | | TaAGL42 | | | | | | | | | | |
| | TC266928 | CJ55336 | DQ923396 | | TaMADS1 | | | | | TaAGL14 | | | | | | MADS32 | | | | |
| | TC266928 | CJ55336 | DQ512361 | ABF57945 | TaAGL15 | | | | | TaAGL14 | | | | | | | | | | |
| | TC266928 | CJ55336 | DQ512360 | ABF57944 | TaAGL14 | | | | | TaAGL14 | | | | | | | | | | |
| | | | | | | | | | | | | | | | | | WPI-1D | | | |
| | | | | | | | | | | | | | | | | | WPI-1B | | | |
| | TC267188 | DY61309 | | CAM59058 | | | | | | | | | | | | | WPI-1A | | | |
| | TC235607 | CJ553283 | AB107992 | | | | | | | | | | | | | | WPI-2A | | | |
| | TC235607 | CJ553283 | AM502881 | CAM59059 | WM15 | | | | | | | | | | | | WPI-2B | | | |
| | | | DQ512343 | ABF57927 | TaAGL26 | | | | | TaAGL26 | | | | | | | WPI-2D | | | |
| | TC242238 | CD374116 | | | | | | | | TaAGL7 | | | | | | | | | | |
| | TC242238 | CD374116 | DQ512363 | ABF57947 | | | | | | | | | | | | | | | | |
| | | | | | | | | | | TaAGL7 | | | | | | | | | | |
| | | | DQ512340 | ABF57924 | TaAGL23 | | | | | TaAGL23 | | | | | | | | | | |
| | | | DQ512330 | | TaAGL1 | | | | | TaAGL1 | | | | | | | | | | |
| | TC255866 | BQ804616 | DQ512335 | ABF57919 | TaAGL18 | | | | | TaAGL18 | | | | | | | | | | |
| AB281427 | | | DQ512339 | ABF57923 | TaAGL21 | WSOC1 BAF56968.1 | | | | | | | | | | | | | | |
| ABF57938 | TC255111 | CD916236 | DQ512338 | ABF57922 | TaAGL20 | | | | | TaAGL38 | | | | | | | | | | |
| | TC255111 | CD916236 | DQ512354 | ABF57938 | TaAGL38 | | | | | TaAGL38 | | | | | | | | | | |
| | TC237274 | DY761306 | DQ512332 | ABF57916 | TaAGL11 | | | | | TaAGL11 | | | | | | | | | | |
| | TC237274 | DY761306 | DQ512332 | ABF57916 | TaAGL11 | | | | | TaAGL11 | | | | | | | | | | |
| | TC271223 | CK207974 | DQ512352 | ABF57936 | TaAGL36 | | | | | | | | | VRT-2 | | | | | | |
| | TC271223 | CK207974 | DQ512352 | ABF57936 | TaAGL36 | | | | | TaAGL36 | | | | VRT-2 | | | | | | |
| | | | DQ512352 | ABF57936 | TaAGL36 | | | | | | | | | | | | | | | |
| | | | DQ512336 | ABF57920 | TaAGL19 | | | | | | | | | | | | | | | |
| | TC269311 | CJ692094 | DQ512333 | ABF57917 | TaAGL13 | | | | | TaAGL13 | | | | | | | | | | |
| | TC269311 | CJ692094 | DQ923397 | ABL11476 | TaMADS10 | | | | | TaAGL13 | | | | | | | | | | |

**Table S3. Average distance to the nearest downstream MIKC-type gene in kbp in different chromosomal segments.** Distances between start points of neighboring MIKC-type genes have been calculated and averaged over chromosomal segments.

| Chromosome segment | mean distance in kb | std dev in kb | total no of genes | No of genes under <1000kb | % of genes under <1000kb |
|---|---|---|---|---|---|
| C | 152154.05 | 83794.14 | 19 | 3 | 15.8% |
| R1 | 45667.80 | 59310.20 | 53 | 29 | 54.7% |
| R2a | 128857.49 | 88909.30 | 24 | 2 | 8.3% |
| R2b | 55115.64 | 46177.91 | 57 | 10 | 17.5% |
| R3 | 29201.63 | 32435.34 | 28 | 10 | 35.7% |

**Table S4. List of all MIKC-type MADS-box genes predicted during this study.** Table includes gene name, IWGSC identifier (if available) and chromosome location, PFAM domain(s), predicted coding sequence and prediction algorithm and guiding sequence FGENESH+ and FGENESH_C predictions are guided by homolog protein or cDNA sequences, respectively.

| Clade | Gene name | IWGSC ID | IWGSC genomic locus | PFAM domains | gene prediction algorithm | guidance |
|---|---|---|---|---|---|---|
| AGL17 | TaAGL17.7B2 | no gene predicted at site | chr1B:40330406..40347378 | PF00319 | FGENESH+ | OsMADS57 |
| AGL17 | TaAGL17.6B1 | no gene predicted at site | chr7B:731976814..731982042 | PF00319; PF01486 | FGENESH+ | OsMADS57 |
| AGL17 | TaAGL17.11D3 | no gene predicted at site | chr7D:64490707..64494277 | PF00319 | FGENESH+ | OsMADS57 |
| AGL17 | TaAGL17.11A2 | no gene predicted at site | chr7A:68342914..68344465 | PF00319; PF01486 | FGENESH+ | OsMADS57 |
| AGL17 | TaAGL17.11A4 | no gene predicted at site | chr7A:68772222..68779045 | PF00319; PF01486 | FGENESH+ | OsMADS25 |
| AGL17 | TaAGL17.11D4 | no gene predicted at site | chr7D:64531588..64533657 | PF00319; PF01486 | FGENESH+ | OsMADS57 |
| Bsister | TaBS.7D1 | no gene predicted at site | chr1D:486299823..486304562 | PF00319; PF01486 | FGENESH+ | OsMADS57 |
| SOC1 | TaSOC1.4B | no gene predicted at site | chr4B:648096828..648098700 | PF00319 | FGENESH+ | OsMADS57 |
| SOC1 | TaSOC1.5B | TraesCS1B01G214600, TraesCS1B01G406400LC | chr1B:389552248..389598897 | PF00319; PF01486 | FGENESH+ | OsMADS50 |
| FLC | TaFLC.4A2 | TraesCS3A01G434700 | chr3A:676849485..676853285 | PF00319 | FGENESH_C | TaFLC.3A |
| Bsister | TaBS.4D2 | TraesCS3D01G090600 | chr3D:45952893..45954851 | PF00319; PF01486 | FGENESH+ | OsMADS29 |
| MIKC* | TaMIKC.2A | TraesCS4A01G290700 | chr4A:594349105..594351879 | PF00319; PF01486 | FGENESH+ | OsMADS62 |
| MIKC* | TaMIKC.2B | TraesCS4B01G023300 | chr4B:17017110..17019148 | PF00319; PF01486 | FGENESH+ | OsMADS62 |
| AGL17 | TaAGL17.10B | TraesCS5B01G002500, TraesCS5B01G002300LC | chr5B:3866145..3867846 | PF00319; PF01486 | FGENESH+ | OsMADS57 |
| SEP | TaSEP1.5B1 | TraesCS7B01G020900, TraesCS7B01G032300LC | chr7B:18498249..18506298 | PF00319; PF01486 | FGENESH+ | OsMADS1 |
| SEP | TaSEP1.5B2 | TraesCS7B01G021100, TraesCS7B01G021000 | chr7B:19111832..19141655 | PF00319; PF01486 | FGENESH+ | OsMADS1 |
| AGL17 | TaAGL17.5D2 | TraesCS7D01G107200 | chr7D:64515345..64528197 | PF00319; PF01486 | FGENESH+ | OsMADS57 |
| Bsister | TaBS.3D | TraesCS7D01G633200LC | chr7D:578483710..578486060 | PF00319; PF01486 | FGENESH_C | TRIAE_CS42_ |

**Table S5. Primer sequences for RT-PCR.**

| Primer Name | Primer Sequence |
|---|---|
| TaAGL41D_1F | 5'-TTCAAGAAGGCGTTCGAGCT-3' |
| TaAGL41D_R | 5'-TCCTCGCTTCTTCTTTCCCC-3' |
| TaMADS2B_1F | 5'-GGATCGAGGACCGGACGA-3' |
| TaMADS2B_R | 5'- CCCCTAGGACGACTGGAGTT-3' |

**Table S6. Expression modules and gene numbers for Figure 5A and S3.**

| No in heatmap | Module | Module Description | color Fig 5 | clade | Name | IWGSC ID |
|---|---|---|---|---|---|---|
| 1 | 10 | medium in spikelet; low in leaf, root | | MIKC* | TaMIKC.2A | TraesCS4A01G290700 |
| 2 | 10 | medium in spikelet; low in leaf, root | | MIKC* | TaMIKC.2B | TraesCS4B01G023300 |
| 3 | 10 | medium in spikelet; low in leaf, root | | MIKC* | TaMIKC.2D | TraesCS4D01G021100 |
| 4 | 10 | medium in spikelet; low in leaf, root | | MIKC* | TaMIKC.1A | TraesCS4A01G169000 |
| 5 | 10 | medium in spikelet; low in leaf, root | | MIKC* | TaMIKC.1B | TraesCS4B01G150200 |
| 6 | 10 | medium in spikelet; low in leaf, root | | MIKC* | TaMIKC.1D | TraesCS4D01G147000 |
| 7 | 9 | low or no expression | | OsMADS32 | TaMADS32B | TraesCS3B01G318300 |
| 8 | 14 | low in root, leaf, grain; medium in spikelet | | OsMADS32 | TaMADS32A | TraesCS3A01G284400 |
| 9 | 14 | low in root, leaf, grain; medium in spikelet | | OsMADS32 | TaMADS32D | TraesCS3D01G284200 |
| 10 | 6 | medium to high in spikelet; low in grain | | SEP1 | TaSEP1.1A | TraesCS4A01G078700 |
| 11 | 6 | medium to high in spikelet; low in grain | | SEP1 | TaSEP1.1B | TraesCS4B01G245800 |
| 12 | 6 | medium to high in spikelet; low in grain | | SEP1 | TaSEP1.1D | TraesCS4D01G245200 |
| 13 | 3 | medium in root, leaf, spikelet, grain | | SEP1 | TaSEP1.2D | TraesCS4D01G243700 |
| 14 | 6 | medium to high in spikelet; low in grain | | SEP1 | TaSEP1.2A | TraesCS4A01G058900 |
| 15 | 9 | low or no expression | | SEP1 | TaSEP1.2B | TraesCS4B01G245700 |
| 16 | 6 | medium to high in spikelet; low in grain | | SEP1 | TaSEP1.3D | TraesCS4D01G276100 |
| 17 | 9 | low or no expression | | SEP1 | TaSEP1.3A2 | TraesCS4A01G028200 |
| 18 | 9 | low or no expression | | SEP1 | TaSEP1.3B | TraesCS4B01G277800 |
| 19 | 10 | medium in spikelet; low in leaf, root | | SEP1 | TaSEP1.3A1 | TraesCS4A01G028100 |
| 20 | 15 | medium in root, leaf, grain; high in spikelet | | SEP1 | TaSEP1.6A | TraesCS5A01G391800 |
| 21 | 15 | medium in root, leaf, grain; high in spikelet | | SEP1 | TaSEP1.6B | TraesCS5B01G396700 |
| 22 | 15 | medium in root, leaf, grain; high in spikelet | | SEP1 | TaSEP1.6D | TraesCS5D01G401700 |
| 23 | 6 | medium to high in spikelet; low in grain | | SEP1 | TaSEP1.5A | TraesCS7A01G122100 |
| 24 | 6 | medium to high in spikelet; low in grain | | SEP1 | TaSEP1.5B1 | TraesCS7B01G020900 |
| 25 | 6 | medium to high in spikelet; low in grain | | SEP1 | TaSEP1.5D | TraesCS7D01G120600 |
| 26 | 9 | low or no expression | | SEP1 | TaSEP1.5B2 | TraesCS7B01G021100 |
| 27 | 1 | high in root, leaf, spikelet; low in grain | | SEP1 | TaSEP1.4D | TraesCS7D01G120500 |
| 28 | 15 | medium in root, leaf, grain; high in spikelet | | SEP1 | TaSEP1.4A | TraesCS7A01G122000 |
| 29 | 15 | medium in root, leaf, grain; high in spikelet | | SEP1 | TaSEP1.4B | TraesCS7B01G020800 |
| 30 | 16 | high in spikelet, grain | | SEP3 | TaSEP3.1A | TraesCS7A01G260600 |
| 31 | 16 | high in spikelet, grain | | SEP3 | TaSEP3.1B | TraesCS7B01G158600 |
| 32 | 16 | high in spikelet, grain | | SEP3 | TaSEP3.1D | TraesCS7D01G261600 |
| 33 | 16 | high in spikelet, grain | | SEP3 | TaSEP3.2A | TraesCS5A01G286800 |
| 34 | 16 | high in spikelet, grain | | SEP3 | TaSEP3.2B | TraesCS5B01G286100 |
| 35 | 16 | high in spikelet, grain | | SEP3 | TaSEP3.2D | TraesCS5D01G294500 |
| 36 | 7 | medium in spikelet, grain | | AGL6 | TaAGL6A | TraesCS6A01G259000 |
| 37 | 7 | medium in spikelet, grain | | AGL6 | TaAGL6B | TraesCS6B01G286400 |
| 38 | 7 | medium in spikelet, grain | | AGL6 | TaAGL6D | TraesCS6D01G240200 |
| 39 | 1 | high in root, leaf, spikelet; low in grain | | AP1 | TaAP1.1A | TraesCS5A01G391700 |
| 40 | 1 | high in root, leaf, spikelet; low in grain | | AP1 | TaAP1.1B | TraesCS5B01G396600 |
| 41 | 15 | medium in root, leaf, grain; high in spikelet | | AP1 | TaAP1.1D | TraesCS5D01G401500 |
| 42 | 1 | high in root, leaf, spikelet; low in grain | | AP1 | TaAP1.3A | TraesCS2A01G261200 |
| 43 | 1 | high in root, leaf, spikelet; low in grain | | AP1 | TaAP1.3B | TraesCS2B01G281000 |
| 44 | 1 | high in root, leaf, spikelet; low in grain | | AP1 | TaAP1.3D | TraesCS2D01G262700 |
| 45 | 1 | high in root, leaf, spikelet; low in grain | | AP1 | TaAP1.2A | TraesCS2A01G174300 |
| 46 | 1 | high in root, leaf, spikelet; low in grain | | AP1 | TaAP1.2B | TraesCS2B01G200800 |
| 47 | 1 | high in root, leaf, spikelet; low in grain | | AP1 | TaAP1.2D | TraesCS2D01G181400 |
| 48 | 11 | medium in spikelet, grain | | AG/STK | TaSTK.1B | TraesCS5B01G115100 |
| 49 | 16 | high in spikelet, grain | | AG/STK | TaSTK.1A | TraesCS5A01G117500 |
| 50 | 16 | high in spikelet, grain | | AG/STK | TaSTK.1D | TraesCS5D01G118200 |
| 51 | 11 | medium in spikelet, grain | | AG/STK | TaSTK.2A | TraesCS1A01G262700 |
| 52 | 11 | medium in spikelet, grain | | AG/STK | TaSTK.2B | TraesCS1B01G273300 |
| 53 | 11 | medium in spikelet, grain | | AG/STK | TaSTK.2D | TraesCS1D01G262700 |
| 54 | 5 | high in spikelet | | AG/STK | TaAG.2A | TraesCS3A01G314300 |
| 55 | 5 | high in spikelet | | AG/STK | TaAG.2B | TraesCS3B01G157500 |
| 56 | 10 | medium in spikelet; low in leaf, root | | AG/STK | TaAG.2D | TraesCS3D01G140200 |
| 57 | 16 | high in spikelet, grain | | AG/STK | TaAG.1A | TraesCS1A01G125800 |
| 58 | 16 | high in spikelet, grain | | AG/STK | TaAG.1D | TraesCS1B01G144800 |
| 59 | 16 | high in spikelet, grain | | AG/STK | TaAG.1B | TraesCS1D01G127700 |
| 60 | 8 | medium in root | | AGL12 | TaAGL12.1A | TraesCS7A01G319400 |
| 61 | 17 | predominantly root; medium low in leaf | | AGL12 | TaAGL12.1B | TraesCS7B01G220200 |
| 62 | 17 | predominantly root; medium low in leaf | | AGL12 | TaAGL12.1D | TraesCS7D01G315900 |
| 63 | 10 | medium in spikelet; low in leaf, root | | AGL12 | TaAGL12.2D | TraesCS2D01G309300 |
| 64 | 13 | medium in root, spikelet, grain; medium low in leaf | | AGL12 | TaAGL12.2A | TraesCS2A01G311100 |
| 65 | 13 | medium in root, spikelet, grain; medium low in leaf | | AGL12 | TaAGL12.2B | TraesCS2B01G328000 |
| 66 | 8 | medium in root | | AGL17 | TaAGL17.10U | TraesCSU01G389400LC |
| 67 | 9 | low or no expression | | AGL17 | TaAGL17.11A1 | TraesCS5A01G001400LC |
| 68 | 9 | low or no expression | | AGL17 | TaAGL17.10B | TraesCS5B01G002500 |
| 69 | 9 | low or no expression | | AGL17 | TaAGL17.10D | TraesCS5D01G002100 |
| 70 | 9 | low or no expression | | AGL17 | TaAGL17.11A3 | TraesCS7A01G151800LC |
| 71 | 9 | low or no expression | | AGL17 | TaAGL17.11B1 | TraesCS7B01G009600 |
| 72 | 9 | low or no expression | | AGL17 | TaAGL17.11B2 | TraesCS7B01G009900 |
| 73 | 9 | low or no expression | | AGL17 | TaAGL17.11D2 | TraesCS7D01G107000 |
| 74 | 9 | low or no expression | | AGL17 | TaAGL17.11D1 | TraesCS7D01G135600LC |
| 75 | 9 | low or no expression | | AGL17 | TaAGL17.2A3 | TraesCS5A01G001300LC |
| 76 | 9 | low or no expression | | AGL17 | TaAGL17.2A2 | TraesCS5A01G001800 |
| 77 | 9 | low or no expression | | AGL17 | TaAGL17.2B | TraesCS5B01G002400 |
| 78 | 9 | low or no expression | | AGL17 | TaAGL17.2D | TraesCS5D01G002200 |
| 79 | 9 | low or no expression | | AGL17 | TaAGL17.2A1 | TraesCSU01G209900 |
| 80 | 9 | low or no expression | | AGL17 | TaAGL17.4A | TraesCS7A01G111700 |
| 81 | 9 | low or no expression | | AGL17 | TaAGL17.4B | TraesCS7B01G009500 |
| 82 | 10 | medium in spikelet; low in leaf, root | | AGL17 | TaAGL17.4D | TraesCS7D01G106600 |
| 83 | 9 | low or no expression | | AGL17 | TaAGL17.5A1 | TraesCS7A01G111800 |
| 84 | 9 | low or no expression | | AGL17 | TaAGL17.5A2 | TraesCS7A01G111900 |
| 85 | 9 | low or no expression | | AGL17 | TaAGL17.5B | TraesCS7B01G009800 |
| 86 | 9 | low or no expression | | AGL17 | TaAGL17.5D1 | TraesCS7D01G106900 |
| 87 | 9 | low or no expression | | AGL17 | TaAGL17.5D2 | TraesCS7D01G107200 |
| 88 | 8 | medium in root | | AGL17 | TaAGL17.3B | TraesCS6B01G322700 |
| 89 | 9 | low or no expression | | AGL17 | TaAGL17.3A | TraesCS6A01G292300 |
| 90 | 13 | medium in root, spikelet, grain; medium low in leaf | | AGL17 | TaAGL17.3D | TraesCS6D01G273500 |
| 91 | 9 | low or no expression | | AGL17 | TaAGL17.6B2 | TraesCS7B01G793900LC |
| 92 | 9 | low or no expression | | AGL17 | TaAGL17.6B3 | TraesCS7B01G794200LC |
| 93 | 9 | low or no expression | | AGL17 | TaAGL17.6U1 | TraesCSU01G169600LC |
| 94 | 9 | low or no expression | | AGL17 | TaAGL17.6U2 | TraesCSU01G236400LC |
| 95 | 9 | low or no expression | | AGL17 | TaAGL17.6U3 | TraesCSU01G236600LC |
| 96 | 9 | low or no expression | | AGL17 | TaAGL17.7B1 | TraesCS1B01G058200 |
| 97 | 9 | low or no expression | | AGL17 | TaAGL17.7D | TraesCS1D01G045600 |
| 98 | 9 | low or no expression | | AGL17 | TaAGL17.8A | TraesCS1A01G045000 |

| | | | | | |
|---|---|---|---|---|---|
| 99 | 9 | low or no expression | AGL17 | TaAGL17.8B | TraesCS1B01G058300 |
| 100 | 9 | low or no expression | AGL17 | TaAGL17.8D | TraesCS1D01G045700 |
| 101 | 8 | medium in root | AGL17 | TaAGL17.9B | TraesCS1B01G058100 |
| 102 | 8 | medium in root | AGL17 | TaAGL17.9D | TraesCS1D01G045500 |
| 103 | 9 | low or no expression | AGL17 | TaAGL17.9A | TraesCS1A01G044900 |
| 104 | 8 | medium in root | AGL17 | TaAGL17.1B | TraesCS2B01G344000 |
| 105 | 8 | medium in root | AGL17 | TaAGL17.1D | TraesCS2D01G325000 |
| 106 | 13 | medium in root, spikelet, grain; medium low in leaf | AGL17 | TaAGL17.1A | TraesCS2A01G337900 |
| 107 | 5 | high in spikelet | AP3 | TaAP3.1A | TraesCS7A01G383800 |
| 108 | 5 | high in spikelet | AP3 | TaAP3.1B | TraesCS7B01G286600 |
| 109 | 5 | high in spikelet | AP3 | TaAP3.1D | TraesCS7D01G380300 |
| 110 | 9 | low or no expression | AP3 | TaAP3.2A | TraesCS6A01G131100 |
| 111 | 9 | low or no expression | AP3 | TaAP3.2B | TraesCS6B01G159400 |
| 112 | 9 | low or no expression | AP3 | TaAP3.2D | TraesCS6D01G120900 |
| 113 | 5 | high in spikelet | PI | TaPI.2A | TraesCS3A01G406500 |
| 114 | 5 | high in spikelet | PI | TaPI.2B | TraesCS3B01G440200 |
| 115 | 5 | high in spikelet | PI | TaPI.2D | TraesCS3D01G401700 |
| 116 | 5 | high in spikelet | PI | TaPI.1B | TraesCS1B01G275000 |
| 117 | 5 | high in spikelet | PI | TaPI.1D | TraesCS1D01G264500 |
| 118 | 16 | high in spikelet, grain | PI | TaPI.1A | TraesCS1A01G264300 |
| 119 | 11 | medium in spikelet, grain | Bsister | TaBS.1A | TraesCS6A01G158100 |
| 120 | 11 | medium in spikelet, grain | Bsister | TaBS.1B | TraesCS6B01G186700 |
| 121 | 11 | medium in spikelet, grain | Bsister | TaBS.1D | TraesCS6D01G147400 |
| 122 | 9 | low or no expression | Bsister | TaBS.2A | TraesCS2A01G422400 |
| 123 | 9 | low or no expression | Bsister | TaBS.2B | TraesCS2B01G440900 |
| 124 | 9 | low or no expression | Bsister | TaBS.2D | TraesCS2D01G418800 |
| 125 | 9 | low or no expression | Bsister | TaBS.3A | TraesCS7A01G672200LC |
| 126 | 9 | low or no expression | Bsister | TaBS.3B | TraesCS7B01G377700 |
| 127 | 9 | low or no expression | Bsister | TaBS.3D | TraesCS7D01G633200LC |
| 128 | 9 | low or no expression | Bsister | TaBS.4A1 | TraesCS3A01G090300 |
| 129 | 9 | low or no expression | Bsister | TaBS.4A2 | TraesCS3A01G090900 |
| 130 | 9 | low or no expression | Bsister | TaBS.4B | TraesCS3B01G105700 |
| 131 | 9 | low or no expression | Bsister | TaBS.4D2 | TraesCS3D01G090600 |
| 132 | 10 | medium in spikelet; low in leaf, root | Bsister | TaBS.4D1 | TraesCS3D01G090300 |
| 133 | 3 | medium in root, leaf, spikelet, grain | Bsister | TaBS.5A4 | TraesCS4A01G137500 |
| 134 | 9 | low or no expression | Bsister | TaBS.5A5 | TraesCS1A01G491400LC |
| 135 | 9 | low or no expression | Bsister | TaBS.5A3 | TraesCS2A01G153100 |
| 136 | 9 | low or no expression | Bsister | TaBS.5A2 | TraesCS2A01G475000 |
| 137 | 9 | low or no expression | Bsister | TaBS.5A1 | TraesCS7A01G475000 |
| 138 | 9 | low or no expression | Bsister | TaBS.5D | TraesCS7D01G632400LC |
| 139 | 9 | low or no expression | Bsister | TaBS.6A2 | TraesCS2A01G733800LC |
| 140 | 9 | low or no expression | Bsister | TaBS.6B2 | TraesCS2B01G665600LC |
| 141 | 10 | medium in spikelet; low in leaf, root | Bsister | TaBS.6A1 | TraesCS6A01G325500LC |
| 142 | 12 | medium low in root, leaf, spikelet, grain | Bsister | TaBS.6B1 | TraesCS3B01G236300LC |
| 143 | 9 | low or no expression | Bsister | TaBS.7B | TraesCS1B01G467500 |
| 144 | 9 | low or no expression | Bsister | TaBS.7D2 | TraesCS1D01G441200 |
| 145 | 9 | low or no expression | Bsister | TaBS.9D | TraesCS3D01G102700 |
| 146 | 9 | low or no expression | Bsister | TaBS.9A | TraesCS7A01G474900 |
| 147 | 9 | low or no expression | Bsister | TaBS.9B | TraesCS7B01G633300LC |
| 148 | 9 | low or no expression | Bsister | TaBS.8D1 | TraesCS2D01G714300LC |
| 149 | 9 | low or no expression | Bsister | TaBS.8D2 | TraesCS3D01G045700LC |
| 150 | 12 | medium low in root, leaf, spikelet, grain | Bsister | TaBS.8A | TraesCS4A01G044400LC |
| 151 | 9 | low or no expression | FLC | TaFLC.1D1 | TraesCS7D01G261900 |
| 152 | 9 | low or no expression | FLC | TaFLC.1D2 | TraesCS7D01G328100LC |
| 153 | 10 | medium in spikelet; low in leaf, root | FLC | TaFLC.1A | TraesCS7A01G260900 |
| 154 | 10 | medium in spikelet; low in leaf, root | FLC | TaFLC.1B | TraesCS7B01G158900 |
| 155 | 4 | medium in root, spikelet, grain; low in leaf | FLC | TaFLC.2B | TraesCS4B01G351500 |
| 156 | 4 | medium in root, spikelet, grain; low in leaf | FLC | TaFLC.2D | TraesCS4D01G346300 |
| 157 | 4 | medium in root, spikelet, grain; low in leaf | FLC | TaFLC.2A | TraesCS5A01G520200 |
| 158 | 3 | medium in root, leaf, spikelet, grain | FLC | TaFLC.3D | TraesCS3D01G427700 |
| 159 | 9 | low or no expression | FLC | TaFLC.3A | TraesCS3A01G434400 |
| 160 | 13 | medium in root, spikelet, grain; medium low in leaf | FLC | TaFLC.3B | TraesCS3B01G469700 |
| 161 | 9 | low or no expression | FLC | TaFLC.4A2 | TraesCS3A01G434700 |
| 162 | 9 | low or no expression | FLC | TaFLC.6A | TraesCS3A01G434900 |
| 163 | 9 | low or no expression | FLC | TaFLC.4A1 | TraesCS3A01G435000 |
| 164 | 9 | low or no expression | FLC | TaFLC.6B | TraesCS3B01G683500LC |
| 165 | 9 | low or no expression | FLC | TaFLC.4D1 | TraesCS3D01G428000 |
| 166 | 9 | low or no expression | FLC | TaFLC.5D | TraesCS3D01G514600LC |
| 167 | 10 | medium in spikelet; low in leaf, root | FLC | TaFLC.5A | TraesCS3A01G432900 |
| 168 | 10 | medium in spikelet; low in leaf, root | FLC | TaFLC.6D | TraesCS3D01G427900 |
| 169 | 10 | medium in spikelet; low in leaf, root | FLC | TaFLC.1D3 | TraesCS3D01G517300LC |
| 170 | 13 | medium in root, spikelet, grain; medium low in leaf | FLC | TaFLC.4B2 | TraesCS3B01G470000 |
| 171 | 3 | medium in root, leaf, spikelet, grain | SOC1 | TaSOC1.2D | TraesCS6D01G014600 |
| 172 | 10 | medium in spikelet; low in leaf, root | SOC1 | TaSOC1.2A | TraesCS6A01G011700 |
| 173 | 13 | medium in root, spikelet, grain; medium low in leaf | SOC1 | TaSOC1.2B | TraesCS6B01G017900 |
| 174 | 15 | medium in root, leaf, grain; high in spikelet | SOC1 | TaSOC1.3B | TraesCS4B01G346700 |
| 175 | 15 | medium in root, leaf, grain; high in spikelet | SOC1 | TaSOC1.3D | TraesCS4D01G341700 |
| 176 | 15 | medium in root, leaf, grain; high in spikelet | SOC1 | TaSOC1.3A | TraesCS5A01G515500 |
| 177 | 9 | low or no expression | SOC1 | TaSOC1.4A | TraesCS7A01G051700LC |
| 178 | 9 | low or no expression | SOC1 | TaSOC1.4D | TraesCS7D01G037000 |
| 179 | 9 | low or no expression | SOC1 | TaSOC1.5A | TraesCS1A01G199900 |
| 180 | 9 | low or no expression | SOC1 | TaSOC1.5B | TraesCS1B01G214600 |
| 181 | 9 | low or no expression | SOC1 | TaSOC1.5D | TraesCS1D01G203400 |
| 182 | 9 | low or no expression | SOC1 | TaSOC1.1B | TraesCS1B01G214500 |
| 183 | 15 | medium in root, leaf, grain; high in spikelet | SOC1 | TaSOC1.1A | TraesCS1A01G199600 |
| 184 | 15 | medium in root, leaf, grain; high in spikelet | SOC1 | TaSOC1.1D | TraesCS1D01G203300 |
| 185 | 2 | medium high in root, leaf, spikelet, grain | SVP | TaSVP.1A | TraesCS6A01G313800 |
| 186 | 2 | medium high in root, leaf, spikelet, grain | SVP | TaSVP.1B | TraesCS6B01G343900 |
| 187 | 2 | medium high in root, leaf, spikelet, grain | SVP | TaSVP.1D | TraesCS6D01G293200 |
| 188 | 2 | medium high in root, leaf, spikelet, grain | SVP | TaSVP.3A | TraesCS4A01G002600 |
| 189 | 2 | medium high in root, leaf, spikelet, grain | SVP | TaSVP.3B | TraesCS4B01G302600 |
| 190 | 2 | medium high in root, leaf, spikelet, grain | SVP | TaSVP.3D | TraesCS4D01G301100 |
| 191 | 2 | medium high in root, leaf, spikelet, grain | SVP | TaSVP.2A | TraesCS7A01G175200 |
| 192 | 2 | medium high in root, leaf, spikelet, grain | SVP | TaSVP.2B | TraesCS7B01G080300 |
| 193 | 2 | medium high in root, leaf, spikelet, grain | SVP | TaSVP.2D | TraesCS7D01G176700 |

**Table S7. Tissues for expression analysis for Figure 5A and S4.**

| row | Stage |
|---|---|
| 1 | 4_grain_ endosperm_ endosperm_3_reproductive_ Dough |
| 2 | 4_grain_ embryo proper_ embryo_3_reproductive_ Dough |
| 3 | 4_grain_ grain_ grain hard dough and ripening_3_reproductive_ 8_Hard dough |
| 4 | 4_grain_ grain_ grain hard dough and ripening_3_reproductive_ 7_Ripening |
| 5 | 4_grain_ grain_ grain milk and soft dough_3_reproductive_ 6_Soft dough |
| 6 | 4_grain_ grain_ grain milk and soft dough_3_reproductive_ 5_milk grain stage |
| 7 | 3_ spike_ lemma_ glumes_3_reproductive_ 5_milk grain stage |
| 8 | 3_ spike_ glumes_ glumes_3_reproductive_ 5_milk grain stage |
| 9 | 3_ spike_ awns_ awns_3_reproductive_ 5_milk grain stage |
| 10 | 3_ spike_ stigma & ovary_ stigma & ovary_3_reproductive_ 4_anthesis |
| 11 | 3_ spike_ anther_ anther_3_reproductive_ 4_anthesis |
| 12 | 3_ spike_ spike_ spike_3_reproductive_ 3_Full boot |
| 13 | 3_ spike_ lemma_ glumes_3_reproductive_ 2_Ear emergence |
| 14 | 3_ spike_ glumes_ glumes_3_reproductive_ 2_Ear emergence |
| 15 | 3_ spike_ awns_ awns_3_reproductive_ 2_Ear emergence |
| 16 | 3_ spike_ spikelets_ spikelets_3_reproductive_ 1_30% spike |
| 17 | 3_ spike_ spike_ spike_3_reproductive_ 1_30% spike |
| 18 | 2_ leaves/shoots_ flag leaf blade (senescence)_ flag leaf blade_3_reproductive_ Dough |
| 19 | 2_ leaves/shoots_ flag leaf blade (senescence)_ flag leaf blade_3_reproductive_ 7_Ripening |
| 20 | 2_ leaves/shoots_ shoot axis_ shoot axis_3_reproductive_ 5_milk grain stage |
| 21 | 2_ leaves/shoots_ peduncle_ peduncle_3_reproductive_ 5_milk grain stage |
| 22 | 2_ leaves/shoots_ Internode #2_ internode_3_reproductive_ 5_milk grain stage |
| 23 | 2_ leaves/shoots_ flag leaf sheath_ flag leaf sheath_3_reproductive_ 5_milk grain stage |
| 24 | 2_ leaves/shoots_ flag leaf blade_ flag leaf blade_3_reproductive_ 5_milk grain stage |
| 25 | 2_ leaves/shoots_ fifth leaf blade (senescence)_ leaf blades excl flag_3_reproductive_ 5_milk grain stage |
| 26 | 2_ leaves/shoots_ flag leaf blade night (-0.25h) 06:45_ flag leaf blade_3_reproductive_ 4_anthesis |
| 27 | 2_ leaves/shoots_ fifth leaf blade night (-0.25h) 21:45_ leaf blades excl flag_3_reproductive_ 4_anthesis |
| 28 | 2_ leaves/shoots_ shoot axis_ shoot axis_3_reproductive_ 3_Full boot |
| 29 | 2_ leaves/shoots_ leaf ligule_ leaf ligule_3_reproductive_ 3_Full boot |
| 30 | 2_ leaves/shoots_ flag leaf sheath_ flag leaf sheath_3_reproductive_ 3_Full boot |
| 31 | 2_ leaves/shoots_ flag leaf blade_ flag leaf blade_3_reproductive_ 3_Full boot |
| 32 | 2_ leaves/shoots_ peduncle_ peduncle_3_reproductive_ 2_Ear emergence |
| 33 | 2_ leaves/shoots_ Internode #2_ internode_3_reproductive_ 2_Ear emergence |
| 34 | 2_ leaves/shoots_ flag leaf sheath_ flag leaf sheath_3_reproductive_ 2_Ear emergence |
| 35 | 2_ leaves/shoots_ flag leaf blade_ flag leaf blade_3_reproductive_ 2_Ear emergence |
| 36 | 2_ leaves/shoots_ fifth leaf blade_ leaf blades excl flag_3_reproductive_ 2_Ear emergence |
| 37 | 2_ leaves/shoots_ peduncle_ peduncle_3_reproductive_ 1_30% spike |
| 38 | 2_ leaves/shoots_ Internode #2_ internode_3_reproductive_ 1_30% spike |
| 39 | 2_ leaves/shoots_ flag leaf sheath_ flag leaf sheath_3_reproductive_ 1_30% spike |
| 40 | 2_ leaves/shoots_ flag leaf blade_ flag leaf blade_3_reproductive_ 1_30% spike |
| 41 | 2_ leaves/shoots_ shoot axis_ shoot axis_3_reproductive_ 0_Flag leaf stage |
| 42 | 2_ leaves/shoots_ flag leaf blade night (+0.25h) 07:15_ flag leaf blade_3_reproductive_ 0_Flag leaf stage |
| 43 | 2_ leaves/shoots_ flag leaf blade night (-0.25h) 06:45_ flag leaf blade_3_reproductive_ 0_Flag leaf stage |
| 44 | 2_ leaves/shoots_ flag leaf blade_ flag leaf blade_3_reproductive_ 0_Flag leaf stage |
| 45 | 2_ leaves/shoots_ fifth leaf sheath_ leaf sheaths excl flag_3_reproductive_ 0_Flag leaf stage |
| 46 | 2_ leaves/shoots_ fifth leaf blade night (+0.25h) 22:15_ leaf blades excl flag_3_reproductive_ 0_Flag leaf stage |
| 47 | 2_ leaves/shoots_ fifth leaf blade night (-0.25h) 21:45_ leaf blades excl flag_3_reproductive_ 0_Flag leaf stage |
| 48 | 2_ leaves/shoots_ fifth leaf blade_ leaf blades excl flag_3_reproductive_ 0_Flag leaf stage |
| 49 | 2_ leaves/shoots_ shoot axis_ shoot axis_2_vegetative_ 3_tillering stage |
| 50 | 2_ leaves/shoots_ shoot apical meristem_ shoot apical meristem_2_vegetative_ 3_tillering stage |
| 51 | 2_ leaves/shoots_ first leaf sheath_ leaf sheaths excl flag_2_vegetative_ 3_tillering stage |
| 52 | 2_ leaves/shoots_ first leaf blade_ leaf blades excl flag_2_vegetative_ 3_tillering stage |
| 53 | 2_ leaves/shoots_ fifth leaf sheath_ leaf sheaths excl flag_2_vegetative_ 2_fifth leaf stage |
| 54 | 2_ leaves/shoots_ fifth leaf blade_ leaf blades excl flag_2_vegetative_ 2_fifth leaf stage |
| 55 | 2_ leaves/shoots_ third leaf sheath_ leaf sheaths excl flag_2_vegetative_ 1_three leaf stage |
| 56 | 2_ leaves/shoots_ third leaf blade_ leaf blades excl flag_2_vegetative_ 1_three leaf stage |
| 57 | 2_ leaves/shoots_ stem axis_ seedling aerial tissues_1_seedling_ seedling stage |
| 58 | 2_ leaves/shoots_ shoot apical meristem_ shoot apical meristem_1_seedling_ seedling stage |
| 59 | 2_ leaves/shoots_ first leaf sheath_ seedling aerial tissues_1_seedling_ seedling stage |
| 60 | 2_ leaves/shoots_ first leaf blade_ seedling aerial tissues_1_seedling_ seedling stage |
| 61 | 2_ leaves/shoots_ coleoptile_ seedling aerial tissues_1_seedling_ seedling stage |
| 62 | 1_roots_ roots_ roots_3_reproductive_ 1_30% spike |
| 63 | 1_roots_ roots_ roots_3_reproductive_ 0_Flag leaf stage |
| 64 | 1_roots_ roots_ roots_2_vegetative_ 3_tillering stage |
| 65 | 1_roots_ root apical meristem_ root apical meristem_2_vegetative_ 3_tillering stage |
| 66 | 1_roots_ roots_ roots_2_vegetative_ 1_three leaf stage |
| 67 | 1_roots_ root apical meristem_ root apical meristem_2_vegetative_ 1_three leaf stage |
| 68 | 1_roots_ axillary roots_ roots_2_vegetative_ 1_three leaf stage |
| 69 | 1_roots_ roots_ roots_1_seedling_ seedling stage |
| 70 | 1_roots_ radicle_ roots_1_seedling_ seedling stage |

#### The *SEP1*, *AGL17*, $B_{sister}$ and *FLC* subfamilies are significantly larger than expected

Four MIKC-type subfamilies were significantly larger than expected: *SEP1*, *AGL17*, $B_{sister}$ and *FLC*. For all four of them, additional phylogenies were generated in order to resolve gene relationships more precisely (Table 1, Figure 2, Figure S3A-D).

Within the *SEP1/SEP3* subfamily three out of five rice orthologs can be assigned to a triad of wheat homoeologs as sister clade, resulting in the expected 1:3 ratio (Figure S3A). In contrast, the *SEP1*-like rice genes *OsMADS1* and *OsMADS5* are sister to 10 (3 + 3 + 4) and 7 (3 + 4) wheat genes, respectively (Figure S3A). The phylogenetic relationship and conserved chromosomal position of the genes within each of the subclades (all genes on chromosomes 4 and 7, respectively) suggests duplications in the lineage leading to *Triticum*, but before hexaploidization. Subsequently, after the initial specification of the three donor species, tandem duplications might have generated a fourth gene in two cases (*TaSEP1.3A* and *TaSEP1.5B*; Figure S3A). All but two *SEP1*-like genes maintained a M-I-K-C structure (Table 1). Exceptions are two *OsMADS1*-like genes in the A subgenome, lacking a K- and a MADS-box, respectively (*TaSEP1.1A* and *TaSEP1.2A*; Figure 1, Figure S3A, Table S2).

In rice, there are three paralogous *Bsister* genes: *OsMADS29*, *OsMADS30* and *OsMADS31*. In wheat, *OsMADS29*-like and *OsMADS31*-like genes were found in the expected 1:3 ratios, in syntenic locations and having a canonical M-I-K-C structure (Figure 2E, Figure S3B). In contrast, the third rice *Bsister* gene *OsMADS30* had 27 homologs, accounting for the skewed rice-to-wheat gene ratio of the *Bsister* clade (Table 1, Figure S3B). However, only half of the 27 *OsMADS30*-like genes possessed a canonical M-I-K-C structure (13 genes, 48 %), the remaining encoded either solely for a MADS-domain (37 %, 10 genes) or a K-domain (15 %, 4 genes) (Table1, Figure S3B). Closely related genes often shared similarly truncated gene structures (Figure S3B); however, they often were found on different non-homoeologous chromosomes in non-syntenic locations. This could be pointing towards transposable elements as a putative mechanism of the duplication of *OsMADS30*-like wheat genes; more precisely, the "capture" of only part of a full-length gene (either MADS- or K-box) and then the subsequent transposition of this sequence to another location.

In rice, there are six *AGL17*-like genes, as compared to 47 in wheat (Table 1). Almost three quarter of the wheat *AGL17*-like genes encoded for a MADS- as well as a K-domain (72 %, 34 genes), whilst 12 and 1 gene only had a MADS- or a K-box, respectively (26 and 2 %) (Table 1, Table S2, Figure S3C). *OsMADS57* and *OsMADS61* only have the expected three homoeologs and *OsMADS27* and *OsMADS23* were not directly sister to wheat genes (Figure S2C). In contrast, *OsMADS25* and *OsMADS59* had 13 and 15 homologs, respectively (Figure S2C). One distinct subclade of *AGL17*-like wheat genes (*TaAGL17.10* and *.11*, 13 genes) could not be assigned a direct rice ortholog. Almost half of all *AGL17*-like genes were located on chromosomes 7 (43 %, 20 genes) (Figure 2E). Given their close proximity, these genes originated likely from tandem duplications.

Wheat has 20 *FLC*-like genes, which is considerably higher than the two *FLC*-like genes from rice (Figure 2D, Figure S3D). Phylogenetic relationships within the *FLC*-subfamily were not very well resolved. A majority of wheat *FLC*-like genes were located on chromosomes 3, all on the respective long arms of the chromosomes and in close vicinity to each other, again pointing towards tandem duplication as a possible mechanism of gene duplication (65 % of genes, Figure 2E, Figure S3D).

### Supplementary Text 2
### Masked Alignment of MADS-domain MIKC-type proteins

```
>GOA_AT1G31140
MRKGKRVIKKIEEKIKRQVTFAKRKKSLIKKAYELSVLCDVHLGLIIFSHSNRLYDFCSTSMENLIMRYQKEKEGQ--
TTAFHSCSDCVKTKESMMREIENLKLNLQLYDGHGLNLLTYDELLSFELHLESSLQHARARKSEFMHQQQQQTDQLKGKEKGGSSWEQLMWQAERQMMT-------
-----------------CQRWG
>AGL65_AT1G18750
MGRVKLKIKRLESTSNRQVTYTKRKNGILKKAKELSILCDIDIVLLMFSPTGRATAFHGSCIEEVISKFAQLTPQERT---
LESLEGLSNQVAIYQAQLMECHRRLS-CWTNIDRIENTEHLDLLEESLRKSIERIQIHKEHYRKNQLLPIECATTQFHSG------QLPMM------DNDHQQ--
QEKKIKSEQPSMYQ------M
>AGL104_AT1G22130
MGRVKLEIKRIENTTNRQVTFSKRRNGLIKKAYELSILCDIDIALIMFSPSDRLSLFSGTRIEDVFSRFINLPKQERESAYIQNKEELEHEVCRLQQQLQMAEEELR
RYEPDPIRFTTMEEYEVSEKQLLDTLTHVVQRRDHLMSNHLSSYEASTM--QPN------G----------ENGTNQTHNQNN-MSEM----GVGSIETK
>SEP4_AT1G24260
MGRGRVELKRIENKINRQVTFAKRRNGLLKKAYELSVLCDAEVALIIFSNRGKLYEFCSSSMLRTLERYQKCNYGAPEPNPALAVESSQQEYLKLKERYDALQRTQR
NLLGEDLGPLSTKELESLERQLDSSLKQIRALRTQFMLDQLNDLQSKERMLTETNKTLRLRLADGYQMPL-NQEEVDHYHSQAFFQP-EPILQIGY---M
>CAL_At1g26310
MGRGRVELKRIENKINRQVTFSKRRTGLLKKAQEISVLCDAEVSLIVFSHKGKLFEYSSSCMEKVLERYERYSYAERQLIPSHVNTNWSMEYSRLKAKIELLERNQR
HYLGEELEPMSLKDLQNLEQQLETALKHIRSRKNQLMNESLNHLQRKEKEIQEENSMLTKQIKERENIL----QLNRSVPHLYMIAH-------S--EDQ
>AP1_At1g69120
MGRGRVQLKRIENKINRQVTFSKRRAGLLKKAHEISVLCDAEVALVVFSHKGKLFEYSTSCMEKILERYERYSYAERQLIPSDVNTNWSMEYNRLKAKIELLERNQR
HYLGEDLQAMSPKELQNLEQQLDTALKHIRTRKNQLMYESINELQKKEKAIQEQNSMLSKQIKEREKIL----QQNQGHQHPYMLSH-------S--DDP
>AGL94_AT1G69540
MGRVKLKIKKLQNMNGRQCTYTKRRHGIMKKAKELSILCDIDVVLLMFSPMGKASICIGHSIGEVIAKFAQLSPQERA---
LENLEVLSEKIRFLQTQLSDIHTRLS-YWTDVDNIDSVDVLQQLEHSLRQSLAQIYGRKASMPQQQLMSSQCKN-QLQTE------DIDFM------TDENMN----
SDILQKSLSFLNLLSPGKN
>AGL12_At1g71692
MARGKIQLKRIENPVHRQVTFCKRRTGLLKKAKELSVLCDAEIGVVIFSPQGKLFELATGTMEGMIDKYMKCTGGGRGSSSQPPNLDPKDEINVLKQEIEMLQKGIS
YMFGGGDGAMNLEELLLLEKHLEYWISQIRSAKMDVMLQEIQSLRNKEGVLKNTNKYLLEKIEENNDANFA--ET--NYSYPLTMPSI-------FQ---
>FLM_AT1G77080
------------------------------------------------------MHFCFSSISKIIDRYEIQHADELRAL---DLEEKIQNYLPHK---
ELLETVQSKLEEPNVDNVSVDSLISLEEQLETALSVSRARKAELMMEYIESLKEKEKLLREENQVLASQVT------------------------------------
M
>AGL67_AT1G77950
MGRVKLELKRIEKSTNRQITFSKRKKGLIKKAYELSTLCDIDLALLMFSPSDRLCLFSGTRIEDVLARYINLPDQERENAVIQNKEELEQEVCRLQQQLQISEEELR
KFEPDPMRLTSMEEIEACEANLINTLTRVVQRREHLLR---KSCEAQSN--QQS------D----------EPEPKQAHNQN------------------
>AGL66_AT1G77980
MGRVKLEIKRIENTTNRQVTFSKRRNGLIKKAYELSILCDIDIALLMFSPSDRLSLFSGTRIEDVFSRYINLSDQERENAVFQSKEELEHEVYKLQQQLLMAEEELR
KYEPDPIRFTTMEEYETCEKQLMDTLTRVNQRREHILSDQLSSYEASALQQQQS------G----------ENGPNEAHNQNNIMGEMHN--VVDNIDIR
>AGL30_AT2G03060
MGRVKLKIKKLENTNGRQSTFAKRKNGILKKANELSILCDIDIVLLMFSPTGKAAICCGSSMEEVIAKFSQVTPQERT---
FESLEDLSTQARILQARISEIHGRLS-YWTEPDKINNVEHLGQLEISIRQSLDQLRAHKEHFGQQQAMQIENANFSMQDG------QIP--------NSNTTN--
QETSFLDETITAYQTLPPQTR
>SEP3_AT2G03710
MGRGKVELKRIENKINRQVTFAKRRNGLLKKAYELSVLCDAEIALLIFSNRGKLYEFCSSGMARTVDKYRKHSYATMD--
PAKDLQDKYQDYLKLKSRVEILQHSQRHLLGEELSEMDVNELEHLERQVDASLRQIRSTKARSMLDQLSDLKTKEEMLLETNRDLRRKV------------------
AIGIFYR-KPFVI-------
>ANR1_At2g14210
MGRGKIVIRRIDNSTSRQVTFSKRRSGLLKKAKELSILCDAEVGVIIFSSTGKLYDYASNSMKTIIERYNRVKEEQHQLLHASEIKFWQREVASLQQQLQYLQECHR
KLVGEELSGMNANDLQNLEDQLVTSLKGVRLKKDQLMTNEIRELNRKGQIIQKENHELQNIVDIMRKENIKRTNAIEGNTTTYA----PPQLQLIQL---
>SVP_AT2G22540
MAREKIQIRKIDNATARQVTFSKRRRGLFKKAEELSVLCDADVALIIFSSTGKLF-----DMKEVLERHNLQSKNEKLD-
PSLELQLENSDHARMSKEIADKSHRLRQMRGEELQGLDIEELQQLEKALETGLTRVIETKSDKIMSEISELQKKGMQLMDENKRLRQQGTQLTEENEREGQSSESIG
NSTG--ASDTSLRLGLPYGG
>AGL17_At2g22630
MGRGKIVIQKIDDSTSRQVTFSKRRKGLIKKAKELAILCDAEVCLIIFSNTDKLYDFASSSVKSTIERFNTAKMEEQELMPASEVKFWQREAETLRQELHSLQENYR
QLTGVELNGLSVKELQNIESQLEMSLRGIRMKREQILTNEIKELTRKRNLVHHENLELSRKVQRIHQENVETSNT----LGHHELVDHQVRLQLSQPSHY
>SHP2_AT2G42830
IGRGKIEIKRIENTTNRQVTFCKRRNGLLKKAYELSVLCDAEVALVIFSTRGRLYEYANNSVRGTIERYKKACSDAVNPPTANT-
QYYQQEASKLRRQIRDIQNLNRHILGESLGSLNFKELKNLESRLEKGISRVRSKKHEMLVAEIEYMQKREIELQNDNMYLRSKITERTQQESSHQGT--VY--
NQNSSNQQPPLQLV-----
>AGL6_At2g45650
MGRGRVEMKRIENKINRQVTFSKRRNGLLKKAYELSVLCDAEVALIIFSSRGKLYEFGS-GIESTIERYNRCYNCSLSNN-
EETTQSWCQEVTKLKSKYESLVRTNRNLLGEDLGEMGVKELQALERQLEAALTATRQRKTQVMMEEMEDLRKKERQLGDINKQLKIKFETEGAFKTFNSAASVA-
SHPN---VNEPFLQIGF-KSN
>SOC1_AT2G45660
MVRGKTQMKRIENATSRQVTFSKRRNGLLKKAFELSVLCDAEVSLIIFSPKGKLYEFASSNMQDTIDRYLRHTKDVSTKPSEENMQHLKYEAANMMKKIEQLEASKR
KLLGEGIGTCSIEELQQIEQQLEKSVKCIRARKTQVFKEQIEQLKQKEKALAAENEKLSEKWGSHESEVWSES--SPS---------EETQLFIGLPSRK
>SEP2_At3g02310
MGRGRVELKRIENKINRQVTFAKRRNGLLKKAYELSVLCDAEVSLIVFSNRGKLYEFCSSNMLKTLERYQKCSYGSIEVN-
AKELENSYREYLKLKGRYENLQRQQRNLLGEDLGPLNSKELEQLERQLDGSLKQVRCIKTQYMLDQLSDLQGKEHILLDANRALSMKLEDMIGVRHHGGQQNIAYHS
QGLYQS-DPTLQIGY--VQ
>AGL79_At3g30260
MGRGRVQLRRIENKIRRQVTFSKRRTGLVKKAQEISVLCDAEVALIVFSPKGKLFEYSASSMERILDRYERSAYAGQDIPPLDSQGECSTECSKLLRMIDVLQRSLR
HLRGEEVDGLSIRDLQGVEMQLDTALKKTRSRKNQLMVESIAQLQKKEKELKELKKQLTKKAGEREDFQTQ--------ESPHELRR-PPPPPLS--AAG
>AP3_AT3G54340
MARGKIQIKRIENQTNRQVTYSKRRNGLFKKAHELTVLCDARVSIIMFSSSNKLHEYISTTTKEIVDLYQTISDVD-----
ATQYERMQETKRKLLETNRNLRTQIKQRLGECLDELDIQELRRLEDEMENTFKLVRERKFKSLGNQIETTKKKNKSQQDIQKNLIHELELREDPHYGDN------
NHHHYYPNAIITFHL-LE---
>AGL16_At3g57230
MGRGKIAIKRINNSTSRQVTFSKRRNGLLKKAKELAILCDAEVGVIIFSSTGRLYDFSSSSMKSVIERYSDAKGETSSENPASEIQFWQKEAAILKRQLHNLQENHR
QMMGEELSGLSVEALQNLENQLELSLRGVRMKKDQMLIEEIQVLNREGNLVHQENLDLHKKVNLMHQQNMEVEGVKIANLLTNGLDMNHVHLQLSQPDHE
>AGL18_At3g57390
-------------------------------------------------------------MEQILSRYGYTTASTE----
REQLEAVLRNDDSMKGELERLQLAIERLKGKELEGMSFPDLISLENQLNESLHSVKDQKTQILLNQIERSRIQEKKALEENQILRKQVEML------QDSSPEAD--
-----EHDTSLQLGLSDNS
```

>SHP1_AT3G58780
LGRGKIEIKRIENTTNRQVTFCKRRNGLLKKAYELSVLCDAEVALVIFSTRGRLYEYANNSVRGTIERYKKACSDAVNPPSANT-
QYYQQEASKLRRQIRDIQNSNRHIVGESLGSLNFKELKNLEGRLEKGISRVRSKKNELLVAEIEYMQKREMELQHNNMYLRAKIAEGAQQESSQGTT--VY--
NQQFSGQQPPLQLV-----
>AGL13_AT3G61120
MGRGKVEVKRIENKITRQVTFSKRKSGLLKKAYELSVLCDAEVSLIIFSTGGKLYEFSN-GVGRTIERYYRCKDNLLDND-
LEDTQGLRQEVTKLKCKYESLLRTHRNLVGEDLEGMSIKELQTLERQLEGALSATRKQKTQVMMEQMEELRRKERELGDINNKLKLETEDHD-FKGFNPVLTAG-
THQNYIS-NGYFLQIGF-KSN
>STK_AT4G09960
MGRGKIEIKRIENSTNRQVTFCKRRNGLLKKAYELSVLCDAEVALIVFSTRGRLYEYANNNIRSTIERYKKACSDSTNTSTINA-
AYYQQESAKLRQQIQTIQNSNRNLMGDSLSSLSVKELKQVENRLEKAISRIRSKKHELLLVEIENAQKREIELDNENIYLRTKVAEVEQHHHQSGS---
EIGNGGSYSD-KKILHLG-----
>XAL2_AT4G11880
MVRGKTEMKRIENATSRQVTFSKRRNGLLKKAFELSVLCDAEVALIIFSPRGKLYEFSSSSIPKTVERYQKRIQDLGSNHRNDNSQQSKDETYGLARKIEHLEISTR
KMMGEGLDASSIEELQQLENQLDRSLMKIRAKKYQLLREETEKLKEKERNLIAENKMLMEKCEMQGRGIIGLDIDDNE---------EVTDLFIGPPRHF
>AG_AT4G18960
SGRGKIEIKRIENTTNRQVTFCKRRNGLLKKAYELSVLCDAEVALIVFSSRGRLYEYSNNSVKGTIERYKKAISDNSNTGSINA-
QYYQQESAKLRQQIISIQNSNRQLMGETIGSMSPKELRNLEGRLERSITRIRSKKNELLFSEIDYMQKREVDLHNDNQILRAKIAENEPSISLGGS---NY-
NNHHYSSQQTALQLV-----
>AGL19_At4g22950
MVRGKTEMKRIENATSRQVTFSKRRNGLLKKAFELSVLCDAEVALVIFSPRSKLYEFSSSSIAATIERYQRRIKEIGNNHRNDNSQQARDETSGLTKKIEQLEISKR
KLLGEGIDACSIEELQQLENQLDRSLSRIRAKKYQLLREEIEKLKAEERNLVKENKDLKEKWLGMGTATIAVNIDDN----------EETGLFIGPPRQS
>AGL24_AT4G24540
MAREKIRIKKIDNITARQVTFSKRRRGIFKKADELSVLCDADVALIIFSATGKLFEFSSSRMRDILGRYSLHASNNKLMDPSTHLRLENCNLSRLSKEVEDKTKQLR
KLRGEDLDGLNLEELQRLEKLLESGLSRVSEKKGECVMSQIFSLEKRGSELVDENKRLRDKLETLERAKL-EALETESVVSSYDS-GDDTSLKLGLPSWE
>AGL21_At4g37940
MGRGKIVIQRIDDSTSRQVTFSKRRKGLIKKAKELAILCDAEVGLIIFSSTGKLYDFASSSMKSVIDRYNKSKIEQQQLLPASEVKFWQREAAVLRQELHALQENHR
QMMGEQLNGLSVNELNSLENQIEISLRGIRMRKEQLLTQEIQELSQKRNLIHQENLDLSRKVQRIHQENVEANT-----FTHREVAVHQIRLQLSQPSDY
>FLC_AT5G10140
MGRKKLEIKRIENKSSRQVTFSKRRNGLIEKARQLSVLCDASVALLVVSASGKLYSFSSDNLVKILDRYGKQHADDLKAL---DHQSKALNYGSHY---
ELLELVDSKLVGSNVKNVSIDALVQLEEHLETALSVTRAKKTELMLKLVENLKEKEKMLKEENQVLASQMEN-N-------------------------------E-
M
>AGL15_At5g13790
MGRGKIEIKRIENANSRQVTFSKRRSGLLKKARELSVLCDAEVAVIVFSKSGKLFEYSSTGMKQTLSRYGNHQSSS-----
ASKAEEDCAEVDILKDQLSKLQEKHLQLQGKGLNPLTFKELQSLEQQLYHALITVRERKERLLTNQLEESRLKEQRAELENETLRRQVQELR-----PS--------
-----NDDTTLQLGLPTNT
>SEP1_At5g15800
MGRGRVELKRIENKINRQVTFAKRRNGLLKKAYELSVLCDAEVALIIFSNRGKLYEFCSSNMLKTLDRYQKCSYGSIEVN-
AKELENSYREYLKLKGRYENLQRQQRNLLGEDLGPLNSKELEQLERQLDGSLKQVRSIKTQYMLDQLSDLQNKEQMLLETNRALAMKLDDMIGVRSHGGEQNVTYQS
QGLYQP-NPTLQMGY--TQ
>PI_AT5G20240
MGRGKIEIKRIENANNRVVTFSKRRNGLVKKAKEITVLCDAKVALIIFASNGKMIDYCCMDLGAMLDQYQKLSGKK-----
DAKHENLSNEIDRIKKENDSLQLELRHLKGEDIQSLNLKNLMAVEHAIEHGLDKVRDHQMEILISK----RRNEKMMAEEQRQLTFQLQ---QQEMASNARGMMM--
-------QFGYRV-QPQEK
>ABS_AT5G23260
MGRGKIEIKKIENQTARQVTFSKRRTGLIKKTRELSILCDAHIGLIVFSATGKLSEFCSNRMPQLIDRYLHTNGLR-D---
HDDQEQLHHEMELLRRETCNLELRLRPFHGHDLASIPPNELDGLERQLEHSVLKVRERKNELMQQQLENLSRKRRMLEEDNNNMYRWLH---
EHRAAFQQAGIDTGEYQQFLEQNSVLQLALPQND
>AGL72_At5g51860
MVRGKIEIKKIENVTSRQVTFSKRRSGLFKKAHELSVLCDAQVAAMIFSQKGRLYEFASSDIRNTIKRYAEYKREVAETHIEQYVQGLKKEMVTMVKKIEVLEVHNR
KMMGQSLDSCSVKELSEIATQIEKSLHMVRLRKAKLYEDELQKLKAKERELKDERVRLSLKVGERPMGMP-GS--KEK---------DETDLFIGFLRP-
>AGL71_At5g51870
MVRGKIEIKKIENVTSRQVTFSKRRSGLFKKAHELSVLCDAQVAAIVFSQSGRLHEYSSSQMEKIIDRYGKFSNAVAERPVERYLQELKMEIDRMVKKIDLLEVHHR
KLLGQGLDSCSVTELQEIDTQIEKSLRIVRSRKAELYADQLKKLKEKERELLNERKRLLEEQNRERLMRP-RT--KHS---------EETDLFIGLPRL-
>AGL8_At5g60910
MGRGRVQLKRIENKINRQVTFSKRRSGLLKKAHEISVLCDAEVALIVFSSKGKLFEYSTSCMERILERYDRYLYSDKQLVRVSQSENWVLEHAKLKARVEVLEKNKR
NFMGEDLDSLSLKELQSLEHQLDAAIKSIRSRKNQAMFESISALQKKDKALQDHNNSLLKKIKEREKKT----QCSNSS---YCVTS-------D--GAS
>AGL42_At5g62165
MVRGKIEMKKIENATSRQVTFSKRRNGLLKKAYELSVLCDAQLSLIIFSQRGRLYEFSSSDMQKTIERYRKYTKDETSNHSQIHLQQLKQEASHMITKIELLEFHKR
KLLGQGIASCSLEELQEIDSQLQRSLGKVRER--------------KEKQLLEENVKLHQKNVINP-----QE--KYKV--------EETDLFIGLPNC-
>MAF2_At5g65050
MGRKKVEIKRIENKSSRQVTFSKRRNGLIEKARQLSILCESSIAVLVVSGSGKLYKSASDNMSKIIDRYEIHHADELEAL---DLAEKTRNYLPLK---
ELLEIVQSKLEESNVDNASVDTLISLEEQLETALSVTRARKTELMMGEVKSLQKTENLLREENQTLASQVGK-K-------------------------------G-
M
>AGL70_At5g65060
MGRRKVEIKRIENKSSRQVTFSKRRKGLIEKARQLSILCESSIAVVAVSGSGKLYDSASDNMSKIIDRYEIHHADELKAL---DLAEKIRNYLPHK---ELLEIVQ-
----------SVDSLISMEEQLETALSVIRAKKTELMMEDMKSLQEREKLLIEENQILASQVGK-K-------------------------------G-M
>AGL69_At5g65070
MGRRKVEIKRIENKSSRQVTFCKRRNGLMEKARQLSILCESSVALIIISATGRLYSFSSDSMAKILSRYELEQADDLKTL---DLEEKTLNYLSHK---
ELLETIQCKIEEAKSDNVSIDCLKSLEEQLKTALSVTRARKTELMMELVKTHQEKEKLLREENQSLTNQMGKMK-------------------------------A-
M
>AGL68_At5g65080
MGRRRVEIKRIENKSSRQVTFCKRRNGLMEKARQLSILCGSSVALFIVSSTGKLYNSSSDSMAKIISRFKIQQADDPETL---DLEDKTQDYLSHK---
ELLEIVQRKIEEAKGDNVSIESLISMEEQLKSALSVIRARKTELLMELVKNLQDKEKLLKEKNKVLASEVGKLK-------------------------------
AVM
>TaAGL17.6B1_FGENESH
MVRGKAVIEKIENQTSRQVTFSKRKSRLFKKGKELGVLCDAQVGIFIFSNTGRLYEYSNS-MKPLIERYQAVKDGQKLL-
ASAEAKFWQAETARLEQQLRTLQENHRQLLGQHLS------------------------------------------------------------------------
--------------------
>TaAGL17.10B_TraesCS5B01G002500
MARGKIVIRRIEKMSNRQVTFSKRRHGLLKKARELAILCDVQVGVIVFSSTGRLYEYASSTMPSIIHKYQSAQEHQQLL-PVSQIM---------------------
-----RLSNLGVNDLCLLENQLEQSLRCMREKK--------------GYIFHQQNIQLSEEIKLIHERNLE-----------------------------
>TaAGL17.11A2_FGENESH
------------------VTFSKRRSGLLKKARELAILCNVQVSAMVFSSTCRLYEYANSTMPSIIENYQSAQEQQQLL-
RVSQVMYWQGGVWKLQKEMQMLQEHHRNLMGERLSNLGVKDLCLIENQLEKSLHHIREKK--------------GYIFHQENIRLSGEIKLIHEQNLE---------
--------------------
>TaAGL17.11A4_FGENESH
MG-GKIVIRRIENMRSR------------------------QVGAMVFSSTCRLYEYANSTMPSIIQNYQSAQEQHQLL-
PVSQVMYWQGEVWKLQQEMQMLQEHHRKLMGERLSKLGAKDLCLMENQLEKRLHRIREKKVAELPQFVLIYNHDGYIFHQENIRLSGEMKLIHEQNLK---------
--------------------

```
>TaSOC1.5B_TraesCS1B01G214600
MVRGKTEMKRIENATSRQVTFSKRRNGLLKKAFELSVLCDVEVALAVFSPRGRLYEFSSTTLQKSIDCYKAYTKDNVNNK------
QVKADTVSLAKKLEALEVSKLKILGENLGGCSAEELNCLEVNIEKSLHIIRGKKTQVLEQQIANLKEKERTLLKDNEDLRGKQRDIEASLVVEPVPSD---------
--ETELYIGLPRCS
>TaSOC1.4B_FGENESH
MARGKTRVKLIEDRTSRQVAFSKRRRGLCKKAFELSVLCDAEVALIVFSPTGRLYKFANAG-----------TPLNHRSQL-------------------
PCRSKNKKNSGNNARGTTYCPLPLPTNGTTRAVPLLRRS----------------------------------------------------------------PRCS
>TaAGL17.7B2_FGENESH
MVRRKTVIERIEDTTSRQVTFSKRKGGLFKKARELGVLCDAQVGILLFSNTVRLYEYSNSSMA-----------------STSATR----
SAAVRAVDLRVLVVVHEKGKGERVARLTASEMSWCGGDSEGDLRRWR-------------------------------FGLLRQHEVMQNGPGQTNLIN--------
----------
>TaBS.7D1_FGENESH
--------KRIDDDGSRQVTFCKRRGTLLKKACELAVLCDVSLGLVVFSSTSKLADYCSTR----------------------------------------------
----------------------------------------------------------------------------------------------------
>TaBS.3D_TraesCS7D01G633200LC
MGRG-----MLDDDVSRRATFGERSGALLTEAQEVAALWEADVGVLIFDSAG------STSWSELMQRYQIITKGK--
QGIDDRRQQLLGEIARLRRERDRLEASVRGLTGDNLPSAT-TELGDLEQQVERALGH------
KLLEQQLDEIHHRVHILEEQNSFLRHMMSEEGRQRAAASALVAELLFGGFFPEESTSLRL-WPGS-
>TaMIKC.2B_TraesCS4B01G023300
MGRVKLAIKRIENNTNRHVTFSKRRNGLIKKAYELSVLCDIDIALLMFSPSRRLCPFSGHGVEDVLLRYLNMSDNDRGEP--
QNREEIQKEIYACQQQLQISEERLRTLFPDPAAFGSMGEIDNCEKSLMDMLTRVVERKNYLLS-NLAPFDPTAPGMQGG------EAQMVH-----DDGQNPGH---
----NTLSTLCLGDDESG
>TaANR1D-likeD_FGENESH
MVRGKIVIRRIENMSRRPVTFSKRRHGLLKKARELAILCDVEVGVIVF-STGHLYEYANSTMPSIIQNYQSAQEQHQLL-
PVSQVMFWQEEVRKLQEEMQMLEEHHRANMGERLSNLAVTNICLMENQLEKSLHRIREKK-----------------------------------------------
--------------------
>OsMADS3_Os01g10504
MGRGKIEIKRIENTTNRQVTFCKRRNGLLKKAYELSVLCDAEVALIVFSSRGRLYEYANNSVKSTVERYKKANSDTSNSGTVNA-
QHYQQESSKLRQQISSLQNANRTIVGDSINTMSLRDLKQVENRLEKGIAKIRARKNELLYAEVEYMQKREVELQNDNMYLRSKVVENEQPLNMAAST-
SEYQQPQHYAHLPTTLQLGSRGVD
>OsMADS32_Os01g52680
MGRGRSEIKRIENPTQRQSTFYKRRDGLFKKARELAVLCDADLLLLLFSASGKLYHFLSPSVREFVERYEATTHTK-----
ADIRQERRAELEKVGSMCDLLEKQLRFMTVDDGEEYTVPSLEALEHNLEAAMRKVRSEKDRKIGGEICYLQNIIRGRQEERYGLCDKIAH--QTLKDCGSTSLSN--
-------GLDLKLGFN---
>OsMADS2_Os01g66030
MGRGKIEIKRIENSTNRQVTFSKRRSGILKKAREISVLCDAEVGVVIFSSAGKLYDYCSTSLSRILEKYQTNSGKI-----
DEKHKSLSAEIDRIKKENDNMQIELRHLKGEDLNSLQPKELIMIEEALDNGIVNVNDKLMDHWERH----VRTDKMLEDENKLLAFKLH---
QQDIAGSMRDLELDFAAQMPI---TFRV-QPQEN
>OsMADS21_Os01g66290
MGRGKIEIKRIENKTSRQVTFCKRRNGLLKKAYELAILCDAEIALIVFSSRGRLYEFSNNSTRSTIERYKKASASTSGSAPVNSHQYFQQEAAKMRHQIQTLQNANR
HLIGESIGNMTAKELKSLENRLEKGISRIRSKKHELLFSEIEYMQKREADLQNENMFLRAKVAEAEHDDQQSSGT--ELAAAAQYSSHQTALHLGY-KVD
>OsMADS51_Os01g69850
ARRGRVQLRRIEDKASRQVRFSKRRAGLFKKAFELALLCDVEVALLVFSPVGKLYEYSSSSIEGTYDRYQQFAGARRDLN---EGSTSINSDENAS---
SRLRDITAWSLQNNADESDANQLEKLEKLLTNALRDTKSKK---ML-------------AKQNGEGSR-------------------------------------S-
S
>OsMADS60_Os02g01360
AG-KKGKKKQAKDELDRQKQAEKKRRRLEKAANSAAIIS-------------------E--LE----KKKQ----KK----RE--E-----
QQRLDEEGAAIAEAVHVLIGEDM--LNKDHSAGFDFAVDAGLIQCADHKGRCIDQPLPSWEVKDLQLQAPYQGMFHQVAC-------PG---------------
VSSLQIG--DIT
>OsMADS29_Os02g07430
MGRGKIEIKRIENATNRQVTFSKRRGGLLKKANELAVLCDARVGVVIFSSTGKMFEYCSCSLRELIEHYQTVTNTH--EEI---
DQQIFVEMTRMRNEMEKLDGGIRRFTGDDLSNLTLADINDLEQQLEFSVTKVRARKHQLLNQQLDNLRRKEHILEDQNSFLCRMINENHHQAAVGDVKAMVEMLT--
--AESTALQL-TPQDP
>OsMADS27_Os02g36924
MGRGKIVIRRIDNSTSRQVTFSKRRNGIFKKAKELAILCDAEVGLMIFSSTGRLYEYSSTSMKSVIDRYGKSKDEQQAVAPNSELKFWQREAASLRQQLHNLQENHR
QLMGEDLSGLNVKELQSLENQLEISLRSVRTKKDHVLIDEIHELNRKGSLVHQENMELYKKISLIRQENAETEGPSEVNPTPYNFAVNPVQLGLSTLHSD
>OsMADS6_Os02g45770
MGRGRVELKRIENKINRQVTFSKRRNGLLKKAYELSVLCDAEVALIIFSSRGKLYEFGS-GITKTLERYQHCCYNAQDSN-
LSETQSWYHEMSKLKAKFEALQRTQRHLLGEDLGPLSVKELQQLEKQLECALSQARQRKTQLMMEQVEELRRKERQLGEINRQLKHKLEVEGTS-NYQGAVVEN-
QPPPHSAA-EPTLQIGY-RST
>OsMADS57_Os02g49840
MGRGKIVIRRIDNSTSRQVTFSKRRNGLLKKAKELSILCDAEVGLVVFSSTGRLYEFSSTNMKTVIDRYTNAKEELLGG-
ATSEIKIWQREAASLRQQLHNLQESHKQLMGEELSGLGVRDLQGLENRLEISLRNIRMRKDNLLKSEIEELHVKGSLIHQENIELSRSLNVMSQQKLEQRGATDANS
TPYSFRINPPSLELSQSEGE
>OsMADS22_Os02g52340
MARERREIKRIESAAARQVTFSKRRRGLFKKAEELSVLCDADVALIVFSSTGKLSHFASSSMNEIIDKYNTHSNNGKAE-
PSLDLNLEHSKYAHLNEQLAEASLRLRQMRGEELEGLSIDELQQLEKNLEAGLHRVMLTKDQQFMEQISELQRKSSQLAEENMQLRNQVSQISPAEK--
GQSSESVLHSGSSQSGDVSLKLGLPAWK
>OsMADS50_Os03g03100
MVRGKTQMKRIENPTSRQVTFSKRRNGLLKKAFELSVLCDAEVALIVFSPRGKLYEFASARIRP-----------EKTAKI-------------------------
---------------------FPRVAIELPSKQSHYFHKEISCETGKVRN--------------------------------------EIGNLVLAF----
>OsMADS47_Os03g08754
GKRERIAIRRIDNLAARQVTFSKRRRGLFKKAEELSILCDAEVGLVVFSATGKLFQFASTSMEQIIDRYNSHSKTQRA-
ESQLDLQGDSSTCARLKEELAETSLRLRQMRGEELHRLNVEQLQELEKSLESGLGSVLKTKSKKILDEIDGLERKRMQLIEENLRLKEQVSRMSRMEE-
EGQSSESVSYPRP--PSDTSLRLGLSSSK
>OsMADS1_Os03g11614
MGRGKVELKRIENKISRQVTFAKRRNGLLKKAYELSLLCDAEVALIIFSGRGRLFEFSSSCMYKTLERYRSCNYNSQDAA-P-
ENEINYQEYLKLKTRVEFLQTTQRNILGEDLGPLSMKELEQLENQIEVSLKQIRSRKNQALLDQLFDLKSKEQQLQDLNKDLRKKLQETSAENVLDGGGHSGSHHQG
LLHH-DHSLQIGYH-DM
>OsMADS14_Os03g54160
MGRGKVQLKRIENKINRQVTFSKRRSGLLKKANEISVLCDAEVALIIFSTKGKLYEYATSCMDKILERYERYSYAEKVLIASDTQGNWCHEYRKLKAKVETIQKCQK
HLMGEDLESLNLKELQQLEQQLENSLKHIRSRKSQLMLESINELQRKEKSLQEENKVLQKELVEKQKV-----------SSSFMMRE-LPTTNIS--VAA
>OsMADS34_Os03g54170
MGRGKVVLQRIENKISRQVTFAKRRNGLLKKAYELSILCDAEVALVLFSHAGRLYQFSSSNMLKTLERYQRYIYASQDAAPSDEMQNNYQEYVNLKAHVEILQQSQR
NLLGEDLAPLATNELEQLESQVVRTLKQIRSRKTQVLLDELCDLKRKEQMLQDANRVLKRKLDEIDVEAAPNCSNGHGGQPEHFFQA-------------
>OsMADS25_Os04g23910
MGRGKIAIKRIDNTMNRQVTFSKRRGGLMKKARELAILCDADVGLIVFSCTGRLYDFSSSSMKSIIERYQEAGEEHCRLLPMSEAKFWQREVTTLRQQVQNLHHNNR
QLLGEEISNFTVRDLQLLQNQVEMSLHSIRNKKDQLLAEEILKLNEKGSLVQKENSELRKKFNIAHQRNIEGESTSSEQKDPGESSTRCIDLELSQKEDE
>OsMADS61_Os04g38770
```

MGRGKIVIRRIDNSTSRQVTFSKRRNGIFKKAKELAILCDAEVGLVIFSSTGRLYEYASTSMKSVIDRYGRAKEEQQHVAPNSELK---------------------
---------------------------------------------------------------------------------------SFIYITEN-----
>OsMADS17_Os04g49150
MGRGRVELKRIENKINRQVTFSKRRNGLLKKAYELSVLCDAEVALIIFSSRGKLYEFGS-GINKTLEKYNSCCYNAQGSN-
LAEHQSWYQEMSRLKTKLECLQRSQRHMLGEDLGPLSIKELQQLEKQLEYSLSQARQRKTQIMMEQVDDLRRKERQLGELNKQLKNKLEAEADSSNCHGTVVSG-
PPP----D-EPTLQIGY-SNG
>OsMADS31_Os04g52410
MGRGRVELKKIENPTNRQVTFSKRRMGLLKKANELAILCDAQIGVIVFSGTGKMYEYSSWRIANIFDRYLKAPSTR-----
MDVQQRIIQEMTRMKDENNRLRIIMRQYMGDDLASLTLQDVSNLEQQIEFSLYKVRLRKQQLLDQQLLEMHSRMQIPGDQSNYLCHMNLIGEQAQAP------LM--
SSQMYNEMTALQL-SPQEE
>OsMADS66_Os05g11380
-GSGSSEGSSIEDTADRQVTFCKRCNGLLKKAYELSMLCDAEVALIVFSSRGRLYEYSNNSVEETIERYKKANSDTSNTSTINA-
QHYQQEAAKLKQHITYLQNSNRFII----------------------------------------------SLSCLDFFFST-------------------------
---------------
>OsMADS58_Os05g11414
GSRGKIEIKRIENTTNRQVTFCKRRSGLLKKAYELSVLCDAEVALVVFSSRGRLYEYSNNSVKETIERYKKANSDTSNASTINA-
QHYQQEAAKLKQQITNLQNSNRTLVGDNITTMNHRELKQLEGRLDKGLGKIRARKNELLCAEIEYMQRRETELQNDNMYLKSKVAESEQTVNMSASTS-
EYHQPQYYPEERKAFMSGKKQCN
>OsMADS4_Os05g34940
MGRGKIEIKRIENSTNRQVTFSKRRAGILKKAREIGVLCDAEVGVVIFSSAGKLSDYCTTTLSRILEKYQTNSGKI-----
DEKHKSLSAEIDRVKKENDNMQIELRHMKGEDLNSLQPKELIAIEEALNNGQANLRDKMMDHWRMH----KRNEKMLEDEHKMLAFRVH---
QQEVEGGIRELELDFAASMPF---TFRV-QPQQE
>OsMADS5_Os06g06750
MGRGKVELKRIENKISRQVTFAKRRNGLLKKAYELSVLCDAEVALIIFSTRGRLFEFSTSCMYKTLERYRSCNYNSCEAS-
ALETESNYQEYLKLKTRVEFLQTTQRNLLGEDLVPLSLKELEQLENQIEISLMNIRSSKNQQLLDQVFELKRKEQQLQDANKDLKRKIQETSGENMLDVGP---S-
QEFLH-H-DPSLHIGY----
>OsMADS55_Os06g11330
MARERREIRRIESAAARQVTFSKRRRGLFKKAEELAVLCDADVALVVFSSTGKLSQFASSNMNEIIDKYTTHSKNGKTDKPSIDLN---------------------
-MRGEELEGLSVEELQQMEKNLEAGLQRVLCTKDQQFMQEISELQRKGIQLAEENMRLRDQMPQVPTAG--DGQSSESVLNSGS--SGDISLKLG-----
>OsMADS63_Os06g11970
MGRVKLQIKRIENIPNRQVTFSKRRNGLIKKAYELSVLCDIDIALLMFSPSGRLSHFSGRRIEDVLTRYINLPESDRGGT--
QNREELQQEIRRCQHQMQLTEEQLRMFEPDPARSASMEDVEASEKFIAGILSRVEERKRYLLC-SMGSFDVTASAMQHL------LPQQQH-----
EGMPPTTSTVVPEMGMQLNALTMGMLGNG
>OsMADS59_Os06g23950
MVRGKTVISRIENTTSRQVTFSKRRSGLFKKAKELAILCDAQVGVLVFSSTGRLYDYSNSSNSRNLT--------------------------------------
YDQLLG------------------------------------------------------------------------------------------------
>OsMADS30_Os06g45650
MGQGKIEMKRIEDATRRQVTFSKRRAGFLKKANELAVLCDAQVGVVVFSDKGKLFDFCSVILMELFHRYEITTRNT--QET---
DEQMVMEITRLRNEIDQLEASLRRQTGEDLSSSTVDELSQLQLQLESSLSKVHARKDELMSQQLEDMRRMHQTVHEQNNFLCRMVTKI------
LNVITIMIMQLGYIVIN--SWRFW-----
>OsMADS16_Os06g49840
MGRGKIEIKRIENATNRQVTYSKRRTGIMKKARELTVLCDAQVAIIMFSSTGKYHEFCSTDIKGIFDRYQQAIGTS-----
IEQYENMQRTLSHLKDINRNLRTEIRQRMGEDLDGLEFDELRGLEQNVDAALKEVRHRKYHVITTQTETYKKKVKHSYEAYETLQQELGLREEPAFGDNT-------
------AMFAFRV-VPHGM
>OsMADS15_Os07g01820
MGRGKVQLKRIENKINRQVTFSKRRNGLLKKAHEISVLCDAEVAAIVFSPKGKLYEYATSRMDKILERYERYSYAEKALIASESEGNWCHEYRKLKAKIETIQKCHK
HLMGEDLESLNLKELQQLEQQLESSLKHIISRKSHLMLESISELQKKERSLQEENKALQKELVERQKNVRGQTQVQAQASSS-MLRD-LPPQNIC--AAV
>OsMADS18_Os07g41370
MGRGPVQLRRIENKINRQVTFSKRRNGLLKKAHEISVLCDADVALIVFSTKGKLYEFSSSSMEGILERYQRYSFDERAVLPTEDQENWGDEYGILKSKLDALQKSQR
QLLGEQLDTLTIKELQQLEHQLEYSLKHIRSKKNQLLFESISELQKKEKSLKNQNNVLQK-LMETEKEKNN--------SSPTPVTA-IPTTNNS--AQP
>OsMADS26_Os08g02070
MARGKVQLRRIENPVHRQVTFCKRRAGLLKKARELSILCEADIGIIIFSAHGKLYDLATGTMEELIERYKSASGE--
QANAGDQRMDPKQEAMVLKQEINLLQKGLRYIYGNRAEHMTVEELNALERYLEIWMYNIRSAKMQIMIQEIQALKSKEGMLKAANEILQEKIVEQNDVGMM--DQ--
QNTNPLTILSYCRGSEMGYS---
>OsMADS23_Os08g33488
MGRGKIEIKRIDNATSRQVTFSKRRSGLFKKARELSILCDAEVGLLVFSSTSRLYDFASSSMKSIIERYNETKEDPHQTMASSEAKLWQQEAASLRQQLHNLQEYHR
QLLGQQLSGLDVEDLQNLESKLEMSLKNIRLRKDNVMMDQIQELSRK------------------------------VV---------------------
>OsMADS62_Os08g38590
MGRVKLPIKRIENTTNRQVTFSKRRNGLIKKAYELSVLCDIDVALLMFSPSGRLSHFSGRGVEDVILRYMNLSEHDRGEA--
QNREEIQQEIYSSQQQLQITEDRLRMFEPDPAAFGTSSEVDGCEKYLMELLTRVVERKNNLLSSHMAPFDATTAAMQGA------GTQMV------
DGGADPGHAAAVGCVDALSTLCLGEDNSG
>OsMADS7_Os08g41950
MGRGRVELKRIENKINRQVTFAKRRNGLLKKAYELSVLCDAEVALIIFSNRGKLYEFCSQSMTKTLEKYQKCSYAGPETAQSEQLKASRNEYLKLKARVENLQRTQR
NLLGEDLDSLGIKELESLEKQLDSSLKHVRTTRTKHLVDQLTELQRKEQMVSEANRCLRRKLEESNHVRG-QGCNLIGYGGNGFFHPAEPTLQIGY----
>OsMADS37_Os08g41960
RKRGKVELRRIEDRTSRQVRFSKRRSGLFKKAYELSVLCDAQVALLVFSPAGRLYEFASSSIDTIFGRYWDLLDTTIDLN---IEARESRVD---
CDPVPKINHITQCVLESNVNELNIAELRGLEEAMTNALTVVKNKL---MMKVASVLPQSEK----------------------------------------------
--C-S
>OsMADS8_Os09g32948
MGRGRVELKRIENKINRQVTFAKRRNGLLKKAYELSVLCDAEVALIIFSNRGKLYEFCSQSMTRTLERYQKFSYGGPDTAQNELVQSSRNEYLKLKARVENLQRTQR
NLLGEDLGTLGIKELEQLEKQLDSSLRHIRSTRTQHMLDQLTDLQRREQMLCEANKCLRRKLEESNQLHG-HGATLLGYGGNGFFHSAEPTLQIGF----
>OsMADS56_Os10g39130
MVRGRTELKRIENPTSRQVTFSKRRNGLLKKAFELSVLCDAEVALIVFSPRGRLYEFASAPLQKTIDRYKAYTKDHVNNKIQQDIQQVKDDTLGLAKKLEALDESRR
KILGENLEGFSIEELRGLEMKLEKSLHKIRLKKTELLEQQIAKLKEKERTLLKDNENLRGKHRNLEAAALVTAGAADA---------DETDLYIGLPERS
>OsMADS68_Os11g43740
MGRVKLKIKKLENSSGRHVTYSKRRSGILKKAKELSILCDIPLILLMFSPNDKPTICVGSSIEDVITKYAQQTPQERA---
LESLEELSSHLGALQCQMADVEKRLS-YWSDPEKVENIDHIRAMEQSLKESLNRIRIHKENFAKQHLMSLQCAAAQFQND------KLPL-------
GGGGAEAQGEQH--HH-LTSLQLGQF---
>OsMADS33_Os12g10520
MVRGKVQMRRIENPVHRQVTFCKRRGGLLKKARELSVLCDADVGVIIFSSQGKLHELATGNMHNLVERYQSNVAGGQMEPGLQRQQVAEQGIFLLREEIDLLQRGLR
STYGGGAGEMTLDKLHALEKGLELWIYQIRTTKMQMMQQEIQFLRNKEGILKEANEMLQEKVKEQQMSLLD--SQ--QPTQPMTYGNF-------FS---
>OsMADS13_Os12g10540
MGRGRIEIKRIENTTSRQVTFCKRRNGLLKKAYELSVLCDAEVALIVFSSRGRLYEYSNNNVKATIDRYKKAHACGSTSGPVNAQQYYQQESAKLRHQIQMLQNTNK
HLVGDNVSNLSLKELKQLESRLEKGISKIRARKNELLASEINYMAKREIELQNDNMDLRTKIAEEEQQVTVSAAM--ELVQAVA-AQQPTELNLGYHHLA
>OsMADS20_Os12g31748
MGRGKVQVRRIENEVSRQVTFSKRRPGLLKKAHEIAVLCDVDVAAIVFSAKGNLFHYASTTMERILEKYDRHELLSENVIEPELEGSMSYDHIKLRGRIEALKKSQR
NLMGQELDSLTLQDIQQLENQIDTSLNNIRSRKNNLLLKSIAELRQKEKLLMEKNTILEKKITELETLHTC--------APPACNTA-VPNLNIC--TAP
>TaAGL17.9A_TraesCS1A01G044900

```
MVRGKAVIERIENTTSRQVTFSKRKSGLFKKARELGVLCDAQVAVLLFSNTGRLYDYSNSNMKSIIERYQHVKEGQQFM-
ASAEAKFWQAEGVRLRQQLHNLQENHRQLLGQHLSGLGLEDLRGLENQLETSIHNIRLTKDQLMIDETEELKKKASLVHQENIELHKKLNIIRQENIDQAEVNGTIS
SEYYIAA-PVRLELSYPAER
>TaAGL17.8A_TraesCS1A01G045000
MVRGKTVIEKIENTTSRQVTFSKRKSGLFKKARELGVLCDAQVAVLLFSNTGRLYDYSSSNMKSLLERYQHVKEGQQLM-
ASTEAKFWQAEGERLRQQLHNLQENHRQLLGQHLCGLGLEDLSGLEKQLETSLHNIRLAKDQLMIDEIEEFNKKGHLVHQENVELHKKLNIIHQENIYKPEANGDY-
--------------------
>TaAG.1A_TraesCS1A01G125800
MGRGRIEIKRIENTTNRQVTFCKRRNGLLKKAYELSVLCDAEVALIVFSGRGRLYEYSNNSVKATIERYKKATSDTSSAGTINA-
QHYQQESAKLKQQITTLQNSNRTLIGDTMATMSHRDLKQLEGRLDKGLGKIRARKNELLCAEIEYMQRREMELQNNNFFLREKVAETEQTLNMAASTSNEYQHPQYC
SQERKSFNSVGR---
>TaSOC1.1A_TraesCS1A01G199600
MVRGKTQMKRIENATSRQVTFSKRRNGLLKKAFELSVLCDAEVALVVFSPRGRLYEFASATLQKSIDRYKAYTKDTVNNKVQPDIQQVKADALSLAKKLEALEDTKR
KILGENLGGCSTEELHFLEGKIEKSLRVIRGKKTQLLEQQIANLKEKERTLLKDNEDLRGK-RNLEAPLFLQPVPRD----------DETELYIGLPRCS
>TaSOC1.5A_TraesCS1A01G199900
MVRGKTEMKRIENETSRQVTFSKRRNGLLKKAFELSVLCDVEVALAVFCPRGRLYEFSSATLQKSIDRYKSYRKDNVNNKVQPDIQQVKADTVSLAKKLEALEVSKL
KILGENLGGCSAEELNCLEVNIEKSLRIIRGKKTQVLEQQIANLKEKERTLLKDNEDLRGKQRDLEASLVVEPVPSD----------DETELYIGLPRCS
>TaSTK.2A_TraesCS1A01G262700
MGRGKIEIKRIENTTSRQVTFCKRRNGLLKKAYELSVLCEAEIALIVFSARGRLYEYASNSTRTTIDRYKKASASASGSAPVNSQQYFQQESAKLRHQIQSLQNANR
NLMGESVGNLTLKELKSLENRLDKGIGRIRAKKHELLFAEIEYMQKLEADLQSENMYLRAKVADAELAAPPSGGA--ELSSSSRYSQSTTALHLGYQ---
>TaPI.1A_TraesCS1A01G264300
MGRGKIEIKRIENSSNRQVTFAKRRAGLVKKAREIGVLCDAEVGVVIFSSAGKLYDFWTTTLPRILEKYQTNSGKI-----
DEKHKSISAEIDRVKKENDNMQIELRHMKGEDVNSLQPKELIAIEEALTNGQTNLRDKMMDHWKMH----RRNEKMLEEEHKLLALRMH---QQD-
DSGMREMELDFTSQMPF---TFRL-QPQED
>TaBS.5A5_TraesCS1A01G491400LC
MAQG------------AQPHHGHRRGA------------------------------QLE---------------------
DQRWKLLADIATLRHERDHLEASVRRQTVEDLPSATATELRGLEHKLECALGKVRETKDKLMEEQLDESHHRVHILEEQNNFLGHMILKVLKHNLCPAKC-------
------------------
>TaAGL17.9B_TraesCS1B01G058100
MVRGKTVIKRIENTTSRQVTFSKRKSGLFKKARELGVLCDAQVGVLLFSNTGRLYDYSNSNMKSIIERYQHVKEGQQFM-
ASAEAKFWQAEGERLRQQLQNLQENHRQLLGQHLSGLGLEDLRGLENQLETSINNIRLTKDQLMIDEIEDLNKKESLVHQENIELHKKLNVIRQENIYEAEVNGTIS
SQYSTAA-PVRLELGHPAQM
>TaAGL17.7B1_TraesCS1B01G058200
MVRRKTVIERIEDTTSRQVTFSKRKGGLFKKARELGVLCDAQVGILLFSNTVRLYEYSNSSVNSIIERYQKVKEGQQFM-
PSAEAKFWQAEAERLRQQLHNLQEDHRQLLGQNLPGLGSEGLKDLENQLETSIHNIRLTKDQLMIDEIEELNKKESLVHQENMELHKKLNIIRRQNLDQVELNETIS
CQYNIAAAPTCRKG------
>TaAGL17.8B_TraesCS1B01G058300
MVRGKTVIEKIKNTTSRQVTFSKRKGGLFKKARELGVLCDAQVGVLLFSNTGRLYDYSNSNMKSLLERYQQVKEGQQFM-
ASAQAKFWQAEGERLRQQLHNLQENNRQLLGQHLSGLGLEDLSGLEKQLETSLHNIRLAKDQLTIDEIEEFNKKGNLVHQENVELHKELNIIHQENIYQPEANGAIS
SQCSIAAALVRLELSKPAEK
>TaAG.1D_TraesCS1B01G144800
MGRGRIEIKRIENTTNRQVTFCKRRNGLLKKAYELSVLCDAEVALIVFSGRGRLYEYSNNSVKATIERYKKATSDTSSAGTINA-
QHYQQESAKLKQQITTLQNSNRTLIGDTMATMSHRDLKQLEGRLDKGLGKIRARKNELLCAEIEYMQRREMELQNNNFFLREKVAETEQTLNMAASTSNEYQQPQYY
SQERKSLNSVGR---
>TaSOC1.1B_TraesCS1B01G214500
MVRGKTEMKRIENATSRQVTFSKRRNGLLKKAFELSVLCDAEVALVVFSPRGRLYEFASATMQKSIDRYKAYTKDTVNNK------
QVKADALSLAKKLEALEDTKRKILGENLGGCSTEELHFLEGKIEKSLRVIRGKKTQLLEQQIAKLKEKERTLLKDNEDLRGKQRNLEAPLLLEPVPRD---------
-DETELYIGLPRCS
>TaSTK.2B_TraesCS1B01G273300
MGRGKIEIKRIENTTSRQVTFCKRRNGLLKKAYELSVLCEAEIALIVFSARGRLYEYASNSTRTTIDRYKKASASASGSAPVNSQQYFQQESAKLRHQIQSLQNANR
NLMGESVGNLTLKELKSLENRLDKGIGRIRAKKHELLFAEIEYMQKLEVDLQSENMYLRAKVADAELAAPPPGGA--ELSSSSRYSQSTTALHLGYQ---
>TaPI.1B_TraesCS1B01G275000
MGRGKIEIKRIENSSNRQVTFAKRRAGLVKKAREIGVLCDAEVGVVIFSSAGKLYDFWTTTLPRILEKYQTNSGKI-----
DEKHKSISAEIDRVKKENDNMQIELRHMKGEDVNSLQPKELIAIEEALTNGQTSLRDKMMDHWKMH----RRNEKMLEEEHKLLALRMH---QQD-
DSNMREMELDFTSQMPF---TFRL-QPQED
>TaBS.7B_TraesCS1B01G467500
MGRGKLEVKRIDDDGSRLVTFCKRRGTLLKKARELAVLCDASLGLVVFSSTGNLADYCSTRY-----------------------------VARLICPVLSL-----
----------------------------------------------MHARAPV-------------------------IFPLSFVK-------------
>TaAGL17.9D_TraesCS1D01G045500
MVRGKTVIERIENTTSRQVTFSKRKSGLFKKARELGVLCDAQVAVVLFSNTGRLYDYSNCNMKSIIERYQHVKEGQQFM-
ASAEAKFWQAEGERLRQQLHNLQENHRQLLGQHLSGLGLEDMRGLESQLETSIHNIRLTKDQLMIDEIEELNKKESLVHQENIELHKKLNIIRQENIYEAEVNGTIS
SQYSTAA-PVRLELSHPAER
>TaAGL17.7D_TraesCS1D01G045600
MVRRKTVIERIEDTTSRQVTFSKRKGGLFKKARELGVLCDAQVGILLFSNTGRLYEYSNSTVNSIIERYQKVKEGQQFM-
ASAEAKLWKVEANRLRQQLHNLQEDHRQLLGQNLPGLGLEGLKDLENQLETSIHNIRLTKDQLMIDEIEELNKKESLVHQENTELYKKLNIIRRQNLDQVEVNETIS
SQYNIAAAPVCLQVSHPAER
>TaAGL17.8D_TraesCS1D01G045700
MVRGKTVIEKIENTTSRQVTFSKRKGGLFKKARELGVLCDAQVGVLLFSNTGRLYDYSNSNMKSLLERYQQVKEGQQFM-
ASAEAKFWQAEGERLRQQLHNLQENNRQLLGQHLSGLGLEDLSGLEKQLETSLHNIRLAKDQLMIDEIEEFNKKGNLVHQENVELHKKLNIIHQENIYQPEANGVIS
SHCSIAAALVRLELSQPAER
>TaAG.1B_TraesCS1D01G127700
MGRGRIEIKRIENTTNRQVTFCKRRNGLLKKAYELSVLCDAEVALIVFSGRGRLYEYSNNSVKATIERYKKATSDTSSAGTINA-
QHYQQESAKLKQQITTLQNSNRTLIGDTMATMSHRDLKQLEGRLDKGLGKIRARKNELLCAEIEYMQRREMELQNNNFFLREKVAETEQTLNMAASTSNEYQQPQYY
SQERKSFNSVGR---
>TaSOC1.1D_TraesCS1D01G203300
MVRGKTQMKRIENATSRQVTFSKRRNGLLKKAFELSVLCDAEVALVVFSPRGRLYEFASATLQKSIDRYKAYTKDTVNNKVQPDIQQVKADALSLAKKLEALEDSKR
KILGENLGGCSTEELHFLEGKIEKSLRVIRGKKTQLLEQQIAKLKEKERTLLKDNEDLRGK-RNLEARLLLEPVQRD----------DETELYIGLPRCS
>TaSOC1.5D_TraesCS1D01G203400
MVRGKTEMKRIENATSRQVTFSKRRNGLLKKAFELSVLCDVEVALAVFSPRGRLYEFSSATLQKSIDRYKAYTKDNVNNKVQPDIQQVKADTVSLAKKLEALEVSKL
KILGENLGGCSAEELNCLEVNIEKSLRTIRGKKTQVLEQQIANLKEKERMLLKDNEDLRGKQRDLEASLVVEPVPSD----------DETELYIGLPRCS
>TaSTK.2D_TraesCS1D01G262700
MGRGKIEIKRIENTTSRQVTFCKRRNGLLKKAYELSVLCEAEIALIVFSARGRLYEYASNSTRTTIDRYKKASASASGSAPVNSQQYFQQESAKLRHQIQSLQNANR
NLMGESVGNLTLKELKSLENRLDKGIGRIRAKKHELLFAEIEYMQKLEADLQSENMYLRAKVAEAELAAPPSGGA--ELSSSSRYSQSTTALHLGYQ---
>TaPI.1D_TraesCS1D01G264500
MGRGKIEIKRIENSSNRQVTFAKRRAGLVKKAREIGVLCDAEVGVVIFSSAGKLYDFWTTTLPRILEKYQTNSGKI-----
DEKHKSISAEIDRVKKENDNMQIELRHMKGEDVNSLQPKELIAIEEALTNGQTNLRDKMMDHWKMH----RRNEKMLEEEHKLLALRMTCISKWSIICTM--
VAYTKSNLRPP---AGRP-EQ---
>TaBS.7D2_TraesCS1D01G441200
```

MGRGKLEVKRIVDDASRRVTFCKRRGTLLKKARELAVLCDVSLGLVVFSSTGKLADYCSTSWSDLIQRYERESTPS-
VQDTGDHHQQLLAEVARLRQEIDHLEGSLRRQTGEELSSVTAAELHDLQHHVGSALGKVRHRK--------------VHLLEE------------------------
--LATAAFLEESTSLQP------
>TaBS.5A3_TraesCS2A01G153100
-------------------------------------------------------------------------------------
MKLLADIATLRHERDHLEASVRRQIVEDLPSTTAAELRGLEHKLECALGKVRETKDKLMEEQLDESHHRVHILEEQNSFLGHMILKVLKHNLCPTKC----------
---------------
>TaAP1.2A_TraesCS2A01G174300
MGRGPVQLRRIENKINRQVTFSKRRSGLLKKAHEISVLCDAEVALIVFSTKGKLYEYSSSSMDVILERYQRYSFEERAVLPIGNQANWGDEYGSLKIKLDALQKSQR
QLLGEQLDPLTTKELQQLEQQLDSSLKHIRSRKNQLLFESISELQKKEKSLKDQNGVLQKHLVETEKEKNN--ATNIHHPSPTPATA-MAPPNIGP-PQP
>TaAP1.3A_TraesCS2A01G261200
MGRGKVQLKRIENKINRQVTFSKRRNGLLKKAHEISVLCDAEVAVIVFSPKGKLYEYATSSMDKILERYERYSYAEKALIASESEGNWCHEYRKLKAKIETIQKCHK
HLMGEDLDSLNLKELQQLEQQLESSLKHIRSRKSHLMMESISELQKKERSLQEENKALQKELVERQKAAASQTQTHAHTSSSFMMRD-APQQNIC--AAA
>TaAGL12.2A_TraesCS2A01G311100
MARGKVQMRRIENPVHRQVTFCKRRMGLLKKAKELSVLCDADIGVMVFSPHGKVYELATGNMQGLIERYKGSNTEAHGESSQNKPEVIQQEVLLLRQEIDLLQKGLR
YMYGENDNHMNLNELQALESNLEIWVHNIRYTKMQIISREIEMLKTKEGILKAANDILQERIIEQSDTGSN-------------MMPFQRTMEGGYY---
>TaAGL17.1A_TraesCS2A01G337900
MGRGKIVIRRIDNSTSRQVTFSKRRNGIFKKAKELGILCDAEVGLVIFSSTGRLYEYASSSMKSVIDRYGRAKEEQQLVAPNSELKFWQREAASLRQQLHNLQENHR
QLMGQDLSGMGVKELQALENQLEISLRCIRTKKDQILIDEIHELNHKGSLVHQENMELYKKINLIRQENVETEAVTEVNRTPYNFAVNSVDLELNSPQND
>TaBS.2A_TraesCS2A01G422400
MGRGKVELKKIENTTSRQVTFSKRRMGLLKKANELAILCDAQVGVIVFSGSGKMYEYASWRIANIFDRYLKAPSTR-----
MDVQQKIIHEMTRMKDESNRLKIIMRQYMGEDLGSLTLQDVSNLEQQIEFSLYKVRLRKQQLLDQQLLEMRQRMHMSEDQSSYMFHMNPARDQPGQS-----ADV--
-DQIYGEMTALKL-SPQE-
>TaBS.5A2_TraesCS2A01G475000
-------------------------------------------------------------------------------------
MKLLADITMLRHERDHLEASVRRQTGEDLPSATAVELRGLEHKLECVLGKVRETKDKLMEEQLDESHHRVHILEEQNSFLGHMILKVLKHNLCPAKC----------
---------------
>TaBS.6A2_TraesCS2A01G733800LC
-----IELLGRSNNVSRQSIFGKSRAGLLKKAQELAMLCEADLGVLIFGSADKQMDYCSTMSNSFPS--------T--------------------------
SSRLQTYFGYSLARCPSADLTCI--LLDCSIFSTR-----------------VHVCGELIDSFRKMNLLK-------------------------SLEI-LT---
>TaAP1.2B_TraesCS2B01G200800
MGRGPVQLRRIENKINRQVTFSKRRSGLLKKAHEISVLCDAEVALIVFSTKGKLYEYSSSSMDVILERYQRYSFEERAVLPIGNQANWGDEYGSLKIKLDALQKSQR
QLLGEQLDPLTTKELQQLEQQLDSSLKHIRSRKNQLLFESISELQKKEKSLKDQNGVLQKHLVETEKEKNN--ATNINHPSPTPATA-MATPNIGP-PQP
>TaAP1.3B_TraesCS2B01G281000
MGRGKVQLKRIENKINRQVTFSKRRNGLLKKAHEISVLCDAEVAVIVFSPKGKLYEYATSSMDKILERYERYSYAEKALIASESEGNWCHEYRKLKAKIETIQKCHK
HLMGEDLDSLNLKELQQLEQQLESSLKHIRSRKSHLMMESISELQKKERSLQEENKALQKELVERQKAAASQT-THTHTSSSFMMRD-APQQNVCS-AAA
>TaAGL12.2B_TraesCS2B01G328000
MARGKVQMRRIENPVHRQVTFCKRRMGLLKKAKELSVLCDADIGVMVFSPHGKVYELATGNMQGLIERYKGSNTEAHGESSQNKPEVIQQEVLLLRQEIDLLQKGLR
YMYGENDNHMNLNELQTLESNLEIWVHNIRYTKMQIISREIEMLKTKEGILKAANDILQERIIEQSDTGSN-------------MMPFQRTMESDYY---
>TaAGL17.1B_TraesCS2B01G344000
MGRGKIVIRRIDNSTSRQVTFSKRRNGIFKKAKELGILCDAEVGLVIFSSTGRLYEYASSSMKSVIDRYGRAKEEQQLVAPNSELKFWQREAASLRQQLHNLQENHR
QLTGQDLSGMGVKELQTLENQLEISLRCIRTKKDQILIDEIHELNHKGSLVHQENMELYRKINLIRQENAETEAATEVNRTPYNFAVNSVDLELNSPQND
>TaBS.2B_TraesCS2B01G440900
MGRGKVELKKIENTTSRQVTFSKRRMGLLKKANELAILCDAQVGVIVFSGSGKMYEYSSWRIANIFDRYLKAPSTR-----
MDVQQKIIHEMTRMKDESNRLKIIMRQYMGEDLGSLTLQDVSNLEQQIEFSLYKVRLRKQQLLDQQLLEMRQRMHMSEDQSSYMFHMNLARDQPGQS-----ADV--
-DQIYGEMTALKL-SPQE-
>TaBS.6B2_TraesCS2B01G665600LC
-----------SNNGSRQSTFGKSRAGLLKMA-ELAMLCEADLGILIFDNVGKQMDYCSTMSNSSPLIFSTLSHGT--------------------------
SSQLQTYFGYSLARCPSAALTSI--LLDWSVFSTR-----------------VHVCGEL----------------------------------------------
>TaAP1.2D_TraesCS2D01G181400
MGRGPVQLRRIENKINRQVTFSKRRSGLLKKAHEISVLCDAEVALIVFSTKGKLYEYSSSSMDVILERYQRYSFEERAVLPIGNQANWGDEYGSLKIKLDALQKSQR
QLLGEQLDPLTTKELQQLEQQLDSSLKHIRSRKNQLLFESISELQKKEKSLKDQNGVLQKHLVETEKEKNN--ATNIHHPSPTPATA-MATPNIGP-PQP
>TaAP1.3D_TraesCS2D01G262700
MGRGKVQLKRIENKINRQVTFSKRRNGLLKKAHEISVLCDAEVAVIVFSPKGKLYEYATSSMDKILERYERYSYAEKALIASESEGNWCHEYRKLKAKIETIQKCHK
HLMGEDLDSLNLKELQQLEQQLESSLKHIRSRKSHLMMESISELQKKERSLQEENKALQKELVERQKAAASQ----THTSSSFMMRD-APQQNIC--AAA
>TaAGL12.2D_TraesCS2D01G309300
MARGKVQMRRIENPVHRQVTFCKRRMGLLKKAKELSVLCDADIGVMVFSPHGKVYELATGNMQGLIERYKGSNTEAHGESSQNKPEVIQQEVLLLRQEIDLLQKGLR
YMYGENDNHMNLNELQALESNLEIWVHNIRYTKMQIISREIEMLKTKEGILKAANDILQERIIEQSDTGSN-------------MMPFQRTMEGGYY---
>TaAGL17.1D_TraesCS2D01G325000
MGRGKIVIRRIDNSTSRQVTFSKRRNGIFKKAKELGILCDAEVGLVIFSSTGRLYEYASSSMKSVIDRYGRAKEEQQLVAPNSELKFWQREAASLRQQLHNLQENHR
QLMGQDLSGMGVKELQTLENQLEISLRCIRTKKDQILIDEIHELNHKGSLVHQENMELYKKINLIRQENAETEAATEVNRTPYNFAVNSVNLELNSPQND
>TaBS.2D_TraesCS2D01G418800
MGRGKVELKKIENTTSRQVTFSKRRMGLLKKANELAILCDAQVGVIVFSGSGKMYEYASWRIANIFDRYLKAPSTR-----
MDVQQKIIHEMTRMKDESNRLKIIMRQYMGEDLGSLTLQDVSNLEQQIEFSLYKVRLRKQQLLDQQLLEMRQRMHMSEDQSSYMFHMNPARDQPGQP-----ADV--
-DQIYGEMTALKL-SPQE-
>TaBS.8D1_TraesCS2D01G714300LC
---------MVDDDASRRATYGERNGGLLREAHELAVLCEAEVGILVFDSAGRQLDYCSTRSV-------------------------------------------
RSVVDLLLP-ISVHSGCTLTLITDL-----------------------------------------------------------------------RP---
>TaBS.4A1_TraesCS3A01G090300
MGRGRTEMKRIHNDVSRRATFGKRRRGLLKKAGELAVLCGVDLGLLVFDDAGKLFDYCSTSWSELIERYESITNHKQFQGIDD-DQQQLAD---
LRRERDRLEASVRRQTGEDLPFATAAELDDLEQRLECVLGEVREMKDKLLEQQLDESYHKVRILQDQNSFLRRLMGQEGQQCAAASAV----ALGGSFP-
ELTALRL-WP---
>TaBS.4A2_TraesCS3A01G090900
MGRGRTEMKRIHNDVSRRATFGKRRRGLLKKAGELAVLCGVDLGLLVFDDAGKLFDYCSTSWSELIERYESITNHKQFQGIDD-DQQPLAD---
LRRERDRLEASVRRQTGEDLPFATAAELDDLEQRLECVLGEVREMKDKLLEQQLAESYHKVHILEDQNSYLRRMMGEEGQQRAAASSV----
AFGGSFPEELTTLRL-WP---
>TaMADS32A_TraesCS3A01G284400
MGRGRSEIKRIDNPTQRQSTFYKRRDGLFKKARELAVLCDADLLLLLFSASGKLYQYLAPSVKEFVERYEASTHTK-----
SDIRQERRRELEKVAKMCDLLEKELRFMTVDDGERYTVPSLAALEHNLEAAMHKVRSEKDRKIGGEMSYLENMIRGKQAERYGLCDKLAH--QSLKVGGSTSLNN--
-------GLDLKLGFSVTV
>TaAG.2A_TraesCS3A01G314300
MGRGRIEIKRIENTTNRQVTFCKRRNGLLKKAYELSVLCDAEVALIVFSSRGRLYEYSNNSVKATIERYKKANSDTSNSGTVNA-
QYYQQESSKLRQQISSLQNSNRSLVRDSVSTMTLRDLKQLEGRLEKGIAKIRARKNELMYAEVEYMQKREMELQNDNIYLRSKVSENEQPVNMSGSASSEYQQQQHY
SQLPTALQLGQQN--
>TaPI.2A_TraesCS3A01G406500
MGRGKIEIKRIENQSNRQVTFSKRKNGILKKAKEISVLCDAEVGVVVFSSAGKLYDFCSTSLSRILEKYQTNSGKI-----
DEKHKSLSAEIDRIKKENDNMQIELRHLKGEDLNSLQPKELIMIEEALDNGLTSLHEKQMEHYDRL----MKHGKMLEDENKLLAFKVH-----
DIAGSMRDLELDFAAQMPI---TFRV-QPQED
>TaFLC.5A_TraesCS3A01G432900

```
ARRGRVELRRIEDRTSRQVRFSKRRAGLFKKAFELSLLCDAEVALLVFXXXXXXXXXXXSSIEGTYDRYQQFAVPGRNLI---QEDATVSNDEDPS---
SRLSEIAAWSLDNNADNSDASSLEKLEKLLKDALRITESKK---TL-------------VKQNSGGST-------------------------------------S-
Y
>TaFLC.3A_TraesCS3A01G434400
ARRGRVELRRIEDRTSRQVRFSKRRAGLFKKAFELVVLCDAEVALLVFSPAGKLYEYASSSIEGTYDRYQRFAGAGTNVN---GGDASSNNDGDPS---
STLKEIASWSIQNNADVSDANKLEKLEKLLTDALRNTKSKK---ML-------------VQQNSGASTR------------------------------------S-
N
>TaFLC.4A2_TraesCS3A01G434700
ARRGRVELRRIKDRTSRQVRFSKRRSGLFKKAFELALLCDTEVSLLVFSPAGRFYEYASSR---------------------------------PA-----------
----------------------------------------------------------------------------------------------------
>TaFLC.6A_TraesCS3A01G434900
ARRGRVELRRIEDRTNRQVCFSKRRSGLFKKAFELSLLCDADVALLVFSPAGKLYEYSSSSIEATYDRYQRFAGPGMNVN---GGDARNKNDGDPS---
SRLEDIASWSLQNNADDSDATKLEKLEKLLTDALRNTKSKK---ML-------------AQLNSGASTS------------------------------------S-
N
>TaFLC.4A1_TraesCS3A01G435000
ARRGRVELRRIEDRTSRQVRFSKRRAGLFKKAFELAVLCDAEVSLLVFSPAGRLYEYASSSIEGTYDRYQAFARAGKDVN---EPGASNNNDGDPS---
SRLKEITSWSLQNNADDSDANELEKLEKLLTDALKNTRSKK---ML-------------AQRNSGAGTS------------------------------------S-
G
>TaBS.4B_TraesCS3B01G105700
MGRGRTEMKRIHNDVSRRATFGKRRRGLLKKARELAVLCGVDLGLLVFDDAGKLFDYCSTSWSELIERYESITNHK--QGID--
DQQPLADIAGLRREHDRLEVSVRRQTGEDLPSATAAELDDLEQRLERVLGEVREMKDKPLEQQLDESYHKVHTLEDQNSFLRRLMGEEGQQRAAASAV----
AFGGSFPEELTALRL-WP---
>TaAG.2B_TraesCS3B01G157500
MGRGRIEIKRIENTTNRQVTFCKRRNGLLKKAYELSVLCDAEVALIVFSSRGRLYEYSNNSVKATIERYKKANSDTSNSGTVNA-
QYYQQESSKLRQQISSLQNSNRSLVRDSVSTMTLRDLKQLEGRLEKGIAKIRARKNELMYAEVEYMQKREMELHNDNIYLRSKVSENEQPMNMSGSTSSEYQQQQHY
SQLPTALQLGQQN--
>TaBS.6B1_TraesCS3B01G236300LC
-----------SNNVSRQSTFGKSRAGLLKKV-QLAMLCEADLGVLIFDSAGKQMDYCSTMSNSSLSIFSTLSHGM-------------------------
SSRLQTYFGYSLARCPSVALTCI--PFDCSVFSTR-----------------VHVCDELIDSFRKMNLLK-------------------------SLKI-LT---
>TaMADS32B_TraesCS3B01G318300
MGRGRSEIKRIDNPTQRQSTFYKRRDGLFKKARELAVLCDADLLLLLFSASGKLYQYLAPSVKEFVERYEAATHTK-----
SDIRQERRAELEKVAKMCDLLEKELRFMTVDDGEQYSVPSLAALEHNLEAAMHKVRSEKDRKIGGEMSYLENMIRGKQAERYGLCDKLAH--QSLKVGGSTSLNN--
-------GLDLKLGFN---
>TaPI.2B_TraesCS3B01G440200
MGRGKIEIKRIENQSNRQVTFSKRKNGILKKAKEISVLCDAEVGVVVFSSAGKLYDFCSTSLSRILEKYQTNSGKI-----
DEKHKSLSAEIDRIKKENDNMQIELRHLKGEDLNSLQPKELIMIEEALDNGLTSLHEKQAL----------------------------------------------
-------------------
>TaFLC.3B_TraesCS3B01G469700
ARRGRVELRRIEDRTSRQVRFSKRRAGLFKKAFELAVLCDAEVALLVFSPAGKLYEYSTSSIEGTYDRYQRFAGAGTNVN---GGDASSNNDGDPS---
PRLKEIASWSLQNNADDSDANKLEKLEKLLTDALRNTKSKK---ML-------------AQQNSGSSTR------------------------------------S-
N
>TaFLC.4B2_TraesCS3B01G470000
ARRGRVELRRIEDRTSRQVRFSKRRSGLFKKAFELAVLCDAEVSLLVFSPAGRLYEYASSSIEGTYDRYQAFAGAGKDVN---EPGASNNNDGDPS---
SRLEEITTWSLQNNADDSDANELEKLEKLLTDALKNTKSKK---ML-------------AQRNSGAGTS------------------------------------S-
S
>TaFLC.6B_TraesCS3B01G683500LC
AQRGRVELRRIEDRTSRQVRFSKRRSGLFKKAFELSLLCDAEVALLVFSPAGKLYEYASSRCTPAATHF--------------------------------------
--------------------FLLTPLQS------------------------------------------------------------------------
>TaBS.8D2_TraesCS3D01G045700LC
---------MVGDDPSRRATFGERNGGLLREAQELAVFCEVEVGILVFDSAGRQLDYCSTRSVSFHRRSSS---------------------------------
SFRLYIDTDYG-FSITSICNLILQCPC----------------------------------------------------------------------------
>TaBS.4D1_TraesCS3D01G090300
MGRGRTEMKKIQNDVSRRATFGKRRRGLLKKAHELAVLCGVDLGLLVFDDAGKLFDYCATSWSELIERYESITKHQQGQGID--
HQQPSADIAGLRREGDHLEASVTRQTGENLSSATAAELDELEQRLECVLGKVRETKDKLLEQQLGESHHKVHILEDQNSFLRRLMGEEGQQRAAASAV---------
-PEELTTLRL-WP---
>TaBS.4D2_TraesCS3D01G090600
MGRGRTEMKRIHNDVSRRATFGKRRRGLLKKARELAVLCGVDHGLLVFDDAGKLFDY----WSELIERYESITKHQQGQGIDH-
QQQPLADIAGLRCEHDRLEASMRRQTGEDLPSATAAELDDLEQRLEFMLSKVREMQARQMEG---------------KSFAE-------------------------
FPLLGFPMSLAWSQVF
>TaBS.9D_TraesCS3D01G102700
MGRGKVEMKRIDNKLSRQATFHKRRRGLLKKAQELAVLCDAHLGVLVFSSTGELYDYCSTSWSELIQRYKGITNAQ--QGTD--
DQKLLEDIARLKRERDHLVASHRMMTGEDMPCTTKEELADLEQKLECALGKVREMKDKFLNQQLEESRHKVSILEEKNSLMRHVMNHDEQPRAVA-----
ELAFGSFFPEESTSLQL-WPQDP
>TaAG.2D_TraesCS3D01G140200
MGRGRIEIKRIENTTNRQVTFCKRRNGLLKKAYELSVLCDAEVALIVFSSRGRLYEYSNNSVKATIERYKKANSDTSNSGTVNA-
QYYQQESSKLRQQISSLQNSNRSLVRDSVSTMTLRDLKQLEGRLEKGIAKIRARKNELMYAEVEYMQKREMELHNDNIYLRSKVSENEQPMNMSGSTSSEYQQQQHY
SQLPTALQLGQQN--
>TaMADS32D_TraesCS3D01G284200
MGRGRSEIKRIDNPTQRQSTFYKRRDGLFKKARELAVLCDADLLLLLFSASGKLYQYLAPSVKEFVERYEAATHTK-----
SDIRQERRAELEKVAKMCDLLEKELRFMTVDDGEQYTVPSLAALEHNLEAAMHKVRSEKDRKIGGEMSYLENMIRGKQAERYGLCDKLAH--QSLKVGGSTSLNN--
-------GLDLKLGFN---
>TaPI.2D_TraesCS3D01G401700
MGRGKIEIKRIENQSNRQVTFSKRKNGILKKAKEISVLCDAEVGVVVFSSAGKLYDFCSTSLSRILEKYQTNSGKI-----
DEKHKSLSAEIDRIKKENDNMQIELRHLKGEDMNSLQPKELIMIEEALDNGLTSLHEKQMEHYDRL----VKHGKMLEDENKLLAFKVH---
QHDIAGSMRDLELDFAAQMPI---TFRV-QPQED
>TaFLC.3D_TraesCS3D01G427700
ARRGRVELRRIEDRTSRQVRFSKRRAGLFKKAFELAVLCDAEVALLVFSPAGKLYEYASSSIEGTYDLYQRFAGAGTNLN---GGDASSNNDGDPS---
STLKEIASWSIQNNADVSDANKLEKLEKLLTDALRNTKSKK---MS-------------AQQNSGAGT-------------------------------------A-
N
>TaFLC.6D_TraesCS3D01G427900
ARRGRVELRRIEDRTSRQVRFSKRRSGLFKKAFELSLLCDAEVALLVFSPAGKLYEYSSSSIEATYDRYQRFAGPGMNVN---GGDASTNNDGDPS---
SRLEEIASWSLQNNADDSDATKLEKPEKLLTDALRNTKSKK---ML-------------AQQNSGASTS------------------------------------S-
E
>TaFLC.4D1_TraesCS3D01G428000
ARRGRVELRRIEDRTSRQVRFSKRRAGLFKKAFELAVLCDAEVSLLVFSPAGRLYEYASSSIEGTYDRYQAFAGAGKDVN---EPGASNNNDGDPS---
SRLEEITSWSLQNNADNSDANELEKLEKLLTDALKNTKSKK---ML-------------AQQNSDAGTS------------------------------------S-
R
>TaFLC.5D_TraesCS3D01G514600LC
```

-RRGRVELRRIEDRTSRQVRFSKRRAGLFKKAFELSLLCDAEVALLVFSPAGKLYEYASSSIEGTYDRYQQFAVPGRNLI---QEDATVCNDEDPS---
SRLGEIAAWSLDNNADNSDASSLEKLEKLLKDALRITESKK---AL-------------AKQNSGGST-------------------------------------P-
N
>TaFLC.1D3_TraesCS3D01G517300LC
-----MELRRIEDRTSRQVRFSKRRSGLFKKAFELAVLCDVEVR------AARL-----------------------------------------------------
--------------LPRRQALRVRILQSMCSAC---IW-------------FRQSSVAEAT---------------------------------------
>TaSVP.3A_TraesCS4A01G002600
-------------------------------------------------------------MNQIIDRYNSHSKIKKADESQLDLH-
EDSNCARLRDELAEASLWLQQMRGEELQSLNVQQLQALEKSLESGLGSVLKTKSQKIMDQISELERKRVQLIEENARLKEQASKM----E-EGQSSESVSYPRP--
PSDTSLRLGLPNSK
>TaSEP1.3A1_TraesCS4A01G028100
MGRGKVEIRRIENKTTRQVTFTKRRNGLLKKAYELSLLCDAEVALIIFSGGGRLFEFSSSCMYKILERYRTCNYNSPEAT-
PAENEINYQEYLKFKTRLEYLQSSQRNILGEDLGPLTMRELEQIENQIDISLKHIRTRKNKVLLDELYDLKSKEQELLDQNKDLRKKLHDISAENALDGGHGSSS-
YPGLLQR-DSSMQIGY--DM
>TaSEP1.3A2_TraesCS4A01G028200
MGRGKVEIRRIENKTTRQVTFTKRRNGLLKKAYELSLLCDAEVALIIFSGGGRLFEFSSSCMYKILERYRTCNYNSPEAT-
PAENEINYQEYLKFKTRLEYLQSSQRNILGEDLGPLTMRELEQIENQIDISLKHIRTRKNKVLLDELYDLKSKEQELLDQNKDLRKKLHDISAENALDGGHGSSS-
YPGLLQR-DSSMQIGY--NL
>TaBS.8A_TraesCS4A01G044400LC
IGRG-----MVDGDASRWATFGERNGGLVREAQELVVLCEAEVDILVFDSAGRQLDYCSTSYMELDT-----------------AKRISVE----------------
------------------------GLERFREVKYYLILIIVDTSGHHAALFEEM------------RQVAE--------SFIEIFAD----LMV-FPMIF
>TaSEP1.2A_TraesCS4A01G058900
MGRGKVEMRRIENKISRQVTFAKRRNGLLKKAYELSLLCDAEVALIIFSGRGRLFEFSSSCMYKTLERYRTCNSNSQEAA-PLENE---------------------
-----------------------------------------------EQELQDENKDLRKKLQDTTGENAVDGGQSSSR----VLQH-DTSMQIGY--HV
>TaSEP1.1A_TraesCS4A01G078700
MAIRPQEMRQIADGTGK-----RRRNPRVGLAY----------------GRG-------NSMYKTLERYRTCNCNSQEAT-
LAENEINYQQYLKLKTRLEYLESSQRNILGEDLGPLSMKELEQIENQIDISLKHIRTRKNKVLLDELYDLKSKEQELQDQNKDLRKKLQDTSAENALDRGQSSSS-
YPGLVQH-DSSMQVGY--DM
>TaBS.5A4_TraesCS4A01G137500
-------------------------------------------------------------------------------------
MKLLADIATLRHERDHLEASVRRQTGEDLPSATAAELRSLEHKLECALGKVRETKDKLMEKQLDESHHRVHILEEQNSFLGHMILKVLKHNLCPAKC----------
---------------
>TaMIKC.1A_TraesCS4A01G169000
MGRVKLKIKKLENTSGRQVTYSKRRSGILKKAKELSILCDIDLILLMFSPSGRPTICVGSPIDEVIAKYAQQTPQERA---
LESLEELSSHLGALQCQMADVQKRLS-YWSDPEKVENIDHIRAMEQSLKESLNRIGIHKENFAKQHLMGLQCAAAQFQNE------QLPL-------
SGGGSDGQEQQH--QQQLTSLHLGQF---
>TaMIKC.2A_TraesCS4A01G290700
MGRVKLAIKRIENNTNRHVTFSKRRNGLIKKAYELSVLCDIDIALLMFSPSRRLCPFSGHGVEDVLLRYLNMSDNDRGEP--
QNREEIQKEIYACQQQLQISEERLRTLFPDPAAFGSTGDIDNCEKFLMEMLTRVVERKNYLLS-NLAPFDPTAPGMQGG------GAQMVH-----EDGQNPGH---
----DTLSTLCLGDDESG
>TaMIKC.1B_TraesCS4B01G150200
MGRVKLKIKKLENTSGRQVTYSKRRSGILKKAKELSILCDIDLILLMFSPSGRPTICVGSPIDEVIAKYAQQTPQERA---
LESLEELSSHLGALQCQMADVQKRLS-YWSDPEKVENIDHIRAMEQSLKESLNRIGIHKENFAKQHLMGLQCPAAQFQNE------QLPL-------
NGGSSDGQEQQH--QQQLTSLHLGQF---
>TaSEP1.2B_TraesCS4B01G245700
MGRGKVEMRRIENKISRQVTFAKRRNGLLKKAYELSLLCDAEVALIIFSGRGRLFEFSSSCMYKTLERYRTCNSNSQEAT-
PLESEINYQEYLKLKTRVEFLQSSQRNILGEDLGPLSMKELDQIENQIDASLKHIRSKKNQVLLDQLFELKSKEQELQDENNDLRKKLQDTTGENAVDGGQCSSR--
---VLH-DTSMQIGY----
>TaSEP1.1B_TraesCS4B01G245800
MGRGKVEMRRIENKISRQVTFAKRRNGLLKKAYELSLLCDAEVALIIFSGRGRLFEFSSSCMYKTLERYRTCNCNSQEAT-
LAENEINYQQYLKLKTRLEYLESSQRNILGEDLGPLSMKELEQIENQIDISLKHIRTRKNKVLLDELYDLKSKEQELQDQNKNLRKKLQDTSVENALDAGQCSSS-
YSGLVQH-DSSVQVGC----
>TaSEP1.3B_TraesCS4B01G277800
MGRGKVEIRRIENKTTRQVTFTKRRNGLL-KAYELSLLCDAEVALVIFSGGGRLFEFSSSCMYKILERYRTCNHNSPEAT-
SAENEINYQEYLKLKTRLEYLQSSQRNILGEDLGPLSMRELEQIENQIDISLKHIRTRKNKVLLHELYDLKSKEQELLDQNKDLRKKLQDISAENALDGGHGSSS-
YPGLLQR-DSSRQIGY--AM
>TaSVP.3B_TraesCS4B01G302600
GKRERIAIRRIENLAARQVTFSKRRRGLFKKAGELSILCDAEVGLAVFSATGKLFQFASSSMNQIIDRYNSHSKIKKADESQLDLH-
EDSNCARLRDELAEASLWLQQMRGEELQSLNVQQLQALEKSLESGLGSVLKTKSQKIMDQISELERKRVQLIEENARLKEQASKM----E-EGQSSESVSYPRP--
PSDTSLRLGLPNSK
>TaSOC1.3B_TraesCS4B01G346700
MVRGKTQMKRIENPTSRQVTFSKRRGGLLKKAFELSVLCDAEVALVVFSPRGRLYEFTSSSMKNTIERYKTVTKDNMSRQVQQDMEKIKADAEGLSKKLDALEACKS
KLLGQNLEECSIEELQSLEVKIEKSLLGIRAMKTRRFEEQLSTLRQKEMTLRQHNEELYSQCQKEQHLASEQG--QQM---------DETDLFLGLPGRS
>TaFLC.2B_TraesCS4B01G351500
ARRGPVELRRIEDRTSRQVRFSKRRSGLFKKAFEFGLLYDAEVALLVFSPAGRLYEYASSSIEDTYDRYQAFGGAGKNLN---EGGASTNNGGDPS---
SRLKEIASWSLQNNADDADANELEKLEKLLTDALENIKSKK---ML-------------AQRNSGAS--------------------------------------T-
I
>TaMIKC.2D_TraesCS4D01G021100
MGRVKLAIKRIENNTNRHVTFSKRRNGLIKKAYELSVLCDIDIALLMFSPSRRLCPFSGHGVEDVLLRYLNMSDNDRGEP--
QNREEIQKEIYACQQQLQISEERLRLFEPDPAAFGSMVEIDNCEKFLMDMLTRVVERKNYLLS-NLAPFDPTAPGMQGG------EAQMVH-----DDGQNPGH---
----DTLSTLCLGDDESG
>TaMIKC.1D_TraesCS4D01G147000
MGRVKLKIKKLENTSGRQVTYSKRRSGILKKAKELSILCDIDLILLMFSPSGRPTICVGSPIDEVIAKYAQQTPQERA---
LESLEELSSHLGALQCQMADVQKRLS-YWSDPEKVENIDHIRAMEQSLKESLNRIGIHKENFAKQHLMGLQCAAAQFQNE------QLPL-------
SGGGSDGQEQQH--QQQLTSLHLGQF---
>TaSEP1.2D_TraesCS4D01G243700
MGRGKVEMRRIENKISRQVTFAKRRNGLLKKAYELSLLCDAEVALIIFSGRGRLFEFSSSCMYRTLERYRTCNSNSQEAT-
PLENEINYQEYLKLKTRVEFLQSSQRNILGEDLGPLSMKELDQIENQIDASLKHIRSKKNQVLLDQLFELKSKEQELQDENNDLRKKLQDTTGDNAVDGGQCSSR--
---VLH-DTSMQIGY----
>TaSEP1.1D_TraesCS4D01G245200
MGRGKVEMRRIENKISRQVTFAKRRNGLLKKAYELSLLCDAEIALIIFSGRGRLFEFSSSCMYKTLERYRTCNCNSQEAT-
LAENEINYQQYLKLKTRLEYLESSQRNILGEDLGPLSIKELEQIENQIDISLKHIRTRKNKVLLDELYDLKSKEQELQDQNKNLRKKLQDTSAQNAPDAGQSSSS-
YPGLVEH-DSSMQVGY--DM
>TaSEP1.3D_TraesCS4D01G276100
MGRGKVEIRRIENKTTRQVTFTKRRNGLLKKAYELSLLCDAEVALVIFSGGGRLFEFSSSCMYKILERYRTCNYNSTEAT-
PAENEINYQEYLKLKTRLEYLQSSQRNILGEDLGPLSMRELEQIENQIDISLKHIRTRKNKVLLDELYDQKSKEQELLDQNKDLRRKLQDISAENALDG-------
YPGLLQR-DSSMQIGY--DM
>TaSVP.3D_TraesCS4D01G301100

GKRERIAIRRIENLAARQVTFSKRRRGLFKKAEELSILCDAEVGLAVFSATGKLFQFASSSMNQIIDRYNSHSKIKKADESQLDLH-
EDSNCARLSDELAEASLWLQQMRGEELQSLNVQQLQALEKSLESGLGSVLKTKSQKIMDQISELENKRVQLIEENARLKEQASKM----E-EGQSSESVSYPRP--
PSDTSLRLGLPNSK
>TaSOC1.3D_TraesCS4D01G341700
MVRGKTQMKRIENPTSRQVTFSKRRGGLLKKAFELSVLCDAEVALVVFSPRGRLYEFASSSMKNTIERYKTVTKDNLGRQVQQDIEKVKADAEGLSKKLDALEACKS
KLLGQNLEECSIEELQSLEVKIERSLLGIRAMKTRRFEEQLSTLRQKEMKLRQDNEELYSQCQKEQHSALEQG--QQV---------DETDLFLGLPGRS
>TaFLC.2D_TraesCS4D01G346300
ARRGPVELRRIEDRTSRQVRFSKRRSGLFKKAFELGLLCDAEVALLVFSPAGRLYEYASSSIEDTYDRYQAFGGAGKNLN---EGGASTNSDGDPS---
SRLKEIASWSLQNNADDADASELEKLEKLLTDALENTKSKK---ML-------------AQRNSGARTS------------------------------------S-
I
>TaAGL17.2A3_TraesCS5A01G001300LC
----------------------------------------------------------------------------------
SVGQLWQREVTTLRQQVHNLQHNNRQLLGEELSGSTVRDLQFLVNQLETSLHCVRKRKEQVMAEEIHELNQKGFLIQKENIELGKKVRITHEQNIEAMRSSEQQSSS
KAAAGRFIDLELRQQEDE
>TaAGL17.11A1_TraesCS5A01G001400LC
MVRGKIVIRRIEDMSSRQVTFSKRRHGLLKKTRELAILCNVQVGVIIFSLTGRIYAYASSTLP-----------------------PWA------------------
----------------CLPSSISTNLHR---------------------------------------------------SINY-----------------
>TaAGL17.2A2_TraesCS5A01G001800
------MIERIDNATNRQLTFSKQRGGLMKKAWELAILCEADLALIVFSSTGRLYDFASSR--PVVN----------------------------------------
----------------------------------------------------------------------------------------------------
>TaSTK.1A_TraesCS5A01G117500
MGRGRIEIKRIENTTSRQVTFCKRRNGLLKKAYELSVLCDAEVALIVFSSRGRLYEYSNNSVKATIDRYKKAHACGSTSGPVNAQQYYQQEAARLRHQIQMLQSTNK
HLVGDSVGNLSLKELKQLESRLEKGIAKIRARKNELLSSEINYMVKREIELQSDNIDLRTKIAEEEQQVTISAAP--ELAHAQAQAQQSLQLNLGY-QLA
>TaSEP3.2A_TraesCS5A01G286800
MGRGRVELKRIENKINRQVTFAKRRNGLLKKAYELSVLCDAEVALIIFSNRGKLYEFCSQSMPKTLERYQKCSYGGPDTAQNELVQSSRNEYLKLKARVENLQRTQR
NLLGEDLGSLGIKDLEQLEKQLDSSLRHIRSTRTQHMLDQLTDLQRKEQMLCEANKCLRRKLEESSQMQG-HAANLLGYGGNGFFHPTEPTLQIGY----
>TaAP1.1A_TraesCS5A01G391700
MGRGKVQLKRIENKINRQVTFSKRRSGLLKKAHEISVLCDAEVGLIIFSTKGKLYEFSTSCMDKILERYERYSYAEKVLVSSEIQGNWCHEYRKLKAKVETIQKCQK
HLMGEDLESLNLKELQQLEQQLESSLKHIRSRKNQLMHESISELQKKERSLQEENKVLQKELVEKQKA-----------SSSFMLRD-PPAANTS--AAV
>TaSEP1.6A_TraesCS5A01G391800
MGRGKVVLQRIENKISRQVTFAKRRNGLLKKAYELSLLCDAEVALVLFSHAGRLYQFSSSNMLKTLEKYQRYIFASQDAAPTDEMQNNYLEYMELKSRVEVLQRSQR
NLLGEDLAPLSTIELEQLEGQVGKTLRQIRSRKTQVLLDEMCDLKRKEQILQDANMTLKRKLGEIELEATPDR------QPEHFFQA-YPSLQPVF----
>TaSOC1.3A_TraesCS5A01G515500
MVRGKTQMKRIENPTSRQVTFSKRRGGLLKKAFELSVLCDAEVALVVFSPRGRLYEFASSSMKNTIERYKTVTKDNISRQVQQDIEKIKADAEGLSKKLDALEACKS
KLLGQNLEECSIEELQSLELKIEKSLLGIRAVKTRRFEEQLSTLRQKETKLRQDNEELYSQCQKEQHLAL-QS--QDV---------DETDLFLGLPGRS
>TaFLC.2A_TraesCS5A01G520200
ARRGPVELRRIEDRTSRQVRFSKRRSGLFKKAFELGLLCDAEVALLVFSPAGRLYEYASSSIEDTYDRYQAFGGAGKNLN---EGGASTNNDKDPS---
SRLKEIASWSLQNNADDSDANELEKLEKLLTDALENTKSKK---ML-------------AQRNRGAR--------------------------------------S-
I
>TaAGL17.2B_TraesCS5B01G002400
MGRGKIAIERIDNATNRQVTFSKRRGGLMKKARELAILCDADLALIIFSSTGRLYNFASSSMEAILERYQEAKQEHCGVLPTSEAKLWQREVTTLRQQVQNLQHNNR
QLLGEELSGSTVRDLQFLVNQVEMSLHSVRKRKEQVIAEEIHELNQKGFLIQKENIELGKKLSIAHKRNIEVMRTSEQQSSSKAAAGRFIDLELRQQEDE
>TaSTK.1B_TraesCS5B01G115100
MGRGRIEIKRIENTTSRQVTFCKRRNGLLKKAYELSVLCDAEVALIVFSSRGRLYEYSNNSVKATIDRYKKAHACGSTSGPVNAQQYYQQEAAKLRHQIQMLQSTNK
HLVGDSVGNLSLKELKQLESRLEKGIAKIRARKNELLSSEINYMVKREIELQSDNIDLRTKIAEEEQQVTISAAL--ELAHAQARTQQSLQLNLRY-QLA
>TaSEP3.2B_TraesCS5B01G286100
MGRGRVELKRIENKINRQVTFAKRRNGLLKKAYELSVLCDAEVALIIFSNRGKLYEFCSQSMPKTLERYQKCSYGGPDTAQNE---
SSRNEYLKLKARVENLQRTQRNLLGEDLGSLGIKDLEQLEKQLDSSLRHIRSTRTQHMLDQLTDLQRKEQMLCEANRCLRRKLEESSQMQG-
HAANLLGYGGNGFFHPTEPTLQIGY----
>TaAP1.1B_TraesCS5B01G396600
MGRGKVQLKRIENKINRQVTFSKRRSGLLKKAHEISVLCDAEVGLIIFSTKGKLYEFSTSCMDKILERYERYSYAEKVLVSSEIQGNWCHEYRKLKAKVETIQKCQK
HLMGEDLESLNLKELQQLEQQLESSLKHIRSRKNQLMHESISELQKKERSLQEENKVLQKELVEKQKA-----------SSSFMMRD-PPAATTS--AAV
>TaSEP1.6B_TraesCS5B01G396700
MGRGKVVLQRIENKISRQVTFAKRRNGLLKKAYELSLLCDAEVALVLFSHAGRLYQFSSSNMLKTLERYQRYIFASQDAAPRDEMQNNYLEYMELKARVEVLQHSQR
NLLGEDLAPLSTTELDQLESQVGKTLRQIRSRKTQVLLDEMCDLKRKEQILEDANMTLKRKLGEIELEATPDR------QPEHFFQA-YPSLQPVF----
>TaAGL17.10D_TraesCS5D01G002100
MARGKIVIRRIEKMSSRQVTFSKRRRGLLKKARELAILCDVQVGVIVFSSTGRLYEYASSTMPSIIHKYQSAQEHQQLL-
PVSQIMFWEGEVRKLQQEMQMLQEHHRTLMGERLSNLGVNDLCLLENQLEKSLRCIREKK---------------------
NIQLSEEIKLIHERNLEKESSNRENNVPATPTSTIMGFQITTN---
>TaAGL17.2D_TraesCS5D01G002200
MGRGKIAIERIDNATNRQVTFSKRRGGLMKKARELAILCDADLALIVFSSTGRLYDFASSSMEAILERYQEAKQEHCGVLPTSEAKLWQREVTTLRQQVQNLQHNNR
QLLGEELSGSTVRDLQFLVNQVEMSLHSVRKRKEQVMAEEIHELNQKGFLI-KENIELGKKLSIAHERNIE--------------------------EDE
>TaSTK.1D_TraesCS5D01G118200
MGRGRIEIKRIENTTSRQVTFCKRRNGLLKKAYELSVLCDAEVALIVFSSRGRLYEYSNNSVKATIDRYKKAHACGSTSGPVNAQQYYQQEAAKLRHQIQMLQSTNK
HLVGDSVGNLSLKELKQLESRLEKGIAKIRARKNELLSSEINYMVKREIELQSDSIDLRTKIAEEEQQVTISVAP--ELAHAQAQAQQSLQLNLGY-QLA
>TaSEP3.2D_TraesCS5D01G294500
MGRGRVELKRIENKINRQVTFAKRRNGLLKKAYELSVLCDAEVALIIFSNRGKLYEFCSQSMPKTLERYQKCSYGGPDTAQNELVQSSRNEYLKLKARVENLQRTQR
NLLGEDLGSLGIKDLEQLEKQLDSSLRHIRSTRTQHMLDQLTDLQRKEQMLCEANKCLRRKLEESSQMQG-HAANLLGYGGNGFFHPTEPTLQIGY----
>TaAP1.1D_TraesCS5D01G401500
MGRGKVQLKRIENKINRQVTFSKRRSGLLKKAHEISVLCDAEVGLIIFSTKGKLYEFSTSCMDKILERYERYSYAEKVLVSSEIQGNWCHEYRKLKAKVETIQKCQK
HLMGEDLESLNLKELQQLEQQLESSLKHIRSRKNQLMHESISELQKKERSLQEENKVLQKELVEKQKA-----------SSSFMMRD-PPAAATS--AAV
>TaSEP1.6D_TraesCS5D01G401700
MGRGKVVLQRIENKISRQVTFAKRRNGLLKKAYELSLLCDAEVALVLFSHAGRLYQFSSSNMFKTLERYQRYIFASQDAVPSDEMQNNYLEYMELKARVEVLQHSQR
NLLGEDLAPLSTTELDQLESQVGKTLRQIRSRKTQVLLDELCDLKRKEQMLQDANMTLKRKLGEIQVEATPDR------QPEHFFQA-YPSLQPVF----
>TaSOC1.2A_TraesCS6A01G011700
MARGRREMRRIEDATSRQVTFSKRRSGLLKKAFELGVLCDAEVALIVFSPRGRLYEYASAPLQKTIDRYLNHTKGSTNEKVEQGVQMCRSEATALKHKIDAIEAYQR
KLSGEGLGSCSAHELQELELQLEKSLSCIRQKKQQKMLDKILELKEKERKLLTENSVLRKEYKALPLLEL-EAEEDERR--------EETELVIGRPSS-
>TaAP3.2A_TraesCS6A01G131100
MGRGKFEIKRIQDATGRQVCYSKRRSGIMKKARELAVLCDAQVAVVVLSSTGKHHHFCSADIKGILNRYQQATGTS-----
TEQYEEMQRTLSRLKDINRNLRTEIRQRMGEDLDALEFGELRGLEQSVDAALEVVRQRKYQVITRQTETYKRKVKHSQKVYKSLQQELSMREDPVFGDNPAFGFM--
------EMYAFHA-VSHGM
>TaBS.1A_TraesCS6A01G158100
MGRGKIEIKRIENATNRQVTFSKRRGGLLKKANELAVLCDARVGVVIFSSTGRMFEYSSSSLRDLIEQYQNATNSQ--EEI---
DQQIFVEMTRMRNEMEKLDGAIRRYTGDDLSSLSLADVNDIEQQLEFSVAKVRARKHQLLNQQLDNLRRKEHILEDQNSFLCRMISENQH----
GGDGKMAVMLTPAFPAESTALQL-TSQDP
>TaAGL6A_TraesCS6A01G259000
MGRGRVELKRIENKINRQVTFSKRRNGLLKKAYELSVLCDAEVALIIFSSRGKLYEFGS-GTTKTLERYQHCCYNAQDSN-
LSETQSWYQEMSKLKAKFEALQRTQRHLLGEDLGPLSVKELQQLEKQLECSLSLARQRKTQLMMEQVEELRRKERQLGDINRQLKHKLDAEGNSNNYAGTVVDE-
QHPNHSAA-EPTLQIGY-RSS
>TaAGL17.3A_TraesCS6A01G292300

MGRGKIVIRRIDNSTNRQVTFSKRRGGLLKKAKELSILCDAEVGLVVFSSTGRLHEFSSTNMKAVIDRYTKAKEEQAGV-
ATSEIKLWQREAASLRQQLHDLQESHKQLMGEELSSLGVRDLQGLENRLEMSLRSIKTRKDNLLRSEIEELHRKGSLIHQENTELCRRLNIMSQQKMESGGATDANS
TPYSFRINPANLELSQAEQE
>TaSVP.1A_TraesCS6A01G313800
MARERREIKRIESAAARQVTFSKRRRGLFKKAEELSVLCDADVALIVFSSTGKLSQFASSSMNEIIDKYSTHSKNGKTD-
PALDLNLEHSKYANLNDQLAEASLRLRQMRGEELEGLSVDELQQLEKNLETGLHRVLQTKDQQFLEQINELHRKSSQLAEENMKLRNQVGQIPTAGK-
DGQSSESVLHSGS--SGDVSLKLGLPPWK
>TaBS.6A1_TraesCS6A01G325500LC
-----------SNNVSRQSIFGKSRAGLLKKAQELAMLCEADLGVLIFDSAGKQMDYCSTMSNSSPSIFSNQSHGT--------------------------
SSRLQTYFGYSLARCPSAALTCI--LLDCSVFSTR-----------------VHACGELIDSFRKMNLLK-------------------------SLEI-LTLGQ
>TaSOC1.2B_TraesCS6B01G017900
MARGRREMRRIEDATSRQVTFSKRRSGLLKKAFELGVLCDAEVALIVFSPRGRLYEYASAPLQKTIDRYLNHTKGSANEKVEQGVQMCRSEATALKHKIDAIEAYQR
KLSGEGLGSCSAHELQELELQLEKSLSCIRQKKQQKMVAKISELKEKERKLLTENSVLREEYKALPLLEL-EAEEDERR--------EETELVIGRPSS-
>TaAP3.2B_TraesCS6B01G159400
MGRGKFEIKRIDDATSRQVCYSKRRSGIMKKARELAVLCDAQVAVIVLSSSGKHHHFCSADIKGVFDRYQQATGTS-----
TEQYENMQRTLSQLKDINRNLRTEIRQRMGEDLDALEFQELRGLEQNVGAALEVVRQRKYHVITRQTETYKKKVKHSEEVYKKLKQELSMREDP------AFGFM--
------EMYAFHA-VSHGM
>TaBS.1B_TraesCS6B01G186700
MGRGKIEIKRIENATNRQVTFSKRRGGLLKKANELAVLCDARVGVVIFSSTGRMFEYSSSSLRDLIEQYQNATNSQ--EEI---
DQQIFVEMTRMRNEMEKLDGAIRRYTGDDLSSLSLADVNDIEQQLEFSVAKVRARKHQLLNQQLDNLRRKEHILEDQNSFLCRMISENQH----
GSDRKMAVMLTPAFPAESTALQL-TSQDP
>TaAGL6B_TraesCS6B01G286400
MGRGRVELKRIENKINRQVTFSKRRNGLLKKAYELSVLCDAEVALIIFSSRGKLYEFGS-GTTKTLERYQHCCYNAQDSN-
LSETQSWYQEMSKLKAKFEALQRTQRHLLGEDLGPLSVKELQQLEKQLECSLSLARQRKTQLMMEQVEELCRKERQLGDINRQLKHKLDAEGNSNNYAGTVVDE-
QHPNHSAA-EPTLQIGY-MPQ
>TaAGL17.3B_TraesCS6B01G322700
MGRGKIVIRRIDNSTNRQVTFSKRRGGLLKKAKELSILCDAEVGLVVFSSTGRLHEFSSTNMKAVIDRYTKAKEEQPGV-
ATSEIKLWQREAASLRQQLHDLQESHKQLMGEELSGLGVRDLQGLENRLEMSLRSIKTRKDNLLRSEIEELHRKGSLIHQENTELCRRLNIMSQQKMESGGATDANS
TPYSFRINPANLELSQAEQE
>TaSVP.1B_TraesCS6B01G343900
MARERREIKRIESAAARQVTFSKRRRGLFKKAEELSVLCDADVALIVFSSTGKLSQFASSSMNEIIDKYSTHSKNGKTD-
PALDLNLEHSKYANLNDQLAEASLRLRQMRGEELEGLSVDELQQLEKNLETGLHRVLQTKDQQFLEQINELHRKSSQLAEENMKLRNQVGQIPTAGK-
DGQSSESVLHSGS--SGDVSLKLGLPPWK
>TaSOC1.2D_TraesCS6D01G014600
MARGRREMRRIEDATSRQVTFSKRRSGLLKKAFELGVLCDAEVALIVFSPRGRLYEYASAPLQKTIDRYLNHTKGSANEKVEQGVQMCRSEATALKHKIDAIEAYQR
KLSGEGLGSCSAHELQELELQLEKSLSCIRQKKQQKMLDKILELKEKERKLLTENVVLREEYKALPLLEL-EAEEDERR--------EETELVIGRPSS-
>TaAP3.2D_TraesCS6D01G120900
MGRGKFEIKRIEDATSRQVSYSKRRSGIMKKARELAVLCDAQVAIVMLSSTGKHHHFCSADIKGIFDRYQQATGTS-----
TEQYENMQRTLSRLKSINRNLRTEIRQRMGEDLDALEFKELRGLEQNVGAALEVVRQRKYHVITRQTEIYKKKVKHSEEVYKKLQQELSMREDPAFGDNRAFGFM--
------EMYAFHA-VSHGM
>TaBS.1D_TraesCS6D01G147400
MGRGKIEIKRIENATNRQVTFSKRRGGLLKKANELAVLCDARVGVVIFSSTGRMFEYSSSSLRDLIEQYQNATNSQ--EEI---
DQQIFVEMTRMRNEMEKLDGAIRRYTGDDLSSLSLADVNDIEQQLEFSVAKVRARKHQLLNQQLDNLRRKEHILEDQNSFLCRMISENQH----
GGDGKMAVMLTPAFPAESTALQL-TSQDP
>TaAGL6D_TraesCS6D01G240200
MGRGRVELKRIENKINRQVTFSKRRNGLLKKAYELSVLCDAEVALIIFSSRGKLYEFGS-GTTKTLERYQHCCYNAQDSN-
LSETQSWYQEMSKLKAKFEALQRTQRHLLGEDLGPLSVKELQQLEKQLECSLSLARQRKTQLMMEQVEELRRKERQLGDINRQLKHKLDAEGNSNNYAGTVVDE-
QHPNHSAA-EPTLQIGY-VPQ
>TaAGL17.3D_TraesCS6D01G273500
MGRGKIVIRRIDNSTNRQVTFSKRRGGLLKKAKELSILCDAEVGLVVFSSTGRLHEFSSTNMKAVIDRYTKAKEEQPGV-
ATSEIKLWQREAASLRQQLHDLQESHKQLMGEELSSLGVRDLQGLENRLEMSLRSIKTRKDNLLRSEIEELHRKGSLIHQENTELCRRLNIMSQQKMESGGATDANS
TPFSFRINPANLELSQAEQE
>TaSVP.1D_TraesCS6D01G293200
MARERREIKRIESAAARQVTFSKRRRGLFKKAEELSVLCDADVALIVFSSTGKLSQFASSSMNEIIDKYSTHSKNGKTD-
PALDLNLEHSKYANLNDQLAEASLRLRQMRGEELEGLSVDELQLLEKNLETGLHRVLQTKDQQFLEQINELHRKSSQLAEENMKLRNQVGQIPTAGK-
DGQSSESVLHSGS--SGDVSLKLGLPPWK
>TaSOC1.4A_TraesCS7A01G051700LC
MGRGKTQMKLIEDRTSRRVAFSKRRSGLHRKAFQLSVLCDAEVTLIVFSPGGRLYEFANAGMQNTIGRYETNTKDSTSNQVHQDIEKIKADAEALSKKLDALEAYKR
KILGSNLEECSIEELQSIEDRTEKKPSYTSASQARRLEEQLAKLRQNVVKLSHHKQELYFQVKMDNEIAF-----------------NTTQL--------
>TaAGL17.4A_TraesCS7A01G111700
MGRGKIAIERIDNTTNRQVTFSKRRGGLMKKARELAILCDADLALIIFSSTGRLYDFASSSMQAILERYQKAKEEHCGVLPTSEAKLWQREVTTLRQQVQNLHHNNR
QLLGEELSGTTIRDLQFLVNQVEMSLHSIRKTKEQVMAAEIHELNQKGFLVQKENVKLDKKFSIAHEQNIEAMRSSEQQSNSKAAAGN-IDLELRQQEDE
>TaAGL17.5A1_TraesCS7A01G111800
MGRGKIAIERIDNTMNRQVTFSKRRGGLMKKARELAILCDADLALIVFSSTGRLYDFASSSMEAILERYQEAKEEHRGVLPTSEAKLWQREVTTLRQQVQNLQHNNR
QLLGEELSGTTVRDIQFLVNQVEMSLHSVRKRKEQIMAAEIHELNQKGFLIQKENIELGKKLSIAHEHNIEAMESSEQQSSSKAAAGN-IDLELRQHEDE
>TaAGL17.5A2_TraesCS7A01G111900
MGRGKIAIERIDNTTNRQVTFSKRRGGLMKKARELAILCDADLALIVFSSTGRLYDFASSSMEAILERYQEAKEEHCGVLPTSEAKLWQREVTTLRQQVQNLQHNNR
QLLGEELSGTTVRDLQFLVNQVEMSLHSVRKRKEQIMAAEIHELNQKGFLIQKENIELGKKLSIAHEHNIEAMASSEQQSTSKAAAGN-IDLELRQQEDE
>TaSEP1.4A_TraesCS7A01G122000
MGRGKVELKRIDNKISRQVTFAKRRNGLLKKAYELSVLCDAEVALIIFSTRGRLFEFSTSCMYKTLERYRSCNFNSEATA-P-
ETESNYQEYLKLKTRVEFLQTTQRNHLGEDLGPLNMKELEQLENQIEISLKHIRATKSQQSLDQLFELKRKEQQLQDVNKDLRKKIQETSVENVLDVGPSGSS-
QQEYF-H-DPSLHIGY----
>TaSEP1.5A_TraesCS7A01G122100
MGRGKVELKRIDNKISRQVTFAKRRNGLLKKAYELSLLCDAEVALIIFSARGRLFEFSTSRMYKTLERYRSCNFNSEATA-P-
ETESNYQEYLKLKTRVEFLQTTQRNLLGEDLGPLNMKELEQLENQIEISLKHIRATKSQQSLDQLFELKRKEKQLQDVNKDLRKKIQETGADSVLDVGPSGSS-
QQEYF-H-DPSLRMGY----
>TaAGL17.11A3_TraesCS7A01G151800LC
MGRGKIVIRRIENMSSRQATFSKRCRGLLKKARELAILCDVQVSVIVFSSTGRLYEYASPIMPSIIQNYQSAQEHHQLL-
PISQVMFLEAFFTLNVPYMMFLTSRHA--------------LCIMQL------------------------------------------------------------
--------------------
>TaSVP.2A_TraesCS7A01G175200
MARERRAIRRIESAAARQVTFSKRRRGLFKKAEELAVLCDADVALVVFSSTGKLSQFASSSMNEIIDKYSTHSKNGKSDQPAIDLNLEHCKYDSLNEQLAEASLRLR
HMRGEELDGLSVGELQQMEKNLETGLQRVLCTKDRQFMQQISDLQHKGTQLAEENMRLKNQMHEVPTAST-DAHSSDSVVHSGS--SGDISLKLALP-WK
>TaSEP3.1A_TraesCS7A01G260600
MGRGRVELKRIENKINRQVTFAKRRNGLLKKAYELSVLCDAEVALIVFSNRGKLYEFCSQSMTKTLDKYQKCSYAGPETTQNEQLKNSRNEYLKLKARVDNLQRTQR
NLLGEDLDSLGIKELESLEKQLDSSLKHIRTTRTQHMVDQLTELQRREQMFSEANKCLRIKLEESNQVHG-HNNNVLSYGGNGFFHPAEPTLHIGY----
>TaFLC.1A_TraesCS7A01G260900
RKRGRVELRRIEDRTSRQVRFSKRRSGLFKKAFELSVLCDAEVALVVFSPAGRLYEFVSTSVEQIFGRCKDIPDTIID-----
DLNIAARDSRGYCESVSELNSFAQMALELDVKEMSMAELTRFEEIVREALAAVK------------------ARLRRDPQK--------------------------
--------------W--

>TaAGL12.1A_TraesCS7A01G319400
MARGKVQLRRIENPVHRQVTFCKRRAGLLKKARELSVLCDADIGIIIFSAHGKLYDLATGTMDGLIERYKSASGEGMAADGGDQRVDPKQEAMVLKQEIDLLQKGLR
YIYGNRAEHMNVDELNALERYLEMWMFNIRSAKMQIMIQEIQALKSKEGMLKAANEILQEKIVEQHDVGMT--DQ--QNTNPLTILSYCRGSEMGYS---
>TaAP3.1A_TraesCS7A01G383800
MGRGKIEIKRIENATNRQVTYSKRRSGIMKKARELTVLCDAQVAIIMFSSTGKYHEFCSTDIKGIFDRYQQAIGTS-----
IEQYENMQRTLNHLKDINRNLRTEIRQRMGEDLDALEFEELRDLEQNVDAALKEVRQRKYHVITTQTETYKKKVKHSQEAYKNLQQELGMREDPAYGDNPAAGGW--
------AMYAFRV-VPHGM
>TaBS.9A_TraesCS7A01G474900
MGRGKVEMKRIDNKASRQVTFHKRRRGLLKKAQELAALCDAHLGVLVFDGTGKLHDYCSTRSVKFCHRYEV------------------VVYTQPRTKI--------
----------------YFPLKIASTLHSCSEWTRPLTSVSIEQTRRPRHTKVMLNNIYARL------------------LASGSIGLN--VSGL------
>TaBS.5A1_TraesCS7A01G475000
MGRGRVEIRRIANDVSRRATFGKRSAGLLKKARELAVLCDVDLGVLVFDGSGRHAHYCSTSWTELIQRYDSITNDQGNEGTD--
HQQLLTDIARLRREHDHLEATLQRHTGEDLPSATAAELRDLEQRLECALDKVREMKDKLMEEQLDESHNRMHILEEQNSFLRHMMSEEGRQRAAASAVVAELLFGGF
FPEEVTSLRL-WP---
>TaBS.3A_TraesCS7A01G672200LC
KGRG-----MADDDVSRRATFGERSGGLLTEAQELAALWEADVGVLVFDRAGKQMDYCSTSWSELMQRYQIITKGK--
EGIDDRHQELLAEITRLRHERDRLEASVRRQTGDDLPPAT-
TDLGDLEQQVEYALGKVRVMKDKLLEQQLDESYHRVHILEDQNSFLRHMMSEEGRRRAAASAVVAELLFGGFFPEESTSLRL-WP---
>TaAGL17.4B_TraesCS7B01G009500
MGRGKIAIERIDNTTNRQVTFSKRRGGLMKKARELAILCDADLALIVFSSTGRLYDFASSSMEAILERYQEAKEEHCGVLPTSEAKLWQREVTTLRQQVQNLHYNNR
QLLGEELSGTSVRDLQFLVNQVEMSLHSIRKRKEQVMAAEIHELNQKGLLVQKENIELDKKLSIAHEQNIEAMKSSEQQSSSKAAAGN-IDLELRQQEDE
>TaAGL17.11B1_TraesCS7B01G009600
MVRGKIVIRRIENMSSRQVTSSKRRRGLLKKARELAILCDVQVGVIVFSSTGCLYEYANPP----------------------------------------------
----------------------------------------------------------------------------------------------------
>TaAGL17.5B_TraesCS7B01G009800
MGRGKIAIERIDNTTNRQVTFSKRRGGLMKKARELAILCDADLALIVFSSTGRLYDFASSSMEAILERYQEAKEEHCGVLPTSEAKLWQREVTTLRQQVQNLQHNNR
QLLGEELSGTTVRDLQFLVNQVEMSLHSVRKRKEQVMAAEIHELNQKGFLIQKENIELGKKLSIAHEHNIEAMASGEQQSSSKAAAGN-IDLELRQQEDE
>TaAGL17.11B2_TraesCS7B01G009900
MVRGKIVIRRVENMSRRQVTFSKRRGGLLKKARELAILCNVQVGVIVFSSTCRLYEYANSTMPSMIQNYQSSQEQHQLL-
PVSQVMFWQGEVQKLQQEMQMLQEHHRKLMGERLLNLGVKDLCLMENQLEKSLHRIREKKEQSSTNHILQLNEKGCIFHQENIRLSREIKLIHEQNLE---------
--------------------
>TaSEP1.4B_TraesCS7B01G020800
MGRGKVELKRIDNKSSRQVTFAKRRNGLLKKAYELSVLCDAEVALIIFSTRGRLFEFSTSCMYKTLERYRSCNFNSEATA-P-
ETENNYQEYLKLKTRVEFLQTTQRNLLGEDLGPLNMKELEQLENQIEISLKHIRATKSQQSLDQLFELKRKEQQLQDVNKDLRKKIQETSAENVLDVGPSGSS-
QQQHF-H-DPSMRIGY----
>TaSEP1.5B1_TraesCS7B01G020900
MGRGKVELKRIDNKVSRQVTFAKRRNGLLKKAYELSVLCDAVVALIIFSTRGRLFEFSTS--YMTLERYRSCNFNSEATA-P-KTE--------------------
RNLLGEDLGPLNMKELEQLDSQIEISLKHIRATKSQQSLDQLFELKHKEQQLQDVNKDLRKKIQETSAESVLDVGPSGSS-QQEHF-H-DPSLCMG-----
>TaSEP1.5B2_TraesCS7B01G021100
MGRGKVELKRIDNKSSRQVTFAKRRNGLLKKAYELSVLCDAEVALIIFSTRGRLFEFSTSCMYKTLERYRSCNFNSEATS-P-ESES-
YQEYLKLKTRVDFLQTTQRNLLGEDLGPLNMKELEQLENHIEMSLKHIRATKSQQSFDQLFELKRKEQQLQDVNKDLRKKIQETSAESVLDVGPSGSS-QQQHF-H-
DPSLRMW-----
>TaSVP.2B_TraesCS7B01G080300
MARERRAIRRIESAAARQVTFSKRRRGLFKKAEELAVLCDADVALVVFSSTGKLSQFASSSMNEIIDKYSTHSKNGKSDQPAIDLNLEHCKYDSLNEQLAEASLRLR
RMRGEELDGLSVGELQQMEKNLETGLQRVLCTKDRQFMQQINDLQQKGTQLAEENMRLKNQMHEVPTASM-DVHSSDSVVHSAS--SGDISLKLALP-WK
>TaSEP3.1B_TraesCS7B01G158600
MGRGRVELKRIENKINRQVTFAKRRNGLLKKAYELSVLCDAEVALIVFSNRGKLYEFCSQSMTKTLDKYQKCSYAGPETTQNEQLKNSRNEYLKLKARVDNLQRTQR
NLLGEDLDSLGIKELESLEKQLDSSLKHIRTTRTQHMVDQLTELQRREQMFSEANKCLRIKLEESNQVHG-HNNNVLSYGGNGFFHPAEPTLHIGY----
>TaFLC.1B_TraesCS7B01G158900
RKRGRVELRRIEDRTSRQVRFSKRRSGLFKKAFELSVLCDAEVALVVFSPAGRLYEFVSTSVEQIFGRCKDIPDTVID-----
DLNIAARDPRGYCESVSELNSFAQMALELDVKEMSMAELTRFEEIVTEALAAVKSIR---IMKVAE-LTQTEARLKRDPQK--------------------------
--------------W--
>TaAGL12.1B_TraesCS7B01G220200
MARGKVQLRRIENPVHRQVTFCKRRAGLLKKARELSVLCDADIGIIIFSAHGKLYDLATGTMDGLIERYKSASGEGVAADGGDQRVDPKQEAMVLKQEIDLLQKGLR
YIYGNRAEHMNVDELNALERYLEIWMFNIRSAKMQIMIQEIQALKSKEGMLKAANEILQEKIVEQHDVGMT--DQ--QNTNPLTILSYCRGSEMGYS---
>TaAP3.1B_TraesCS7B01G286600
MGRGKIEIKRIENATNRQVTYSKRRSGIMKKARELTVLCDAQVAIIMFSSTGKYHEFCSTDIKGIFDRYQQAIGTS-----IEQYENMQRTLSHLKDINRNLRTEI-
-RMGEDLDALEFEELRDLEQNVDAALKEVRQRKYHVITTQTETYKKKVKHSQEAYKNLQQELGMREDPAYGDNP-------------AMYAFRV-VPHGM
>TaBS.3B_TraesCS7B01G377700
MGRG-----MADDGVSRRATFGERSGALLTEAQELAALWEADVGVLVFDGAGRQMDYCSTSWSELMQRYQIITKGK--
EGIDDRHQQLLAEITRLRRERDRLEASIRRQTGDDLPSAT-
AGLGDLEQQVERALGKVRETKDKLLEQQLDEIHHRVHILEDQNSFLRHMMSEEGRQRAAASAVVAELLFGGFFPEESTSLRL-WPGN-
>TaBS.9B_TraesCS7B01G633300LC
MGRGKVVMKRIDNKVARQVTFHKRRRGLLKKAQELAVLCDAHLGVLVFNSTGKLHDYCSTRSVCLLATFLT-------------
LHGPFVRPSRFYSGRCILILGTRLFGRTDL-------------FLQFGYYSLR------------------SILWNGGSSMQNLISCIVRKKV--------------
----------------
>TaAGL17.6B2_TraesCS7B01G793900LC
MVRGKAVIEKIENQTSRQVTFSKRKSRLFKKGKELGVLCDAQVGIFIFSNTGRLYEYSNS-MKPLIERYQAVKDGQKLL-
ASAEAKFWQAESARLEQQLRTLQENHR--------------------------YD----------------------------------------------------
--------------------
>TaAGL17.6B3_TraesCS7B01G794200LC
MVRGKAVIEKIENQTSRQVTFSKRKSGLFKKGKELGVLCDAQVGIFIFSNTGRLYEYSNS-MKPLIERYQAVKDGQKLL-
ASAEAKFWQAETARLEQQLRTLQENHRQLLGQHLSGLSFEDLKHLQNHLETSLYNIRLTKDKFIGDEMQELNMNESLMRQENIELHRKLNIVHQENTEQGAVNMGIF
TKCNITAKSVRLDSCH-ADW
>TaSOC1.4D_TraesCS7D01G037000
MGRGKTQVKLIEDRTSQQVPFSKLRSGLHRKAFQLSVLCDAEIALVVFSPGGRLYEFANAG----------------------------------------------
----------------------------------------------------------------------------------------------------
>TaAGL17.4D_TraesCS7D01G106600
MGRGKIAIERIDNTTNRQVTFSKRRGGLMKKARELAILCDADLALIVFSSTGRLYDFASSSMEAILERYQEAKEEHCGVLPTSEAKLWQREVTTLRRQVQNLHHNNR
QLLGEELSGTTVRDLQFLVNQVEMSLHSIRKRKEQVMAAEIHELNQKGLLVQKENIELDKKLSIAHEQNIEAMKSSEQQSSSKAAAGN-IDLELRQQEDE
>TaAGL17.5D1_TraesCS7D01G106900
MGRGKIAIERIDNTTNRQVTFSKRRGGLMKKARELAILCDADLALIVFSSTGRLYDFTSSSMEAILERYQEAKEEHSGVLPSSEAKLWQREVTTLRQQVQNLQHNNR
QLLGEELSGTNIRDLQFLVNQVEMSLHSVRKRKEQVMAAEIHGLNQKGFLIQKENIELGKKLSIAHEHNIESMVSSEQQSSSKAAAGN-IDLELRQQEDE
>TaAGL17.11D3_FGENESH
MARGKIVIRRIENMSSRKATFSKRRHGLLMKSRELAILCDVQVSVMVFSSTSRLYEYASSIMPSIIQNYQSAQEHHHLL-
SVSQVMLWEGEVRKLHQEIQMLQEHHGKFLIQFLP----------------------------------------------DIELRGESPDLKSGNGD---------
--------------------
>TaAGL17.11D2_TraesCS7D01G107000
MVRGKIVIRRIENMRSRQVTFSKRRRGLLKKARELAILCDVQVGVIIFSSTGRLYEYANSTM---------------------------------------------
--------------------------------------------------------------------------------------------QFA-----

>TaAGL17.5D2_TraesCS7D01G107200
MGRGKIVIERIDNPTNRQVTFSKRRGGLMKKARELAILCDADLALIVFSSTGRLYDFASSSMEAILERYQEAKEEHYGVLPTSEAKLWQREVTTLRQQVQNLQHNNR
QLLGEELSGTTARDLLFLVNQVETSLHSVRKRKEQLMAAEIHELNQKGFHIQKENVELGKKLSIAHEHKIEAMDSHVLN---------------------
>TaSEP1.4D_TraesCS7D01G120500
MGRGKVELKRIDNKISRQVTFAKRRNGLLKKAYELSVLCDAEVALIIFSTRGRLFEFSTSCMYKTLERYRSCNFNSEATA-P-
EIESNYQEYLKLKTRVEFLQTTQRNLLGEDLGPLNMKELEQLENQIEISLKHIRATKSQQSLDQLFDLKRKEQQLQDVNKDLRKKIQETTAQNVLDVGPSGSS-
QQEHF-H-DPSLRIGY----
>TaSEP1.5D_TraesCS7D01G120600
MRRGKVELKRIDNKSSRQVTFAKRRNGLLKKAHELSVLCDAEVALIIFSTRGRLFEFSTSCMYKTLERYRSCNFNSEATA-P-
EPESGYQEYLKLKQRVEFLQTTQRNLLGEDLGPLNMKELEQLENHVEISLKHIRATKSQQSFDQLLELKRKEQKLQDVNKDLRKKIQETSAESVLDDGPSGSS-
QQQHV-H-DPSLRMWY----
>TaAGL17.11D1_TraesCS7D01G135600LC
LVRSKIVIRRIENMSSRQVTSSKRRRSLLKKARELAILCDVQVGVIVFSSTGCLYEYANPPLPSF------------------------------------------
----------------------------------------------KTTNLHKSTISC------------------------------------------
>TaSVP.2D_TraesCS7D01G176700
MARERRAIRRIESAAARQVTFSKRRRGLFKKAEELAVLCDADVALVVFSSTGKLSQFASSSMNEIIDKYSTHSKNGKSDQPAIDLNLEHCKYDSLNEQLAEASLRLR
HMRGEELDGLSVGELQQMEKNLETGLQRVLCTKDRQFMQQISDLQQKGTQLAEENMRLKNQMHEVPTVST-DAHSSDSVVHSGS--SGDISLKLALP-WK
>TaSEP3.1D_TraesCS7D01G261600
MGRGRVELKRIENKINRQVTFAKRRNGLLKKAYELSVLCDAEVALIVFSNRGKLYEFCSQSMTKTLDKYQKCSYAGPETTQNEQLKNSRNEYLKLKARVDNLQRTQR
NLLGEDLDSLGIKELESLEKQLDSSLKHIRTTRTQHMVDQLTELQRREQMFSEANKCLRIKLEESNQVHG-HNNNVLGYGGNGFFHPAEPTLHIGY----
>TaFLC.1D1_TraesCS7D01G261900
RKRGRVELRRIEDRTSRQVRFSKRRSGLFKKAFELSVLCDAEVALVVFSPAGRLYEFVSTSVEQIFGRCKDIPDMVID-----
DLNIAARDSRGYCESVSELNSFAQMALELDVKEMSIAELTRFEEIVTEVLAAVKSIR---IMKVAE-LTQTEARLKRDPQK--------------------------
--------------W--
>TaAGL12.1D_TraesCS7D01G315900
MARGKVQLRRIENPVHRQVTFCKRRAGLLKKARELSVLCDADIGIIIFSAHGKLYDLATGTMDGLIERYKSASGEGMTGDGGDQRVDPKQEAMVLKQEIDLLQKGLR
YIYGNRAEHMNVDELNALERYLEIWMFNIRSAKMQIMIQEIQALKSKEGMLKAANEILQEKIVEQHDVGMT--DQ--QNTNPLTILSYCRGSEMGYS---
>TaFLC.1D2_TraesCS7D01G328100LC
RKRGRVELRRIEDRTSRQVRFSKRRSGLFKKAFELSVLCDAEVALVVFSPAGRLYEFVSSDTRY-------------------------------------------
-----------------------TVYPYVYYIL---I---------------------------------------------------------------
>TaAP3.1D_TraesCS7D01G380300
MGRGKIEIKRIENATNRQVTYSKRRSGIMKKARELTVLCDAQVAIIMFSSTGKYHEFCSTDIKGIFDRYQQAIGTS-----IEQYENMQRTLSHLKDINRNLRTEI-
-RMGEDLDALEFEELRDLEQNVDAALKEVRQRKYHVITTQTETYKKKVKHSQEAYKNLQQELGMREDPAYGDNP------------AMYAFRV-VPHGM
>TaBS.5D_TraesCS7D01G632400LC
MGRGRVEIKRIANDVSRRATFGKRSAGLLKKARELAVLCDVDLGVLVFDGAGRQLDYCSTSWSDLIERYESINHGE--------
YQKLLADIATLRHERDHLEASVRRQAGEDLPSATAAELRDLEHKLECALG---------------------------------------------------------
----------------
>TaAGL17.6U1_TraesCSU01G169600LC
MVRGKTVIEKIENKTSRQVTFSKRKSGLFKKGKELGILCDAQVGIFIFSNTGRLYEYSNS-MKPLIERYQAVKDGQKLL-
ASAEAKFWQAESARLEQQLRNLQENHRQLLGQHLSGLSFEDLKCLQNQLETSLYNIRLTKDKFIGDEIQELNTNESLMLQENIELHRKLNIVRQENTEQGVM----
FQKNPLPSKLPRIK-------
>TaAGL17.2A1_WM31A_TRIAE_CS42_5AS_TGACv1_394239_AA1279110
MGRGKIANERIDNATNRQVTFSKRRGGLMKKARELAILCDADLALIVFSSTGRLYDFASSRMEAILERYQEAKQEHCGVLPTSEAKLWQREVTTLRQQVHNLQHNNR
QLLGEELSGSTVRDLQFLVNQLETSLHSVRKRKEQVMAEEIHELNQKGFLIQKENIELGKKVRITHEQNIEAMRSSEQQSSSKAAAGRFIDLELRQQEDE
>TaAGL17.6U2_TraesCSU01G236400LC
MVRGKAVIEKIENQTSRQVTFSKRKSGLFKKGKELGVLCDAQVGIFIFSNTGRLYEYSNSGYVHI---------------AFSCSRIWIDNYSV------
NLIATYRYFFNSCIQSV------------------------------------------------------------------------------------------
>TaAGL17.6U3_TraesCSU01G236600LC
MVRGKAVIEKIENQTSRQVTFSKRKSGLFKKGKELGVLCDAQVGIFIFSNTGRLYEYSNS-MKPLIERYQAVKDGQKLL-
ASAEAKFWQAESARLEQQLHTLQENHR--------------------------YD----------------------------------------------------
--------------------
>TaAGL17.10U_TraesCSU01G389400LC
MARGKIVIRRIEKMSSRQVTFSKRRHGLMKKAHELAILCDVQVGVIVFSSTGHLYEYASSTMPSIIHKYQSAQENHQLL-
PVSQIMFWEGEVRKLQQEMQILQDHHRKLMGERLSNLGVNDLCLLENQLEKSLRCIREKKEQSFANYIVQLNEK---------------------------------
--------------------
